# IFN-γ-Induced Intestinal Epithelial Cell-Type-Specific Programmed Cell Death: PANoptosis and Its Modulation in Crohn’s Disease

**DOI:** 10.1101/2025.01.24.634679

**Authors:** Chansu Lee, Ji Eun Kim, Yeo-Eun Cha, Ji Hwan Moon, Eun Ran Kim, Dong Kyung Chang, Young-Ho Kim, Sung Noh Hong

## Abstract

Crohn’s disease (CD) is a chronic inflammatory bowel disease (IBD) and is considered a Th1-mediated disease, supported by the over-expression of interferon-gamma (IFN-γ) in the intestinal lamina propria. To evaluate the response of intestinal epithelial cells (IECs) to IFN-γ, we established small intestinal organoids (enteroids) derived from non-IBD controls and CD patients. IFN-γ stimulated programmed cell death (PCD) of IECs in both control and CD enteroids in a dose-dependent manner. Pyroptosis, apoptosis and necroptosis were activated in enteroids, suggesting that PANoptosis was the main process of IFN-γ-induced PCD in IECs. The response to IFN-γ depends on the cell type of the IECs. IFN-γ induced depletion of enterocytes with upregulation of PANoptosis-associated genes, while leading to expansion of goblet cells without significant change in PANoptosis-associated gene expression. Individual PCD inhibitors were insufficient to block IFN-γ-induced cytotoxicity, whereas the selective JAK1 inhibitor (upadacitinib) effectively blocked IFN-γ-induced cytotoxicity and PANoptosis. Furthermore, PANoptosis was significantly activated in surgically resected tissues and in publicly available single-cell RNA-sequencing datasets of intestinal tissues from patients with CD. In conclusion, IFN-γ induces PANoptosis in enterocytes, which can be treated with a selective JAK1 inhibitor in patients with CD.

## Introduction

Interferon-gamma (IFN-γ) is a key cytokine in cellular immunity, playing a critical role against intracellular pathogens and modulating inflammatory responses in host tissues (1). Its function, which has been extensively studied in immune cells, is essential in activating macrophages and natural killer cells to enhance pathogen clearance and in promoting Th1 cell differentiation (2). It also enhances antigen presentation by immune cells, thereby improving pathogen recognition capabilities (3, 4).

Crohn’s disease (CD) is a chronic inflammatory bowel disease (IBD) characterized by skipped bowel inflammation and damage that can affect any part of the gastrointestinal tract, particularly the small bowel (5). The prevalence of people diagnosed with CD has increased throughout the world over the past several decades, and the pooled incidence rate per 100,000 person-years has ranged from 4.1 to 10.7 in the United States (6, 7). In patients with CD, the production of IFN-γ by intestinal lamina propria and lymph node T cells is increased compared to patients with ulcerative colitis (UC) and healthy controls (8, 9), indicating that CD is classified as a Th1-mediated immune disease (10). However, the direct effects of IFN-γ on IECs, which are over-expressed in the lamina propria of patients with CD, are incompletely understood.

Although previous studies have shown that IFN-γ affects IECs by modulating cell proliferation and apoptosis (11), altering tight junction integrity (12, 13), and enhancing mucin and antimicrobial peptide secretion (14, 15), traditional experimental models using cell lines or animal studies have limitations in capturing the complex, cell-type-specific responses of diverse IECs. Intestinal epithelium-derived organoids are an advanced platform for culturing different types of epithelial cells (16) and recapitulating the crypt-villus structure of the intestine (17, 18). Patient-derived intestinal organoids provide a physiologically relevant framework for in-depth analysis of IEC behavior, facilitating the preservation of disease- and patient-specific pathologies (19).

Therefore, in this study, human small bowel organoids (enteroids) derived from healthy controls (Ctrl enteroids) and patients with CD (CD enteroids) were used to investigate what is the major response of IECs induced by IFN-γ to elucidate; which programmed cell death (PCD) pathways cause IFN-γ-induced cell death on IECs in the absence of intestinal microbiota and immune cells; what are epithelial cell type-specific responses to IFN-γ; and how these IFN-γ-induced responses of IECs differ in CD patient-derived enteroids. Next, we evaluated whether the identified responses could be verified in human samples from patients with CD and in publicly available data. Finally, we explored which PCD inhibitors could block IFN-γ-induced PCD in IECs.

## Method

### 1. Human enteroids model

A total of 22 human enteroid lines (10 enteroid lines derived from non-inflammatory bowel disease (IBD) controls [Ctrl enteroids] and 12 enteroid lines derived from patients with CD [CD enteroids]) were established as previously described (20, 21) and used in the experiments. The demographics, biopsy site, and clinical diagnosis of the enrolled patients are listed in Table 1. Small intestinal crypts were isolated from endoscopic biopsy specimens using 10 mM EDTA (ThermoScientific, San Jose, CA, USA) and 1 mM DTT (ThermoScientific) chelation. Isolated crypts were 3D cultured in Matrigel (Corning, New York, NY, USA) with maintenance medium [50% Wnt3a-conditioned medium (ATCC#CRL-2647, Manassas, VA, USA) + 50% 2 × basal medium : advanced DMEM/F12 (ThermoScientific) supplemented with antibiotic–antimycotic solution (ThermoScientific), 10 mmol/L HEPES (ThermoScientific), GlutaMAX (ThermoScientific), N2 (ThermoScientific), B27 (ThermoScientific), and 1 mmol/L N-acetylcysteine (Sigma–Aldrich, St. Louis, MO, USA) + 50 ng/mL recombinant human epidermal growth factor (Sigma– Aldrich), 100 ng/mL recombinant human noggin (R&D Systems, Minneapolis, MN, USA), 500 ng/mL recombinant human R-spondin-1 (PeproTech, Cranbury, NJ, USA), 10 μM Y27632 (Selleckchem, Houston, TX, USA) + 10 mM nicotinamide (Sigma–Aldrich), 10 μM SB202190 (Sigma–Aldrich), 10 nM Prostaglandin E2 (PGE2, Cayman Chemical, Ann Arbor, MI, USA). The medium was changed every 2 days and the organoids were passaged at a ratio of 1:2-1:4 after 7 to 9 days of culture. To recapitulate the structure and function of the intestinal epithelium, the intestinal organoids cultured for 2 days were grown in a differentiation medium (maintenance medium without Wnt3A conditional medium, SB202190, nicotinamide and PGE2) and described as enteroids. The differentiation medium was changed every 2 days and the enteroids cultured for 5 to 7 days were used for further experiments.

**Table 1.** Clinical Characteristics of Enrolled Subjects.

(A) Non-IBD patient-derived control enteroids
| ID | Demographics |  | Reason for performing endoscopy | Type of endoscopy | Sampling site of normal intestinal epithelium | Experiment |  |  |  |  |  |  |  |
| --- | --- | --- | --- | --- | --- | --- | --- | --- | --- | --- | --- | --- | --- |
|  | Age | Sex |  |  |  | ORA & MTT | EdU | WB | TUNEL | H&E | IHC | RNA-seq | scRNA-seq |
| N2 | 56 | M | to evaluate the ileal submucosal tumor (isolated polypoid primary lymphangiectasia) | SBE | mid ileum | p12 | p13 | p18 |  |  |  | p13 | p19 |
| #66 | 29 | M | to evaluate the jejunal submucosal tumor (ectopic pancreas) | SBE | proximal jejunum | p15 | p16 | p18 |  |  |  | p16 | p19 |
| #291 | 39 | M | to evaluate the small bowel bleeding (caused by Meckel's diverticulum) | SBE | distal ileum |  |  |  | p10 | p10 | p10 | p12 |  |
| #292 | 43 | M | to evaluate the small bowel bleeding (caused by Meckel's diverticulum) | SBE | distal ileum |  |  |  |  |  |  | p8 |  |
| #291 | 39 | M | to evaluate the small bowel obstruction after small bowel resection and anastomosis due to Meckel's diverticulum | SBE | distal ileum |  | p16 |  |  |  |  |  |  |
| #294 | 32 | M | to evaluate the cause of the small bowel obstruction (no mucosal lesions, bowel adhesion was causing the obstruction) | SBE | distal ileum | p13 | p15 |  | p18 | p18 | p18 | p16 |  |
| #295 | 62 | M | for endoscopic resection of early colon cancer | CFS | terminal ileum | p14 |  |  |  |  |  |  |  |
| #379 | 72 | M | for endoscopic resection of early colon cancer | CFS | terminal ileum | p8 |  |  | p11 | p11 | p11 |  |  |
| F#38 | 61 | M | for endoscopic resection of colonic benign tumor (tubular adenoma) | CFS | terminal ileum | p8 |  |  |  |  |  | p10 |  |
| F#203 | 58 | F | for endoscopic resection of colonic benign tumor (tubular adenoma) | CFS | terminal ileum |  |  | p8 |  |  |  |  |  |

(B) CD patient-derived control enteroids
| ID | Demographics |  | Montreal classification | Smoking | Surgery history | Medication during biopsy acquisition |  |  |  |  | Type of endoscopy | Sampling site | Inflammation at biopsy site | Experiment |  |  |  |  |  |  |  |
| --- | --- | --- | --- | --- | --- | --- | --- | --- | --- | --- | --- | --- | --- | --- | --- | --- | --- | --- | --- | --- | --- |
|  | Age | Sex |  |  |  | IMMs | IFX | ADA | UST | VDZ |  |  |  | ORA & MTT | EdU | WB | TUNEL | H&E | IHC | RNA-seq | scRNA-seq |
| #96 | 29 | M | A2L1+4B2p | non-smoker | none | Aza | — | — | — | — | SBE | mid jejunum | non-inflamed | p16 | p18 | p18 |  |  |  | p20 | p22 |
| #97 | 30 | M | A2L3B2 | non-smoker | none | Aza | — | — | — | — | SBE | distal ileum | non-inflamed | p18 |  |  |  |  |  |  |  |
| #123 | 49 | M | A2L3B2p | ex-smoker | Right hemicolectomy | MTX | Y | F | — | — | CFS | terminal ileum | inflamed | p8 | p10 |  |  |  |  | p12 |  |
| #293 | 27 | M | A2L3B2p | non-smoker | none | — | — | — | — | — | SBE | distal ileum | inflamed | p14 |  |  | p10 | p10 | p10 | p12 |  |
| #296 | 34 | M | A2L1B2 | ex-smoker | SB R&A | intolerance | F | — | Y | — | SBE | distal ileum | non-inflamed | p9 | p11 |  | p13 | p13 | p13 |  | p15 |
| #375 | 43 | F | A3L3B3p | non-smoker | SB R&A | Aza | — | F | Y | — | SBE | distal ileum | non-inflamed | p12 |  |  |  |  |  | p10 |  |
| #376 | 34 | M | A2L1B2 | ex-smoker | SB R&A, ileocectomy | Aza | F | Y | F | — | SBE | distal ileum | non-inflamed | p11 |  |  |  |  |  | p13 |  |
| #389 | 48 | F | A3L3B1p | non-smoker | none | Aza | Y | — | — | — | SBE | terminal ileum | inflamed | p8 |  |  |  |  |  | p10 |  |
| #391 | 31 | M | A2L1B2p | non-smoker | none | 6MP | — | — | — | — | SBE | distal ileum | non-inflamed | p15 |  |  | p12 | p12 | p12 |  |  |
| #405 | 47 | F | A2L1B2 | non-smoker | SB R&A | — | — | — | — | — | SBE | distal ileum | inflamed | p10 | p14 |  |  |  |  |  |  |
| F#18 | 31 | M | A2L3B2 | non-smoker | SB R&A | intolerance | F | F | Y | F | SBE | distal ileum | inflamed | p12 | p6 |  |  |  |  | p9 |  |
| F#35 | 42 | M | A2L3B3p | ex-smoker | SB R&A | intolerance | — | — | Y | — | SBE | mid ileum | inflamed | p9 |  |  |  |  |  | p6 |  |
| F#63 | 37 | M | A2L1B3p, | ex-smoker | SB R&A | Aza | — | F | Y | — | SBE | terminal ileum | inflamed | p9 |  |  |  |  |  | p6 |  |
| F#303 | 33 | F | A2L3+4B3p | non-smoker | SB R&A, ileocectomy | MTX | F | F | Y | F | SBE | distal ileum | inflamed | p8 |  | p6 |  |  |  |  |  |
| F#307 | 28 | F | A2L1B3p | non-smoker | none | Aza | Y | — | — | — | SBE | distal ileum | non-inflamed | p9 |  | p6 |  |  |  |  |  |

Abbreviation: IBD, inflammatory bowel disease; CD, Crohn's disease; CFS, colonoscopy; SBE, single-balloon enteroscopy; ORA, organoid reconstitution assay; MTT, 3-(4,5-dimethylthiazol-2-yl)-2,5-diphenyltetrazolium bromide assay; Edu, 5-ethynyl-2'-deoxyuridine assay; WB, Western blot; H&E, Hematoxylin and Eosin staining; IHC, immunohistochemistry; RNA-seq, bulk RNA sequencing; scRNA-seq, single-cell RNA sequencing; p, passage; SB R&A, small bowel resection and anastomosis; IMM, immunomodulator; Aza, azathioprine; MTX, methotrexate; IFX, infliximab; ADL, adalimumab; UST, ustekinumab; VDZ, vedolizumab; Y, in use; F, failure.

### 2. Organoid reconstitution assay

To evaluate the IFN-γ-induced responses in the enteroids, the intestinal organoids were cultured in a maintenance medium for 2 days to obtain a stable number of organoids. The culture medium was then replaced every 2 days with differentiation medium containing different concentrations of human recombinant IFN-γ (R&D Systems). The number of enteroids was counted after 7 days of embedding the mechanically and chemically disrupted organoid under an inverted microscope. Organoid-forming efficiency was expressed as the percentage of the number of IFN-γ-treated enteroids after 7 days of culture relative to the number of IFN-γ-free enteroids after 7 days of culture. The 50% cell growth inhibition concentration (GI50) was calculated as the concentration of IFN-γ required for 50% inhibition of maximal organoid-forming efficiency of human enteroids. Assays were performed in triplicate with 6 Ctrl enteroids and 15 CD enteroids.

### 3. MTT assay

Ten microliter of MTT (Sigma-Aldrich) was added to each well of the 96-well culture plates containing enteroids and incubated for 3 hours until purple precipitates were visible. The optical density (OD) of the formazan precipitate per well was measured and expressed as the ratio of the OD of IFN-γ-treated enteroids after 7 days of culture to the OD of IFN-γ-free enteroids after 7 days of culture. GI50 was calculated as the concentration of IFN-γ required for 50% inhibition of maximal OD value of human enteroids. Assays were performed in triplicate with 6 Ctrl enteroids and 7 CD enteroids.

### 4. EdU assay

Two hundred micrograms of EdU (Abcam, Cambridge, UK) was added 2 h prior to fixation with cold 4% paraformaldehyde (Biosesang, Seongnam, Korea). Incorporation of EdU into DNA was determined using the Click-iT™ EdU Alexa Fluor® 488 Imaging Kit (ThermoScientific). The number of EdU-positive cells was measured in 5 randomly selected enteroids (size > 50 μm) from independent 3 Ctrl enteroids and 5 CD enteroids.

### 5. Western blot analysis

Proteins were isolated from 2 human enteroids via lysis and homogenized using Protein Extraction Reagent (ThermoScientific). Prepared protein samples were separated using Precast Protein Gels (Bio-Rad, Hercules, CA, USA) for electrophoresis, and electrotransferred onto polyvinylidene difluoride membranes (Millipore, MA, USA). The blots on membranes were probed with specific primary antibodies, caspase-3 (CASP3, Cell signalling #9662, full length: 35 kDa, cleaved form [c-CASP3): 19 kDa and 17 kDa), RIP3 (RIPK3, Cell signalling #95702, 46 kDa), phospho-RIP3 (p-RIPK3, Cell signalling #93654, 46-62 kDa), MLKL (Cell signalling #14993, 54 kDa), phospho-MLKL (p-MLKL, Cell signalling #91689 54 kDa), GSDMD (Cell signalling #97558, 53 kDa), cleaved GSDMD (c-GSDMD, Cell signalling #36425, 30 kDa), and GAPDH (Abcam, ab8245) overnight at −4°C in a cold room. Subsequently, immunoreactive proteins were detected with HRP-conjugated secondary antibody (Invitrogen) and enhanced chemiluminescence reagents (ThermoScientific). Western blot was performed on the independent 3 Ctrl enteroids and 3 CD enteroids.

### 6. TUNEL Assay

Apoptosis-associated DNA fragmentation was detected by TUNEL assay using the In Situ Cell Death Detection Kit (Merck, Darmstadt, Germany). Positive control sections were incubated with 10 U/mL recombinant DNase I, and negative control sections were processed in the same manner without the terminal transferase enzyme. The number of TUNEL-positive cells was measured in 10 randomly selected enteroids (size > 50 μm) from independent 3 Ctrl and CD enteroids.

### 7. Hematoxylin and Eosin (H&E) staining and Immunohistochemistry (IHC)

Human enteroids were fixed in 4% paraformaldehyde for 30 minutes at room temperature and double embedded in HistoGel (ThermoScientific) and paraffin. Histologic evaluation was performed with H&E staining. The area occupied by 10 randomly selected enteroids (size >50 μm) from each of the IFN-γ-free and IFN-γ-treated Ctrl enteroids (n = 3) and CD enteroids (n = 3) was measured using ImageJ.

After heat-induced epitope retrieval with citrate buffer, immunohistochemical staining was performed against OLFM4 (1:200, Invitrogen/PA5-32954, Carlsbad, CA, USA), proliferating cell nuclear antigen (PCNA, 1:6000, Abcam/ab29), and lysozyme (LYZ, 1:200, Abcam/ab108508). OLFM4-stained cells were calculated in randomly selected 5 enteroids (size >50 μm) from each of the IFN-γ-free and IFN-γ-treated Ctrl enteroids (n = 3) and CD enteroids (n = 3).

Using serial sections of surgical specimens obtained from patients with Crohn’s disease (n = 5), IHC was performed with antibodies against of c-CASP3 (1:500, Cell Signaling #9661), p-MLKL (1:1,000, R&D Systems #MAB9187), and c-GSDMD (1:1,000, Cell Signaling #36425) using serial sections of surgical specimens obtained from patients with Crohn’s disease (n = 5).

### 8. Alcian blue assay

After heat-induced epitope retrieval with citrate buffer, paraffin-embedded organoids were fixed with 3% acetic acid and stained with Alcian blue solution (Sigma-Aldrich) for 30 minutes and washed with distilled water. The Alcian blue-stained area was quantified using ImageJ and expressed as a percentage of the total enteroid area. The Alcian blue-stained area was measured in randomly selected 5 enteroids (size >50 μm) from each of IFN-γ-free and IFN-γ-treated Ctrl enteroids (n = 3) and CD enteroids (n = 3).

### 9. RNA sequencing (RNA-seq)

Total RNA was extracted from each of IFN-γ-free and IFN-γ-treated Ctrl enteroids (n = 6 pairs) and CD enteroids (n = 10 pairs) using the RNeasy Mini Kit (QIAGEN, Hilden, Germany). RNA-seq was performed with >10 μg RNA and an RNA integrity number >8. cDNA libraries were sequenced on HiSeq2500 in 100 bp paired-end mode. Reads from files in FASTQ format were mapped to the hg19 human reference genome. The batch effect was adjusted using the ComBat function in the sva R package (version 3.48.0) (22). DEG analysis was performed using the edgeR R package (version 3.42.4) (23). Sequence read counts were normalized to fragments per kilobase of exon per million mapped fragments (FPKM). Differences in FPKM between IFN-γ-free and IFN-γ-treated enteroids were evaluated by paired *t*-test with Bonferroni correction. Unsupervised hierarchical clustering analysis using the Euclidean distance and complete linkage algorithm was used to generate a heat map with associated dendrogram.

### 10. Single-cell RNA sequencing (scRNA-Seq)

IFN-γ-free and IFN-γ-treated enteroids were mechanically disrupted by vigorous pipetting and enzymatically digested into single-cell suspension using TrypLE Express (ThermoScientific). Single-cell suspensions isolated from each of IFN-γ-free and IFN-γ-treated Ctrl enteroids (n = 2 pairs) and CD enteroids (n = 2 pairs) were combined. Bar-coded sequencing libraries were prepared and sequenced on a HiSeq X Ten system, targeting 15,000 er sample. Reads were aligned to a GRCh38 human reference genome and processed using the CellRanger 7.1.0 pipeline (10X Genomics). The raw gene expression matrix was filtered using Seurat R package (version 5.1.1) and selected according to the following criteria: 5000 ≥ nFeature_RNA > 200 & percent.mt ≤ 15). Cells that passed the filtering criteria were integrated using Harmony R package (version 1.2.1). Epithelial celltype was annotated based on the differentially expressed epithelial lineage specific genes found in the cluster. The epithelial lineage specific genes were based on the published single-cell RNA sequencing data (https://panglaodb.se/) (24). When several cell markers were expressed significantly in a single cluster, the cell type was assigned based on the number of cell markers expressed or established.

Single-cell analysis of human intestinal cells was performed using a publicly available single-cell RNA-seq dataset containing dataset of IECs originated from healthy adult and pediatric CD (https://www.gutcellatlas.org/) (25) using the same pipeline used for scRNA-seq of IFN-γ-free and IFN-γ-treated human enteroids.

### 11. Trajectories Analysis

Slingshot was used to illustrate lineage differentiation within all clusters (26). The start point was set at cluster 2 (intestinal stem cell [ISC]). The endpoint was inferred by pseudotime ordering along the trajectory of the differentiated cell-enriched clusters.

### 12. Identification of Protein-Protein Interaction (PPI) Networks

The construction and visualization of the PPI network based on DEGs were performed using the online STRING database (http://string-db.org/) (27). The combined score > 0.4 was set as the threshold value. Further subnetworks were clustered by the MCL inflation parameter (set to 3).

## Results

### Effect of IFN-γ on Organoid Formation, Viability and Proliferation in Ctrl and CD Enteroids

To evaluate the IFN-γ-induced response of human IECs, Ctrl and CD enteroids were cultured with a range of 0 to 100 ng/mL IFN-γ (Figure 1A). As the concentration of IFN-γ in the culture media increased, disruption of organoid structure, growth arrest, and cell death were observed in IFN-γ-treated Ctrl and CD enteroids. There were no significant differences in IFN-γ-induced organoid-forming efficiency and cell viability between Ctrl and CD enteroids (*p*=0.502 and *p*=0.514, respectively; Figure 1B-C). Depending on the concentration of IFN-γ, the organoid-forming efficiency of Ctrl and CD enteroids was significantly reduced. Based on the organoid reconstitution assay, the GI50 values of Ctrl and CD enteroids were estimated to be 331.6 pg/mL and 269.1 pg/mL, respectively (Figure 1B). Organoid cell viability was significantly decreased in response to IFN-γ. Based on the MTT assay, the GI50 values of Ctrl and CD enteroids were 78.9 pg/mL and 311.5 pg/mL, respectively (Figure 1C). The changes in organoid formation and cell viability were dynamic between 100 pg/mL and 1000 pg/mL IFN-γ. In this study, the IFN-γ concentration of 100 pg/mL in the culture medium was chosen as the experimental dose to evaluate the IFN-γ-induced molecular and cellular response on human enteroids.

**Figure 1.**
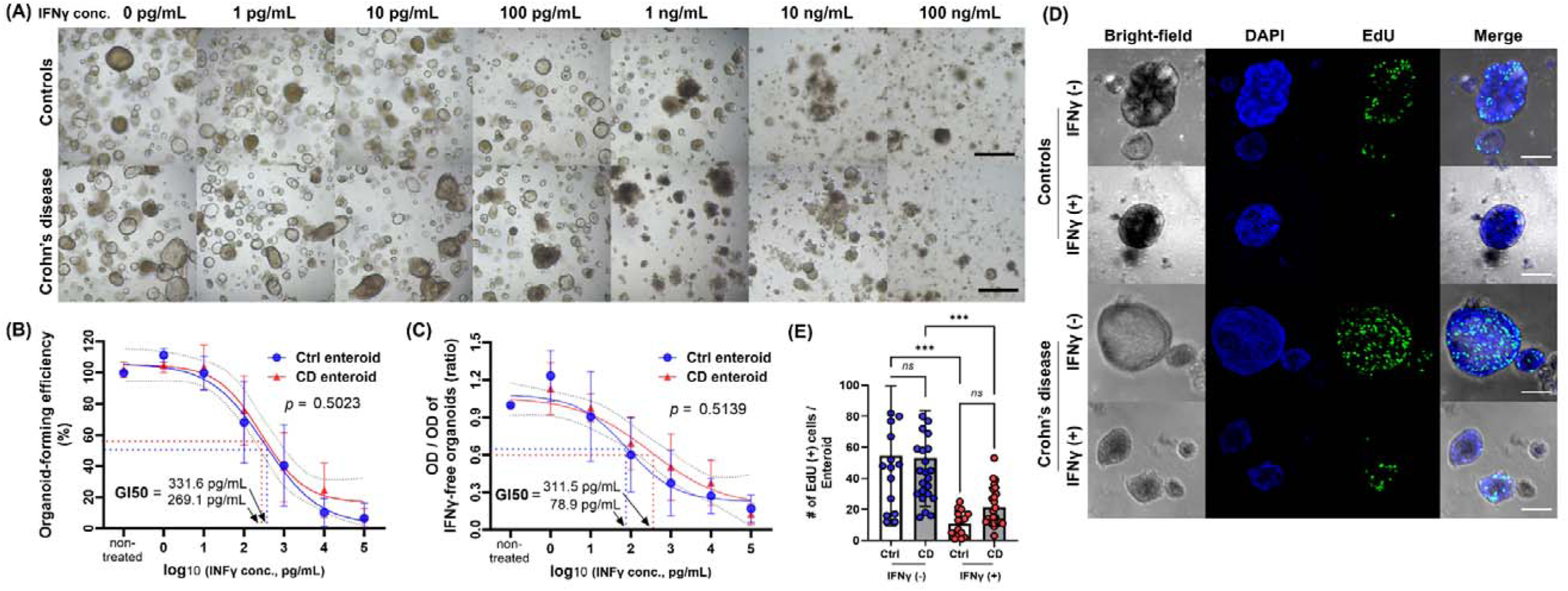
Effect of IFN-γ on Ctrl and CD enteroids. (A) Bright field microscopic image of different concentrations of IFN-γ-treated Ctrl and CD enteroids. Scale bar, 100 μm. (B) Organoid-forming efficiency as measure by organoid reconstitution assay. Assays were performed in triplicate with seven Ctrl enteroids and five CD enteroids. (C) Cell viability as measured by MTT assay. Assays were performed in triplicate with seven Ctrl enteroids and five CD enteroids. GI50, 50% growth inhibition concentration. (D) Fluorescence microscopy image of the EdU assay. Scale bar, 25 μm. (E) Comparison of EdU-positive cells per enteroids. Assays were performed on three Ctrl enteroids and five CD enteroids. Differences were evaluated by one-way ANOVA with multiple comparisons; ** *p* < 0.01, *** *p* < 0.001

As the concentration of IFN-γ in the culture media increased, the number of EdU-positive proliferating cells in IFN-γ-treated Ctrl enteroids was decreased in IFN-γ-treated Ctrl enteroids (Figure 1D. Supplementary Material S1). Two hours after EdU injection, the average number of EdU-positive proliferating cells in IFN-γ-treated enteroids was significantly lower than in 100 pg/mL IFN-γ-untreated enteroids (Ctrl enteroids: *p*<0.001; CD enteroids: *p*=0.001, Figure 1E). The number of EdU-positive proliferating cells in IFN-γ-treated CD enteroids was numerically high, but did not reach a statistically significant difference (10.9 ± 8.0 vs. 21.5 ± 12.4; *p*=0.634).

These data suggest that a high concentration of IFN-γ induces apparent cytotoxicity in human enteroids. Based on previous studies, possible mechanisms of IFN-γ-induced cytotoxicity may result from induction of PCD pathways (28–30).

### IFN-γ-induced programmed cell death (PCD) in Ctrl and CD enteroids

The main types of PCD include apoptosis, necroptosis, and pyroptosis. PANoptosis is a newly defined inflammatory PCD pathway that is uniquely regulated by multifaceted PANoptosome complexes and highlights significant crosstalk and coordination among pyroptosis, apoptosis, and/or necroptosis (31). To determine which type of PCD, including apoptosis, necroptosis, pyroptosis, and PANoptosis, is induced in IECs of human enteroids by IFN-γ, the gene expression profiles of these PCDs were analyzed using bulk RNA-seq data performed on the paired samples of IFN-γ-free and IFN-γ-treated Ctrl and CD enteroids. Principal component analysis (PCA) revealed a clear separation of the gene expression profiles of enteroids depending on IFN-γ treatment (Figure 2A). However, no clear separation was observed between the Ctrl and CD enteroids. The heatmap showed that the expression of apoptosis-, necroptosis-, pyroptosis-, and PANoptosis-associated genes was up-regulated in the IFN-γ-treated human enteroids (Figure 2B, Supplementary Material S2). Paired comparison plot demonstrated the gene expression of key molecules involved in apoptosis (BAK1, CASP3), necroptosis (RIP, MLKL), pyroptosis (CASP1, GSDMD), and PANoptosis (ZBP1, CASP8) was significantly increased in IFN-γ-treated all Ctrl and CD enteroids (*p*<0.05 for all, Figure 2C). There were no significant differences in the expression of these key molecules between Ctrl and CD enteroids, except for CASP1. CASP1 was higher in IFN-γ-treated CD enteroids, but the IFN-γ-induced increasing pattern was similar to that of IFN-γ-treated Ctrl enteroids (*p*<0.0001).

**Figure 2.**
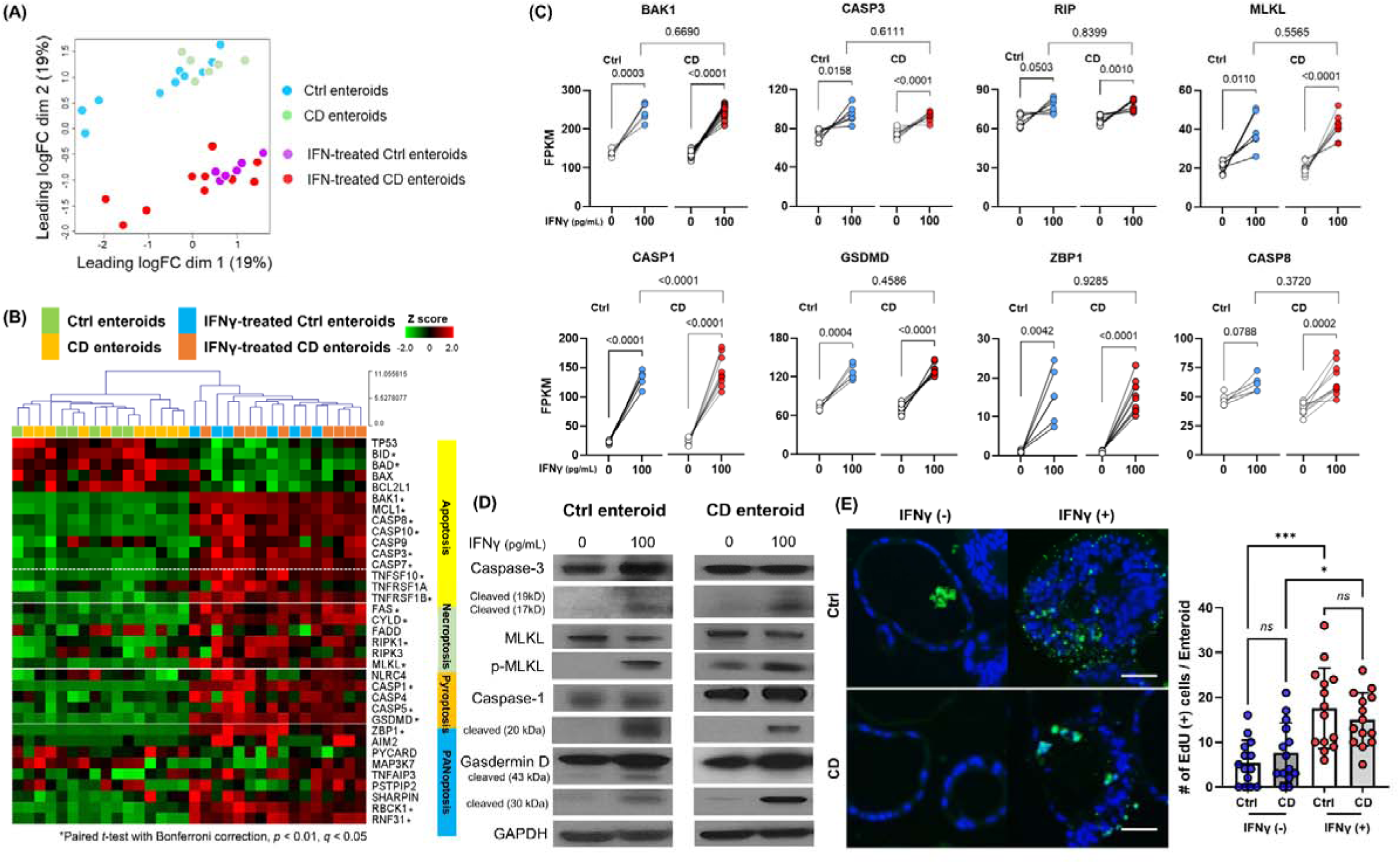
Programmed cell death (PCD) pathway in IFN-γ-treated Ctrl and CD enteroids. (A) Principal component analysis (PCA) plot of bulk RNA-seq data set (IFN-γ-free and IFN-γ-treated Ctrl enteroids, n = 6 pair; IFN-γ-free and IFN-γ-treated CD enteroids, n = 10 pair). (B) Heatmap with hierarchical clustering of apoptosis, necroptosis, pyroptosis, and PANoptosis -associated genes in human enteroids treated with and without IFN-γ 100 pg/mL. Differences were evaluated by paired *t*-test with Bonferroni correction; \**p*-value < 0.01, adjusted *p*-value < 0.01, and *q*-value < 0.05. (C) Gene expression level [fragments per kilobase of transcript per million mapped reads (FPKM)] for the key molecules of PCD pathways in IFN-γ-free and IFN-γ-treated Ctrl and CD enteroids. Paired *t*-test were used for comparison of the paired IFN-γ-free and IFN-γ-treated enteroids. Unpaired *t*-test were used for comparison of the IFN-γ-treated Ctrl and CD enteroids. (D) Western blotting for the key molecules of PCD pathways in IFN-γ-free and IFN-γ-treated Ctrl enteroids (n = 4) and CD enteroids (n = 3). (E) TUNEL staining in IFN-γ-free and IFN-γ-treated Ctrl enteroids (n = 3) and CD enteroids (n = 3). Scale bar, 25 μm. Differences of TUNEL-positive cells per orgnaoids in the IFN-γ-free and IFN-γ-treated Ctrl and CD enteroids were evaluated by one-way ANOVA with multiple comparisons; ** *p* < 0.01.

The activation of apoptosis, necroptosis and pyroptosis should be accurately identified by Western blot analysis, which demonstrated the upregulation of cleaved caspase-3, phosphorylated MLKL and cleaved caspase-1, and cleaved GSDMD in IFN-γ-treated Ctrl and CD enteroids (Figure 2D). In addition, TUNEL-positive cells per enteroid were higher in IFN-γ-treated enteroids compared to IFN-γ-free enteroids (Figure 2E). Although DNA fragmentation detected by TUNEL is used as a marker of apoptosis, any type of PCD also produces DNA fragments that react with TUNEL (20, 32, 33). Positive TUNEL staining indicates that cell death by PCD has occurred in IFN-γ-treated enteroids.

PANoptosis is a unique form of inflammatory PCD characterized by the coordinated activation of pyroptosis, apoptosis, and necroptosis. Although there is no single, definitive master regulatory marker for PANoptosis, the expression and activation of certain genes, such as ZBP1, CASP8, and CASP1, have been implicated in its regulation and activation (34). Taken together, our results indicate that IFN-γ induced PCD in human IECs through the PANoptosis in both Ctrl and CD enteroids. In pathway analysis, DEG was over-expressed in the programmed cell death and apoptosis pathways in REACTOME, WikiPathways, and PathBank databases. Gene set enrichment analysis (GSEA) identified activation of Regulation of Apoptosis, Cell Killing pathway as well as IFN-γ signaling pathway (Supplementary Material S3). REACTOME network plot revealed the central role of programmed cell death as a key regulatory pathway interacting with a wide range of genes and biological processes (Supplementary Material S4). The activation of programmed cell death pathway and cell killing pathway in IFN-γ-tre4ted enteroids supports the activation of PANoptosis in pathway analysis and GSEA.

### IFN-γ-induced altered epithelial cell differentiation in Ctrl and CD enteroids

IFN-γ can affect cell fate determination and differentiation (35), resulting in altered or inhibited differentiation of specific cell types within the enteroids. To evaluate IEC differentiation and cell-type-specific PCD-associated gene expression in response to IFN-γ, scRNA-seq was performed on IFN-γ-free and IFN-γ-treated enteroids (Supplementary Material S5). After excluding poor quality cells, a total of 11,316 cells were analyzed. To characterize the cellular subpopulation, the analyzed cells were divided into 17 clusters (0 to 16) based on the presence of different sets of co-expressed genes (Supplementary Material S6).

Cell clusters were assigned to epithelial cell lineages based on epithelial lineage-specific marker expression (Figure 3A-B, Supplementary Material S7). In cluster 2, the expression of the ISC marker OLFM4 was significantly higher than in the other clusters (average log2 fold change [logFC]=1.80, *p*<0.0001). The proliferating cell marker MKI67 was expressed exclusively in cluster 10 (logFC=2.00, *p*<0.0001). Cells in clusters 8, 11, and 12 expressed multiple enterocyte (EC) markers and were annotated as ECs, and several Paneth cell (PC) markers were significantly highly expressed in cluster 5. In cluster 4 and 7, the goblet cell (GC) markers, MUC4 and MUC 13, were significantly upregulated compared to the remaining clusters (*p*< 0.0001 for both). On the other hand, clusters 0, 1, 3, 6, 9, 13, 14, 15 and 16 could not be annotated with a specific epithelial cell type or were annotated with a rare cell type and were not included in the analysis of IEC differentiation and intestinal epithelial cell type-specific response. Clusters 0, 1, 3, and 9 cells were in stress-adaptive phase as indicated by metabolic and mitochondrial dysfunction, and clusters 6, 14, and 16 cells consisted of damaged or pre-damaged cells, as indicated by increased immune signaling and potential apoptotic processes. Cluster 13 cells did not express the exclusive epithelial lineage-specific markers, so these clusters could not be classified as a specific epithelial cell type and were not included in the analysis. Cluster 15 significantly expressed microfold (M) cell markers; however, M cells can be considered a rare cell type, estimated at <1% of total IECs.

**Figure 3.**
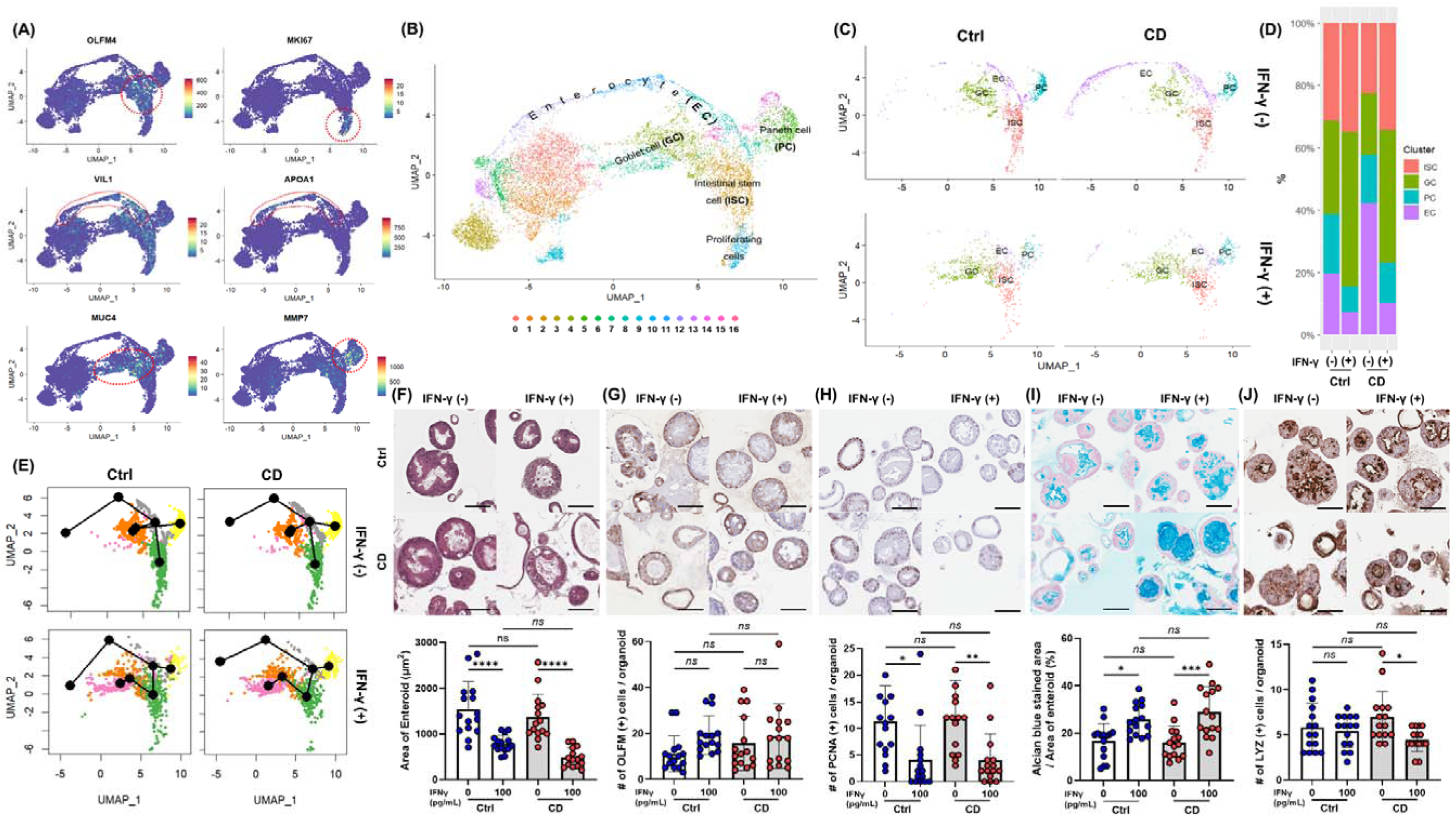
IFN-γ-induced epithelial cell differentiation in Ctrl and CD enteroids. (A) Expression of intestinal epithelial lineage-specific markers. Single-cell RNA-sequencing analyzed 11,316 cells derived from IFN-γ-free and IFN-γ-treated Ctrl enteroids (n = 2 pair) and IFN-γ-free and IFN-γ-treated CD enteroids (n = 2 pair). To identify different intestinal epithelial cell types, lineage-specific markers were applied: OLFM4 for intestinal stem cells (ISCs), MKI67 for proliferating cells, VIL1 for enterocytes (ECs), MMP7 for Paneth cells (PCs), and MUC4 and TFF3 for goblet cells (GCs). (B) FeaturePlot annotation based on epithelial cell type. (C) Effect of IFN-γ on epithelial cell types in Ctrl and CD enteroids. (D) Number and percentage of epithelial cell types occupying Ctrl and CD enteroids treated with IFN-γ. (E) Trajectory analysis of IFN-γ treated Ctrl and CD enteroids using Slingshot. (F) Hematoxylin and eosin (H&E) staining of Ctrl and CD enteroids treated with and without IFN-γ to measure the area of enteroids. H&E staining, immunohistochemistry, and alcian blue assay were performed on the IFN-γ-free and IFN-γ-treated Ctrl enteroids (n = 3 pair) and IFN-γ-free and IFN-γ-treated CD enteroids (n = 3 pair). (G) IHC for OLFM4 to compare OLFM4-stained intestinal stem cells in Ctrl and CD enteroids treated with and without IFN-γ. (H) IHC for PCNA to compare PCNA-stained proliferating cells in Ctrl and CD enteroids treated with and without IFN-γ. (I) Alcian blue staining to compare the percentage of Alcian blue-stained area to the total area of enteroids in Ctrl and CD enteroids treated with and without IFN-γ. (J) IHC for lysozyme to compare lysozyme-stained Paneth cells in Ctrl and CD enteroids treated with and without IFN-γ. Scale bar 100 μm. Differences were evaluated by one-way ANOVA with multiple comparisons; * *p* < 0.05, ** *p* < 0.01, **** *p* < 0.0001

IFN-γ treatment altered the differentiation of specific cell types within the enteroids (Figure 3C-D, Supplementary Material S8). In IFN-γ-treated Ctrl and CD enteroids, the proportion of ISC was significantly increased (Ctrl enteroids: 487/4,554 vs. 318/1,559, *p*<0.0001; CD enteroids: 322/2,900 vs. 307/2,303 *p*=0.0147), while the proportion of proliferating cells was decreased compared to IFN-γ-free enteroids (ctrl enteroids: 227/4,554 vs. 14/1,559, *p*<0.0001, CD enteroids: 98/2,900 vs. 14/2,303 *p*<0.0001). The proportion of ECs was significantly reduced (Ctrl enteroids: 302/4,554 vs. 66/1,559, *p*=0.0004, CD enteroids: 602/2,900 vs. 90/2,303 *p*<0.0001), while the proportion of GCs increased (Ctrl enteroids: 461/4,554 vs. 452/1,559, *p*<0.0001, CD enteroids: 287/2,900 vs. 381/2,303, *p*<0.0001). Unlike other secretory lineage cell types, the proportion of PCs was reduced in IFN-γ-treated enteroids (Ctrl enteroids: 296/4,554 vs. 74/1,559, *p*=0.0116, CD enteroids: 220/2,900 vs. 117/2,303 *p*=0.0003). Slingshot analysis indicated no significant difference in differentiation trajectories between Ctrl and CD enteroids, suggesting that cells from both groups follow similar developmental paths (Figure 3E).

H&E staining of IFN-γ-treated enteroids showed a slightly blurred basal surface on the cell membrane and increased vesicle formation in the cytoplasm (Figure 3F). ECs constitute the majority of the enteroids. The significant reduction in size of IFN-γ-treated enteroids suggests a decrease in ECs, indicating loss and impaired differentiation. This effect was evident in both Ctrl and CD enteroids, as shown by the notable decrease in overall enteroid size upon IFN-γ treatment. (Ctrl enteroid, *p*<0.0001; CD enteroid, *p*<0.0001, Figure 3F). IHC with OLFM4 showed that the number of cytoplasmic OLFM4-stained cells per enteroid was numerically higher in IFN-γ-treated Ctrl and CD enteroids compared to IFN-γ-free Ctrl and CD enteroids, but not statistically significant (Ctrl enteroid, *p*=0.1821; CD enteroid, *p*=0.8509, Figure 3G). The number of PCNA-positive proliferating cells per enteroid was significantly higher in IFN-γ-free Ctrl and CD enteroids compared to IFN-γ-treated Ctrl and CD enteroids (Ctrl enteroid, *p*=0.0150; CD enteroid, *p*=0.0051, Figure 3H). Alcian blue staining revealed that the proportion of Alcian blue-stained area per total enteroid area was significantly increased in IFN-γ-treated Ctrl and CD enteroids compared with IFN-γ-free Ctrl and CD enteroids (Ctrl enteroid, *p*=0.0189; CD enteroid, *p*=0.0003, Figure 3I). The number of LYZ-positive PC per enteroid was significantly higher in IFN-γ-free enteroids compared to IFN-γ-treated enteroids in CD enteroids (*p*=0.0200, Figure 3J), but not in Ctrl enteroids (*p*=0.9616). There was no significant difference in the number of each type of IEC between the Ctrl and CD enteroids.

### IFN-γ-induced epithelial cell type-specific expression of PCD-associated genes

The expression of PANoptosis-related genes identified in previous studies (36–38) was upregulated in IFN-γ-treated Ctrl and CD enteroid cells, except in goblet cell populations (Figure 4A). Interestingly, in GCs derived from IFN-γ-treated enteroids, the expression of PCD-related genes was not up-regulated compared with IFN-γ-free enteroids (Figure 4B).

**Figure 4.**
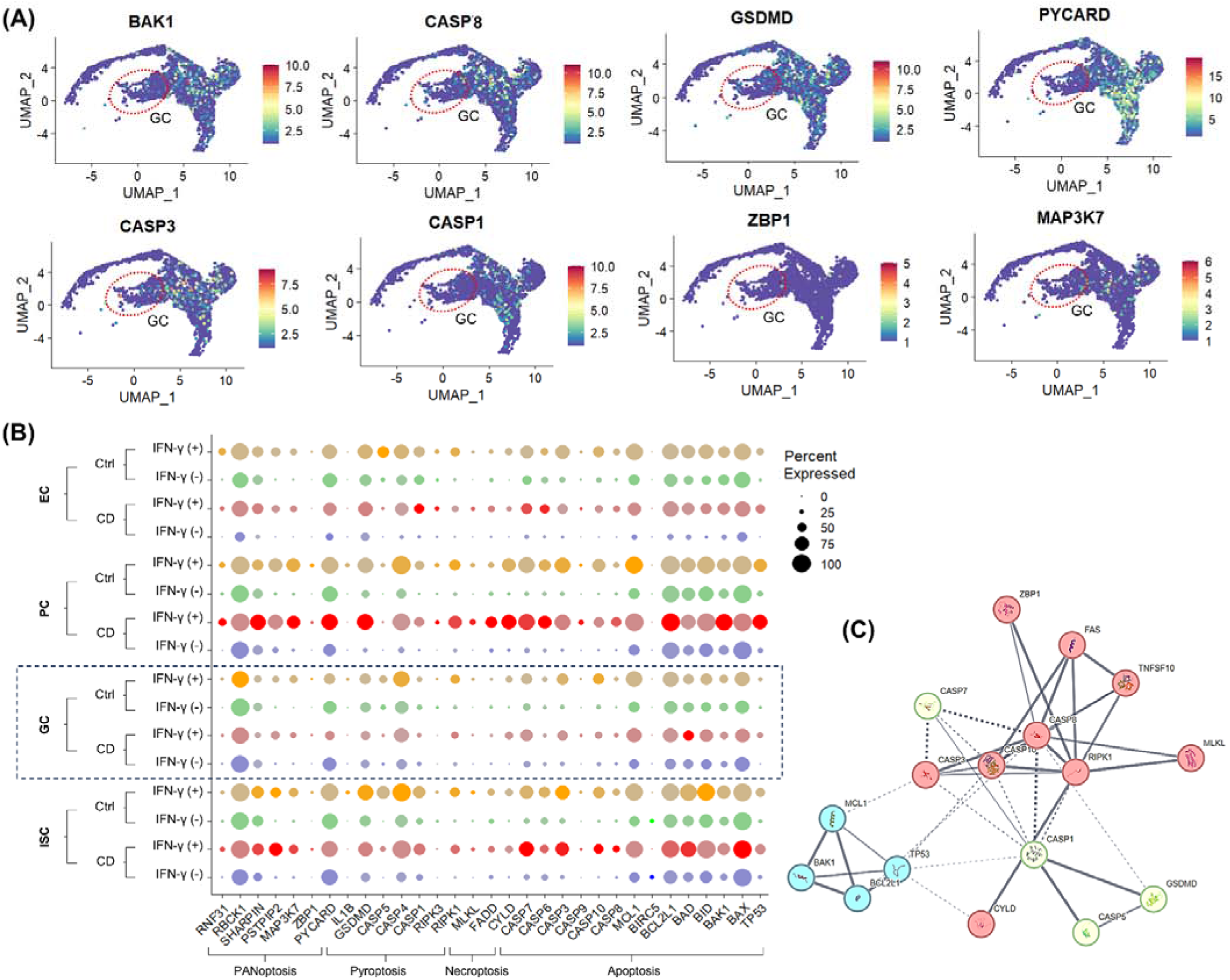
IFN-γ-induced epithelial celltype-specific responses of PCD-associated genes (A) FeaturePlot for PCD-associated gene expression. The red dashed circle indicates the goblet cell (GC) population. (B) DotPlot for PCD-associated gene expression in Ctrl and CD treated with and without IFN-γ. (C) Protein-protein interaction network of DEGs in PCD pathways.

The protein-protein interaction network was constructed using 13 DEGs among PCD-associated genes, and the 17 nodes and 41 edges were identified (Figure 4C). To screen the hub genes, the critical subnetworks were extracted by MCL clustering, and 3 significant modules were found. The most significant module 1 consisted of 10 nodes and 21 edges. The top cellular component of GO was Death-inducing signaling complex (false discovery rate [FDR]=4.16e-11; GO:0031264). Module 2 consisted of 4 nodes and 6 edges, of which the top cellular GO was Bcl-2 family protein complex (FDR=1.84e-06; GO:0097136). Module 3 consisted of 3 nodes and 3 edges, of which the top cellular component of GO was inflammasome complex (FDR=1.56e-06; GO:0061702). Module 1 and Module 3 were associated with RIPK1.

Single-cell RNA-seq of human enteroids revealed that IFN-γ altered epithelial cell differentiation in Ctrl and CD enteroids, including expansion of the GC population and depletion of the EC population, along with upregulation of PANoptosis-associated genes. Next, we thought that the IFN-γ-induced responses of IECs identified in the enteroid model should be validated in human tissue.

### Validation of PANoptosis and Epithelial Cellular Responses in Human CD Samples

In surgically resected intestinal specimens from patients with CD, IHC revealed that the expression of apoptosis marker c-CASP3, necroptosis marker p-MLKL, and pyroptosis marker c-GSDMD was expressed in IECs, confirming the occurrence PANoptosis in the inflamed mucosa of CD (Figure 5A). Subsequently, the IFN-γ-induced epithelial cell type-specific response was evaluated using publicly available scRNA-seq datasets of IECs from CD and healthy controls (25).

**Figure 5.**
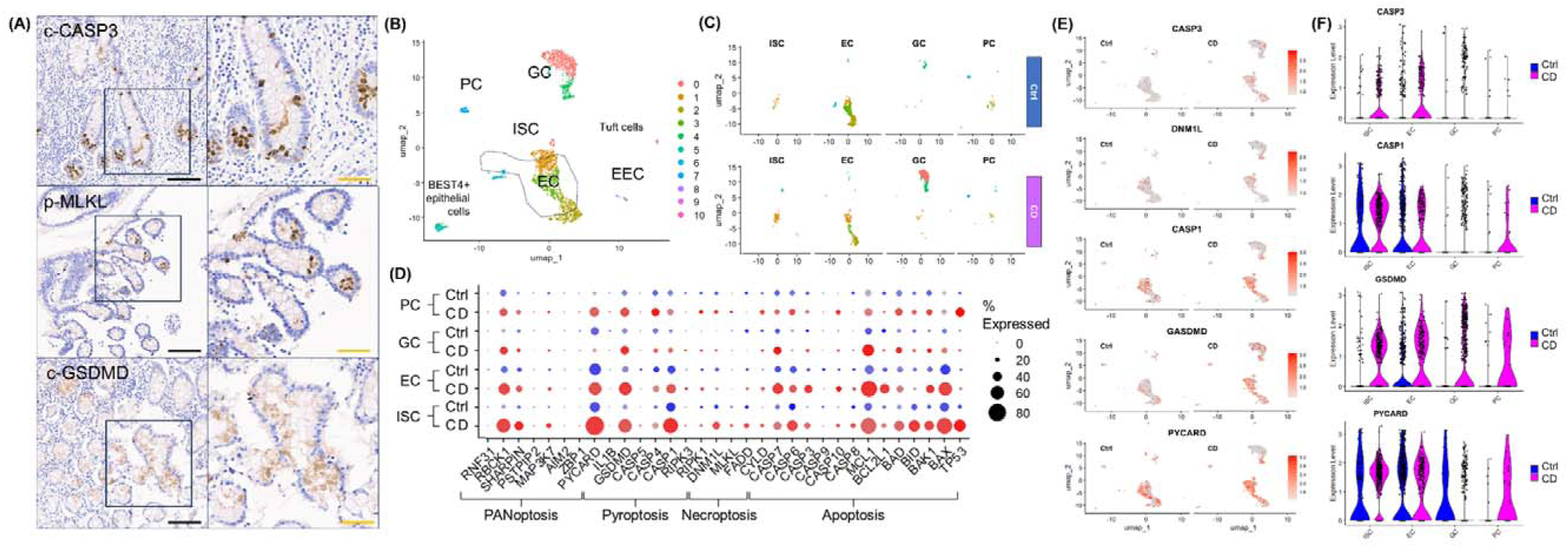
Validation of PANoptosis and Epithelial Cellular Responses in Human CD Samples. (A) Immunohistochemical staining for cleaved CASP3 (c-CASP3), phospho-MLKL (p-MLKL), and cleaved GSDMD (c-GSDMD) in surgically resected intestinal tissue from patients with CD (n=5). (B) FeaturePlot annotation based on epithelial cell type of publicly available datasets from normal and CD patients (https://www.gutcellatlas.org/). (C) FeaturePlot splited by disease status and epithelial cell types. (D) DotPlot of PCD-related gene expression in publicly available datasets from normal and CD patients. (E) ViolinPlot of key PCD molecules splited by disease status in publicly available datasets from normal and CD patients. (F) FeaturePlot of key PCD molecules splited by epithelial cell types in publicly available datasets from normal and CD patients. Scale bar: black 200 μm, orange 100 μm. Publicly available datasets was downloaded from Gut Cell Atlas (https://www.gutcellatlas.org/).

Because IFN-γ is abundant in the inflamed mucosa of CD (39), we hypothesized that scRNA-seq datasets of IECs from pediatric patients with CD would be comparable to IFN-γ-treated enteroids, whereas scRNA-seq datasets of IECs from healthy adults would be comparable to IFN-γ-free enteroids. Following the same quality control and filtering criteria applied to the scRNA-seq analysis of our enteroids, a total of 2,271 IEC datasets were identified, including 1320 IEC datasets from patients with CD (CD-IECs) and 951 IEC datasets from healthy adults (Ctrl-IECs). Similar to the IFN-γ-treated enteroids, the proportion of EC decreased in CD-IECs (57.7%), while the proportion of ISC, GC, and increased in CD-IECs (ISC 1.9%, GC 3.1%, and PC 2.0%) compared with Ctrl-IECs (EC 12.9%, ISC 3.4%, GC 28.4%, and PC 6.8%) (Figure 5B-C). Similar to enteroid models, the expression of PANoptosis-related genes was minimal in GC and PC, whereas the expression of PANoptosis-related genes was up-regulated in ISC and EC in scRNA-seq datasets from CD-IECs compared to Ctrl-IECs (Figure 5D). The expression of key PANoptosis-related genes, CASP3, DNM1L, CASP1, GSDMD, and PYCARD, was upregulated in ISC and EC, especially in CD-IECs (Figure 5E-F). These findings not only demonstrate that PANoptosis occurred in IECs of CD patients, but also that the features observed in IFN-γ-treated enteroids are similarly expressed in the scRNA-seq IEC dataset obtained from fresh patient samples.

### Effects of PCD inhibitors on IFN-γ-induced cytotoxicity

scRNA-seq of human enteroids as well as IECs showed that IFN-γ directly induces cytotoxicity in ECs via PANoptosis. In randomized clinical trials, anti-IFN-γ monoclonal antibodies failed to demonstrate efficacy in moderate to severe CD (40). Therefore, we focused on the small molecules capable of blocking IFN-γ-induced PCD and cytotoxicity in human enteroids.

Based on the previous studies, we selected the individual PCD inhibitors as potential therapeutic molecules for IFN-γ-induced cytotoxicity: Pan-caspase inhibitor, Z-VAD-FMK, to block the apoptosis pathway; RIPK1 inhibitor, necrostatin-1, and MLKL inhibitor, necrosulfonamide, to block necroptosis; caspase-1 inhibitor, AC-YVAD-CMK, and GSDMD inhibitor, LDC7559, to prevent pyroptosis; and selective JAK1 inhibitor, upadacitinib. These PCD inhibitors and upadacitinib were administered into culture medium of human enteroids treated with IFN-γ 100 pg/mL and 400 pg/mL to evaluate the organoid-forming efficiency and MTT assay (Figure 6A). Individual PCD inhibitors failed to block IFN-γ-induced cytotoxicity, whereas upadacitinib can effectively block IFN-γ-induced cytotoxicity (Figure 6B-C).

**Figure 6.**
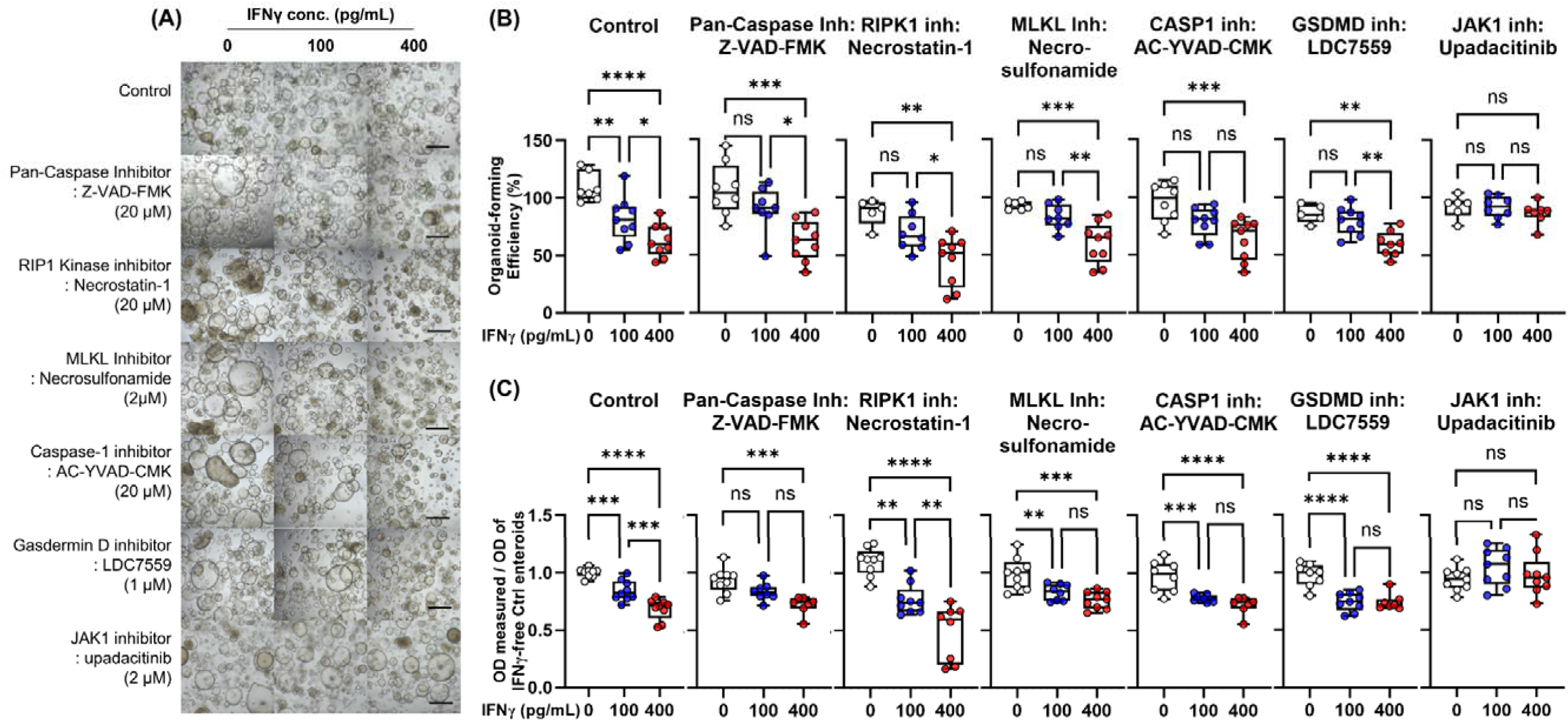
Effects of PCD inhibitors [pan-caspase inhibitor (Z-VAD-FMK), RIPK1 inhibitor (necrostatin-1), MLKL inhibitor (necrosulfonamide), caspase-1 inhibitor (AC-YVAD-CMK), GSDMD inhibitor (LDC7559), and JAK1 inhibitor (upadacitinib)] on IFN-γ-induced cytotoxicity. (A) Bright-field microscopic image of human enteroids treated with IFN-γ 0, 100 pg/mL, and 400 pg/mL in combination with PCD inhibitors. Black scale bar: 100 μm. (B) Organoid-forming efficiency. (C) Cell viability as measured by MTT assay. Assays were performed in triplicate with three Ctrl enteroids. Differences were evaluated by one-way ANOVA with multiple comparisons; ** *p* < 0.01, *** *p* < 0.001

These results suggest that individual PCD inhibitors may be insufficient to prevent IFN-γ-induced cytotoxicity. In condition of PANoptosis, even if an individual PCD is blocked, it can be bypassed by other active PCD pathways. On the other hand, upadacitinib interferes with the downstream signaling of IFN-γ by inhibiting the activity of JAK1 kinase, resulting in effective blocking of PANoptosis.

### Effects of upadacitinib on IFN-γ-treated enteroids

We then evaluated the ability of the selective JAK1 inhibitor, upadacitinib, to prevent IFN-γ-induced cytotoxicity. 2 μM upadacitinib was estimated to be equivalent to the therapeutic dose used in clinical trials (Supplementary Material S9) (41). Although up to 20 μM upadacitinib did not show any significant morphological changes in human enteroids, higher concentrations, such as 100 μM upadacitinib, induced enteroids to be small and thick-walled.

Western blotting showed that the upregulation of cleaved caspase-3, phosphorylated RIP3, phosphorylated MLKL, and cleaved form of GSDMD protein in IFN-γ-treated Ctrl and CD enteroids was prevented by upadacitinib treatment (Figure 7A). Impaired organoid-forming efficiency and cell viability in IFN-γ-treated Ctrl and CD enteroids was significantly improved by upadacitinib treatment (Figure 7B-D).

**Figure 7.**
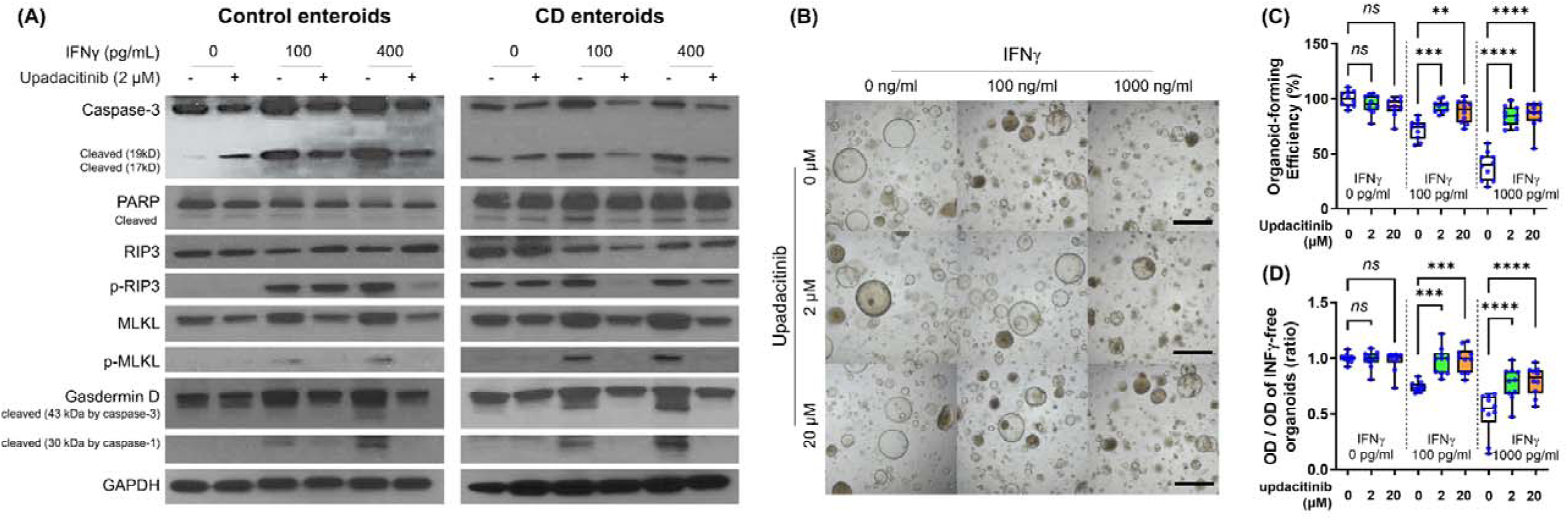
Effects of upadacitinib on IFN-γ-treated Ctrl and CD enteroids. (A) Western blotting for the key molecules of PCD signaling pathways to identify the effect of upadacitinib in Ctrl enteroid (n=3) and CD enteroid (n=3) treated with and without IFN-γ stimulation at concentrations of 100 pg/mL and 400 pg/mL. (B) Bright-field microscopic image of human enteroids treated with IFN-γ 0, 100, and 1000 pg/mL accompanied by 0, 2, and 20 μM upadacitinib. Scale bar: 100 μm. (C) Organoid-forming efficiency. (D) Cell viability as measured by MTT assay. Assays were performed in triplicate with three Ctrl enteroids. Differences were evaluated by one-way ANOVA with multiple comparisons; ** *p* < 0.01, *** *p* < 0.001, and **** *p* < 0.0001.

## Discussion

In this study, we identified that the major response observed with IFN-γ treatment of human enteroids was a decrease in organoid-forming efficiency, cell viability, and proliferation; the main PCD pathway of IFN-γ-induced PCD was PANoptosis; IFN-γ altered epithelial cell differentiation including depletion of the EC population and expansion of the GC population; PANoptosis occurs in inflamed IECs in patients with CD; and IFN-γ-induced PCD was blocked by the selective JAK1 inhibitor upadacitinib.

Although the key initiators, effectors and executioners in the main PCD pathways, including apoptosis, necroptosis, and pyroptosis, were unique and historically delineated them as distinct signaling pathways, recent studies have identified mechanistic overlap and extensive, multifaceted cross-talk among these pathways, leading to the establishment of the concept of PANoptosis (36, 42). PANoptosis was first introduced in 2019 (42) and is characterized by the combined activation of pyroptosis (“P”), apoptosis (“A”) and/or necroptosis (“N”), and is regulated by the PANoptosome complex, which is composed of key components of pyroptosis, apoptosis and/or necroptosis, such as caspase-8, NLRP3, caspase-1, GSDMD, and MLKL, that cannot be accounted for by any of the individual PCD pathways alone (30, 34, 42–49). PANoptosis is known to be important for host defense through the ZBP1-PANoptosome during influenza A virus and coronavirus infection (30, 48) and through the AIM2-PANoptosome during infections with herpes simplex virus and Francisella novicida (49). IFN-γ is primarily produced by T lymphocytes, particularly CD4+ and CD8+ T cells, as well as some natural killer (NK) cells upon viral infections. In this study, we demonstrated that IFN-γ induced up-regulation and activation of key molecules of pyroptosis, apoptosis, and necroptosis, as well as PANoptosis mediators, such as ZBP1, AIM2, TNFAIP3 (A20), SHARPIN, RBCK1 (HOIL1), and RNF31 (HOIP) in human enteroids, which indicated the main PCD of IFN-γ-treated enteroids is PANoptosis. In previous experiments with cancer cell lines, PANoptosis could not be induced by IFN-γ alone, but only by the addition of TNF to IFN-γ (30, 45). IFN-γ induces exposed cells to be sensitive to TNF signaling with upregulation of TNFR1 and TNFR2. Our study revealed that IFN-γ alone induced PANoptosis in ECs among the various intestinal cell types. Immortalized cancer cell lines have stem cell-like properties that may be resistant to IFN-γ, resulting in the different response of our patient-derived enteroid models. Furthermore, our ex vivo results were validated in human tissue samples that PANoptosis occurred in inflamed IECs of CD patients. The patient-derived enteroid model may mimic the complex responses of IECs in the human gut and could be used as an experimental model of IBD (50).

scRNA-seq for Ctrl and CD enteroids indicated that IFN-γ altered epithelial cell differentiation, including expansion of the GC population and depletion of the EC and proliferating cell populations. These findings were validated using publicly available scRNA-seq datasets of IECs from CD patients compared with those from healthy adults. Previous studies also reported that IFN-γ acts as a direct intestinal secretagogue for both GCs and PCs to release mucin and antimicrobial peptides (14, 15). This may explain why IFN-γ-induced cytotoxicity is maximized in the absorptive cell lineages, while it is minimized in the secretory lineage cells. Within the secretory lineage, GCs are more resistant to IFN-γ than PCs. Previous studies have reported that IFN-γ mediates PC cytotoxicity through suppression of mTOR, which is distinct from canonical PCD, including apoptosis, necroptosis, and pyroptosis (51). In response to IFN-γ, no differences were observed between Ctrl enteroids and CD enteroids in most aspects. Notably, however, IFN-γ-induced autophagy signaling was significantly upregulated in CD enteroids compared to Ctrl enteroids. Autophagy is known to be impaired in CD and to play an important role in CD pathophysiology (52). In our study, the expression of mTOR and autophagy-related genes induced by IFN-γ in Ctrl and CD enteroids (Supplementary Material S10). IFN-γ induced the upregulation of mTOR and autophagy-related genes in PCs, especially in CD enteroids..

IFN-γ signals mainly through the JAK/STAT pathway to achieve transcriptional activation of IFN-γ-inducible genes (53). Dysregulation of JAK/STAT signaling is recognized as a major contributor to various diseases, including IBD. In this study, IFN-γ-induced cytotoxicity in human enteroids can be prevented by the selective JAK1 inhibitor upadacitinib, but not by the individual PCD inhibitors. Dysregulated T-cell-derived IFN-γ induced pro-apoptotic gene expression and ISC injury through JAK/STAT signaling *in vivo* transplantation and organoid models (54). IFN-γ-induced cytotoxic effects on IECs were synergized with TNF-α through the caspase-8-JAK1/2-STAT1 pathway (55). Upadacitinib has demonstrated its efficacy in the treatment of moderate-to-severe CD and has been used in clinical practice (41). Recent research in the pharmaceutical industry has focused on investigating various small molecules for potential application in the treatment of IBD. Our study suggests that individual PCD inhibitors may not be effective in preventing IFN-γ-induced cytotoxicity. Furthermore, patient-derived enteroid models can be used to discover and facilitate potential therapeutic molecules.

The limitation of our study is the lack in vivo validation. While the organoid model is advanced experimental platform, it does not fully replicate the complexity of the human gut environment, including immune responses and microbial interactions. The lack of microbiome consideration was another limitation of our study. We could not account for the impact of the gut microbiome on IEC responses to IFN-γ, which may remain with the knowledge gap in understanding the response of IECs in IBD. IFN-γ effect on other cell types, such as immune cells and stromal cells, could not be evaluated due to limitation of organoid model. According to the findings of previous studies, IFN-γ in combination with TNF-α was demonstrated to induce PANoptosis in macrophages and fibroblasts (44, 45, 55). In addition, the combination of individual PCD inhibitors for the efficacy in blocking PANoptosis was not evaluated, which would have been beneficial. While individual PCD inhibitors show promise in preclinical studies, their clinical use has not been approved due to potential side effects, with the exception of Upadacitinib (56). Therefore, the combination of individual PCD inhibitors is unlikely to be beneficial in clinical application and was not evaluated in this study.

Excessive IFN-γ induces excessive IEC damage, which may contribute to the development and progression of the Th1-mediated immune disorder CD and Th1-related pathology in the gut (29, 57–59). However, the mechanism of IFN-γ-induced IEC damage in the human intestine has been poorly documented. This study demonstrated that PANoptosis is the major mechanism of IFN-γ-induced IEC injury, which was managed by a selective JAK1 inhibitor. Notably, ECs were susceptible to IFN-γ-induced PANoptosis. Our findings may be helpful in understanding the underlying pathophysiology of CD and the possible mechanisms why upadacitinib promotes mucosal healing in patients with IBD in clinical practice (41, 60).

## Resource availability

The datasets generated and/or analyzed in the current study are not publicly available prior to publication, but are available on reasonable request from the corresponding author.

## Acknowledgments

This work was supported by the National Research Foundation of Korea [NRF] grant funded by the Korea government [MSIP] [2022R1A2B5B02002092, 2019R1I1A01062205, and RS-2023-00225239] and Future Medicine 20*30 Project of the Samsung Medical Center [SMO1230011].

## Author Contributions

CL, JEK: study concept and design, data acquisition, analysis, and interpretation, and drafting of the manuscript. Y-EC, JWM: data acquisition. ERK, DKC, YK: material support, study supervision. SNH: study concept and design, data acquisition, analysis, and interpretation, drafting of the manuscript, obtained funding.

## REFERENCES

1. Zhang J. Yin and yang interplay of IFN-gamma in inflammation and autoimmune disease. J Clin Invest.. (2007) 117(4):871–3.

2. Ivashkiv LB. IFNγ: signalling, epigenetics and roles in immunity, metabolism, disease and cancer immunotherapy. Nat Rev Immunol. (2018) 18(9):545–58.

3. Zhou F. Molecular mechanisms of IFN-gamma to up-regulate MHC class I antigen processing and presentation. Int Rev Immunol. (2009) 28(3-4):239–60.

4. Malik A, Sharma D, Aguirre-Gamboa R, McGrath S, Zabala S, Weber C, et al. Epithelial IFNγ signalling and compartmentalized antigen presentation orchestrate gut immunity. Nature. (2023) 623(7989):1044-52.

5. Koh S-J, Hong SN, Park S-K, Ye BD, Kim KO, Shin JE, et al. Korean clinical practice guidelines on biologics for moderate to severe Crohn’s disease. Intest Res. (2023) 21(1):43–60.

6. Lewis JD, Parlett LE, Jonsson Funk ML, Brensinger C, Pate V, Wu Q, et al. Incidence, Prevalence, and Racial and Ethnic Distribution of Inflammatory Bowel Disease in the United States. Gastroenterology. (2023) 165(5):1197–205.e2.

7. Shivashankar R, Tremaine WJ, Harmsen WS, Loftus EV, Jr. Incidence and Prevalence of Crohn’s Disease and Ulcerative Colitis in Olmsted County, Minnesota From 1970 Through 2010. Clin Gastroenterol Hepatol. (2017) 15(6):857–63.

8. Fuss IJ, Neurath M, Boirivant M, Klein JS, de la Motte C, Strong SA, et al. Disparate CD4+ lamina propria (LP) lymphokine secretion profiles in inflammatory bowel disease. Crohn’s disease LP cells manifest increased secretion of IFN-gamma, whereas ulcerative colitis LP cells manifest increased secretion of IL-5. J Immunol. (1996) 157(3):1261–70.

9. Fais S, Capobianchi MR, Pallone F, Di Marco P, Boirivant M, Dianzani F, et al. Spontaneous release of interferon gamma by intestinal lamina propria lymphocytes in Crohn’s disease. Kinetics of in vitro response to interferon gamma inducers. Gut. (1991) 32(4):403–7.

10. Imam T, Park S, Kaplan MH, Olson MR. Effector T Helper Cell Subsets in Inflammatory Bowel Diseases. Front Immunol. (2018) 9:1212.

11. de Bruin AM, Demirel Ö, Hooibrink B, Brandts CH, Nolte MA. Interferon-γ impairs proliferation of hematopoietic stem cells in mice. Blood. (2013) 121(18):3578–85.

12. Bruewer M, Luegering A, Kucharzik T, Parkos CA, Madara JL, Hopkins AM, et al. Proinflammatory cytokines disrupt epithelial barrier function by apoptosis-independent mechanisms. J Immunol. (2003) 171(11):6164–72.

13. Bardenbacher M, Ruder B, Britzen-Laurent N, Schmid B, Waldner M, Naschberger E, et al. Permeability analyses and three dimensional imaging of interferon gamma-induced barrier disintegration in intestinal organoids. Stem Cell Res. (2019) 35:101383.

14. Farin HF, Karthaus WR, Kujala P, Rakhshandehroo M, Schwank G, Vries RG, et al. Paneth cell extrusion and release of antimicrobial products is directly controlled by immune cell-derived IFN-γ. J Exp Med. (2014) 211(7):1393–405.

15. Yue R, Wei X, Zhao J, Zhou Z, Zhong W. Essential Role of IFN-γ in Regulating Gut Antimicrobial Peptides and Microbiota to Protect Against Alcohol-Induced Bacterial Translocation and Hepatic Inflammation in Mice. Front Physiol. (2020) 11:629141.

16. Hong SN, Dunn JC, Stelzner M, Martín MG. Concise Review: The Potential Use of Intestinal Stem Cells to Treat Patients with Intestinal Failure. Stem Cells Transl Med. (2017) 6(2):666–76.

17. Sato T, Vries RG, Snippert HJ, van de Wetering M, Barker N, Stange DE, et al. Single Lgr5 stem cells build crypt-villus structures in vitro without a mesenchymal niche. Nature. (2009) 459(7244):262–5.

18. Lei NY, Jabaji Z, Wang J, Joshi VS, Brinkley GJ, Khalil H, et al. Intestinal subepithelial myofibroblasts support the growth of intestinal epithelial stem cells. PLoS One. (2014) 9(1):e84651.

19. Kim J, Koo BK, Knoblich JA. Human organoids: model systems for human biology and medicine. Nat Rev Mol Cell Biol. (2020) 21(10):571–84.

20. Lee C, An M, Joung JG, Park WY, Chang DK, Kim YH, et al. TNFα Induces LGR5+ Stem Cell Dysfunction In Patients With Crohn’s Disease. Cell Mol Gastroenterol Hepatol. (2022) 13(3):789–808.

21. Lee C, Song JH, Cha YE, Chang DK, Kim YH, Hong SN. Intestinal Epithelial Responses to IL-17 in Adult Stem Cell-derived Human Intestinal Organoids. J Crohns Colitis. (2022) 16(12):1911–23.

22. Johnson WE, Li C, Rabinovic A. Adjusting batch effects in microarray expression data using empirical Bayes methods. Biostatistics. (2007) 8(1):118–27.

23. Robinson MD, McCarthy DJ, Smyth GK. edgeR: a Bioconductor package for differential expression analysis of digital gene expression data. Bioinformatics. (2010) 26(1):139–40.

24. Franzén O, Gan LM, Björkegren JLM. PanglaoDB: a web server for exploration of mouse and human single-cell RNA sequencing data. Database (Oxford). (2019) 2019:baz046. doi: 10.1093/database/baz046.

25. Zilbauer M, James KR, Kaur M, Pott S, Li Z, Burger A, et al. A Roadmap for the Human Gut Cell Atlas. Nat Rev Gastroenterol Hepatol. (2023) 20(9):597–614.

26. Street K, Risso D, Fletcher RB, Das D, Ngai J, Yosef N, et al. Slingshot: cell lineage and pseudotime inference for single-cell transcriptomics. BMC Genomics. (2018) 19(1):477.

27. Szklarczyk D, Kirsch R, Koutrouli M, Nastou K, Mehryary F, Hachilif R, et al. The STRING database in 2023: protein-protein association networks and functional enrichment analyses for any sequenced genome of interest. Nucleic Acids Res. (2023) 51(D1):D638–d46.

28. Upton JW, Chan FK. Staying alive: cell death in antiviral immunity. Mol Cell. (2014) 54(2):273–80.

29. Nenci A, Becker C, Wullaert A, Gareus R, van Loo G, Danese S, et al. Epithelial NEMO links innate immunity to chronic intestinal inflammation. Nature. (2007) 446(7135):557–61.

30. Karki R, Lee S, Mall R, Pandian N, Wang Y, Sharma BR, et al. ZBP1-dependent inflammatory cell death, PANoptosis, and cytokine storm disrupt IFN therapeutic efficacy during coronavirus infection. Sci Immunol. (2022) 7(74):eabo6294.

31. Wang L, Zhu Y, Zhang L, Guo L, Wang X, Pan Z, et al. Mechanisms of PANoptosis and relevant small-molecule compounds for fighting diseases. Cell Death Dis. (2023) 14(12):851.

32. Grasl-Kraupp B, Ruttkay-Nedecky B, Koudelka H, Bukowska K, Bursch W, Schulte-Hermann R. In situ detection of fragmented DNA (TUNEL assay) fails to discriminate among apoptosis, necrosis, and autolytic cell death: a cautionary note. Hepatology. (1995) 21(5):1465–8.

33. Yu P, Zhang X, Liu N, Tang L, Peng C, Chen X. Pyroptosis: mechanisms and diseases. Signal Transduct Target Ther. (2021) 6(1):128.

34. Kuriakose T, Man SM, Malireddi RK, Karki R, Kesavardhana S, Place DE, et al. ZBP1/DAI is an innate sensor of influenza virus triggering the NLRP3 inflammasome and programmed cell death pathways. Sci Immunol. (2016) 1(2):aag2045. doi: 10.1126/sciimmunol.aag2045.

35. Guo L, Lin L, Wang X, Gao M, Cao S, Mai Y, et al. Resolving Cell Fate Decisions during Somatic Cell Reprogramming by Single-Cell RNA-Seq. Mol Cell. (2019) 73(4):815–29.e7.

36. Wang Y, Kanneganti TD. From pyroptosis, apoptosis and necroptosis to PANoptosis: A mechanistic compendium of programmed cell death pathways. Comput Struct Biotechnol J. (2021) 19:4641–57.

37. Wei S, Chen Z, Ling X, Zhang W, Jiang L. Comprehensive analysis illustrating the role of PANoptosis-related genes in lung cancer based on bioinformatic algorithms and experiments. Front Pharmacol. (2023) 14:1115221.

38. Samir P, Malireddi RKS, Kanneganti TD. The PANoptosome: A Deadly Protein Complex Driving Pyroptosis, Apoptosis, and Necroptosis (PANoptosis). Front Cell Infect Microbiol. (2020) 10:238.

39. Neurath MF. Cytokines in inflammatory bowel disease. Nature Reviews Immunology. (2014) 14(5):329–42.

40. Reinisch W, de Villiers W, Bene L, Simon L, Rácz I, Katz S, et al. Fontolizumab in moderate to severe Crohn’s disease: a phase 2, randomized, double-blind, placebo-controlled, multiple-dose study. Inflamm Bowel Dis. (2010) 16(2):233–42.

41. Loftus EV, Jr., Panés J, Lacerda AP, Peyrin-Biroulet L, D’Haens G, Panaccione R, et al. Upadacitinib Induction and Maintenance Therapy for Crohn’s Disease. N Engl J Med. (2023) 388(21):1966–80.

42. Malireddi RKS, Kesavardhana S, Kanneganti TD. ZBP1 and TAK1: Master Regulators of NLRP3 Inflammasome/Pyroptosis, Apoptosis, and Necroptosis (PAN-optosis). Front Cell Infect Microbiol. (2019) 9:406.

43. Karki R, Sharma BR, Lee E, Banoth B, Malireddi RKS, Samir P, et al. Interferon regulatory factor 1 regulates PANoptosis to prevent colorectal cancer. JCI Insight. (2020) 5(12).

44. Malireddi RKS, Karki R, Sundaram B, Kancharana B, Lee S, Samir P, et al. Inflammatory Cell Death, PANoptosis, Mediated by Cytokines in Diverse Cancer Lineages Inhibits Tumor Growth. Immunohorizons. (2021) 5(7):568–80.

45. Karki R, Sharma BR, Tuladhar S, Williams EP, Zalduondo L, Samir P, et al. Synergism of TNF-α and IFN-γ Triggers Inflammatory Cell Death, Tissue Damage, and Mortality in SARS-CoV-2 Infection and Cytokine Shock Syndromes. Cell. (2021) 184(1):149–68.e17.

46. Wang Y, Pandian N, Han JH, Sundaram B, Lee S, Karki R, et al. Single cell analysis of PANoptosome cell death complexes through an expansion microscopy method. Cell Mol Life Sci. (2022) 79(10):531.

47. Sundaram B, Pandian N, Mall R, Wang Y, Sarkar R, Kim HJ, et al. NLRP12-PANoptosome activates PANoptosis and pathology in response to heme and PAMPs. Cell. (2023) 186(13):2783–801.e20.

48. Zheng M, Karki R, Vogel P, Kanneganti TD. Caspase-6 Is a Key Regulator of Innate Immunity, Inflammasome Activation, and Host Defense. Cell. (2020) 181(3):674–87.e13.

49. Lee S, Karki R, Wang Y, Nguyen LN, Kalathur RC, Kanneganti TD. AIM2 forms a complex with pyrin and ZBP1 to drive PANoptosis and host defence. Nature. (2021) 597(7876):415–9.

50. Nanki K, Fujii M, Shimokawa M, Matano M, Nishikori S, Date S, et al. Somatic inflammatory gene mutations in human ulcerative colitis epithelium. Nature. (2020) 577(7789):254–9.

51. Araujo A, Safronova A, Burger E, López-Yglesias A, Giri S, Camanzo ET, et al. IFN-γ mediates Paneth cell death via suppression of mTOR. Elife. (2021) 10:e60478. doi: 10.7554/eLife.60478.

52. Alula KM, Theiss AL. Autophagy in Crohn’s Disease: Converging on Dysfunctional Innate Immunity. Cells. (2023) 12(13):1779. doi: 10.3390/cells12131779.

53. Hu X, Li J, Fu M, Zhao X, Wang W. The JAK/STAT signaling pathway: from bench to clinic. Signal Transduct Target Ther. (2021) 6(1):402.

54. Takashima S, Martin ML, Jansen SA, Fu Y, Bos J, Chandra D, et al. T cell-derived interferon-γ programs stem cell death in immune-mediated intestinal damage. Sci Immunol. (2019) 4(42).

55. Woznicki JA, Saini N, Flood P, Rajaram S, Lee CM, Stamou P, et al. TNF-α synergises with IFN-γ to induce caspase-8-JAK1/2-STAT1-dependent death of intestinal epithelial cells. Cell Death Dis. (2021) 12(10):864.

56. Chauvier D, Ankri S, Charriaut-Marlangue C, Casimir R, Jacotot E. Broad-spectrum caspase inhibitors: from myth to reality? Cell Death Differ. (2007) 14(2):387–91.

57. Berg DJ, Davidson N, Kühn R, Müller W, Menon S, Holland G, et al. Enterocolitis and colon cancer in interleukin-10-deficient mice are associated with aberrant cytokine production and CD4(+) TH1-like responses. J Clin Invest. (1996) 98(4):1010–20.

58. Ito R, Shin-Ya M, Kishida T, Urano A, Takada R, Sakagami J, et al. Interferon-gamma is causatively involved in experimental inflammatory bowel disease in mice. Clin Exp Immunol. (2006) 146(2):330–8.

59. Powrie F, Leach MW, Mauze S, Menon S, Caddle LB, Coffman RL. Inhibition of Th1 responses prevents inflammatory bowel disease in scid mice reconstituted with CD45RBhi CD4+ T cells. Immunity. (1994) 1(7):553–62.

60. Panés J, Louis E, Bossuyt P, Joshi N, Lee WJ, Lacerda AP, et al. Induction of endoscopic response, remission, and ulcer-free endoscopy with upadacitinib is associated with improved clinical outcomes and quality of life in patients with Crohn’s disease. Inflamm Bowel Dis. (2024) izae200. doi: 10.1093/ibd/izae200. Online ahead of print.

