## Supplementary material for "IFN-γ-Induced Intestinal Epithelial Cell-Type-Specific Programmed Cell Death: PANoptosis and Its Modulation in Crohn’s Disease": supple materials

**Supplementary Materials**

Supplementary Material S1. (A) Confocal microscopy image of the EdU assay on the Ctrl enteroid treated with different concentrations of IFN-γ. (B) Representative confocal microscopy image of patient-derived enteroid of similar shape and size treated with and without IFN-γ. Scale bar = 50 μm.

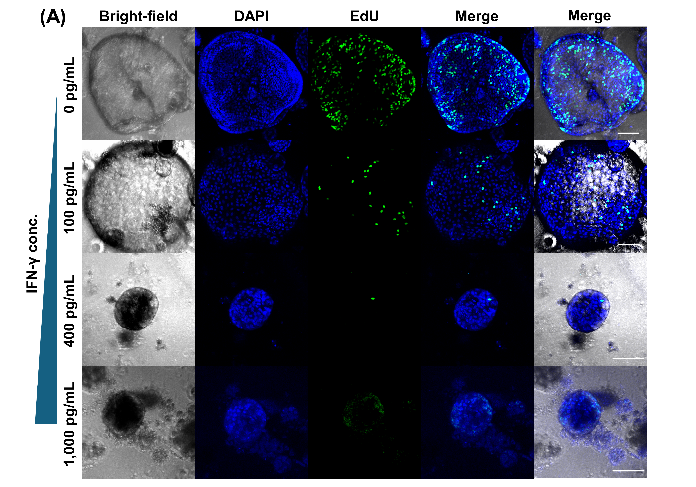

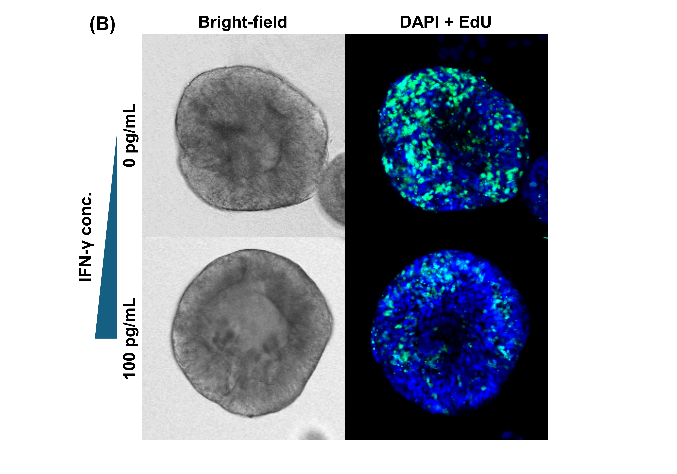

Supplementary Material S2. Gene expression associated with programmed cell death that changed with IFN-γ treatment in human enteroids

|  |  | Mean FPKM of IFN-γ-free enteroids | S.D. of FPKM for IFN-γ-free enteroids | Mean FPKM of IFN-γ-treated enteroids | S.D. of FPKM for IFN-γ-treated enteroids | log2FC | *p*-value | Adj. *p*-value | FDR *q*-value |
| --- | --- | --- | --- | --- | --- | --- | --- | --- | --- |
|  | Signifcant gene in *t*-test (*p* < 0.01, adj. *p* < 0.01, and *q* < 0.05) | | | | | | | | |
| TP53 |  | 71.009 | 5.777 | 62.754 | 4.723 | -0.18 | 1.33E-04 | 4.00E-03 | 2.35E-04 |
| BCL2L1 |  | 536.495 | 31.299 | 480.732 | 27.277 | -0.16 | 9.01E-06 | 2.70E-04 | 1.69E-05 |
| MCL1 |  | 342.195 | 18.222 | 454.584 | 24.192 | 0.41 | 1.67E-14 | 5.00E-13 | 1.25E-13 |
| BAK1 |  | 135.437 | 8.224 | 248.330 | 16.741 | 0.87 | 0.00E+00 | 0.00E+00 | 0.00E+00 |
| CASP8 |  | 44.984 | 3.411 | 69.869 | 6.637 | 0.64 | 5.08E-12 | 1.52E-10 | 1.90E-11 |
| CASP10 |  | 36.503 | 4.016 | 80.381 | 8.020 | 1.14 | 2.00E-15 | 6.00E-14 | 2.00E-14 |
| CASP3 |  | 74.142 | 4.586 | 93.636 | 6.338 | 0.34 | 1.52E-10 | 4.57E-09 | 4.57E-10 |
| CASP7 |  | 110.490 | 10.833 | 169.832 | 12.911 | 0.62 | 1.69E-14 | 5.06E-13 | 1.01E-13 |
| TNFSF10 |  | 109.138 | 17.094 | 253.148 | 39.849 | 1.21 | 2.21E-11 | 6.62E-10 | 7.36E-11 |
| TNFRSF1B |  | 46.301 | 6.722 | 83.895 | 7.954 | 0.86 | 8.88E-15 | 2.58E-13 | 6.44E-14 |
| FAS |  | 56.749 | 4.955 | 75.145 | 6.632 | 0.41 | 1.67E-09 | 5.00E-08 | 4.17E-09 |
| CYLD |  | 30.903 | 2.973 | 48.965 | 4.100 | 0.66 | 4.31E-14 | 1.29E-12 | 2.15E-13 |
| RIPK1 |  | 66.665 | 4.017 | 77.999 | 4.519 | 0.23 | 2.90E-08 | 8.69E-07 | 6.21E-08 |
| MLKL |  | 19.901 | 2.826 | 40.499 | 7.110 | 1.03 | 1.57E-09 | 4.72E-08 | 4.29E-09 |
| CASP1 |  | 22.754 | 4.491 | 138.742 | 22.000 | 2.61 | 5.79E-13 | 1.74E-11 | 2.48E-12 |
| CASP5 |  | 11.110 | 2.856 | 20.948 | 5.676 | 0.91 | 3.12E-06 | 9.35E-05 | 6.23E-06 |
| GSDMD |  | 71.910 | 6.441 | 129.459 | 9.471 | 0.85 | 0.00E+00 | 0.00E+00 | 0.00E+00 |
| ZBP1 |  | 0.908 | 0.350 | 15.312 | 5.015 | 4.08 | 8.07E-09 | 2.42E-07 | 1.86E-08 |
|  | Non-signifcant gene in *t*-test (*p* > 0.01, adj. *p* > 0.01, or *q* > 0.05) | | | | | | | | |
| BAX |  | 130.591 | 8.723 | 120.477 | 8.780 | -0.12 | 0.00278 | 0.08343 | 0.00363 |
| CASP9 |  | 20.370 | 0.885 | 21.080 | 0.837 | 0.05 | 0.02680 | 0.80389 | 0.03216 |
| TNFRSF1A |  | 46.301 | 6.722 | 83.895 | 7.954 | 0.86 | 0.00257 | 0.07712 | 0.00351 |
| FADD |  | 43.530 | 2.022 | 43.687 | 1.933 | 0.01 | 0.82357 | 1.00000 | 0.82357 |
| RIPK3 |  | 42.515 | 4.997 | 44.992 | 4.283 | 0.08 | 0.14300 | 1.00000 | 0.15322 |
| NLRC4 |  | 0.538 | 0.130 | 0.639 | 0.122 | 0.25 | 0.03167 | 0.95010 | 0.03654 |
| CASP4 |  | 86.885 | 9.398 | 102.656 | 15.936 | 0.24 | 0.00230 | 0.06907 | 0.00329 |
| MAP3K7 |  | 40.851 | 1.987 | 39.451 | 1.580 | -0.05 | 0.03566 | 1.00000 | 0.03963 |
| PSTPIP2 |  | 41.998 | 2.939 | 42.775 | 3.145 | 0.03 | 0.47598 | 1.00000 | 0.49239 |

Supplementary Material S3. Pathway analysis and Gene set enrichment analysis of

1. REACTOME pathway analysis

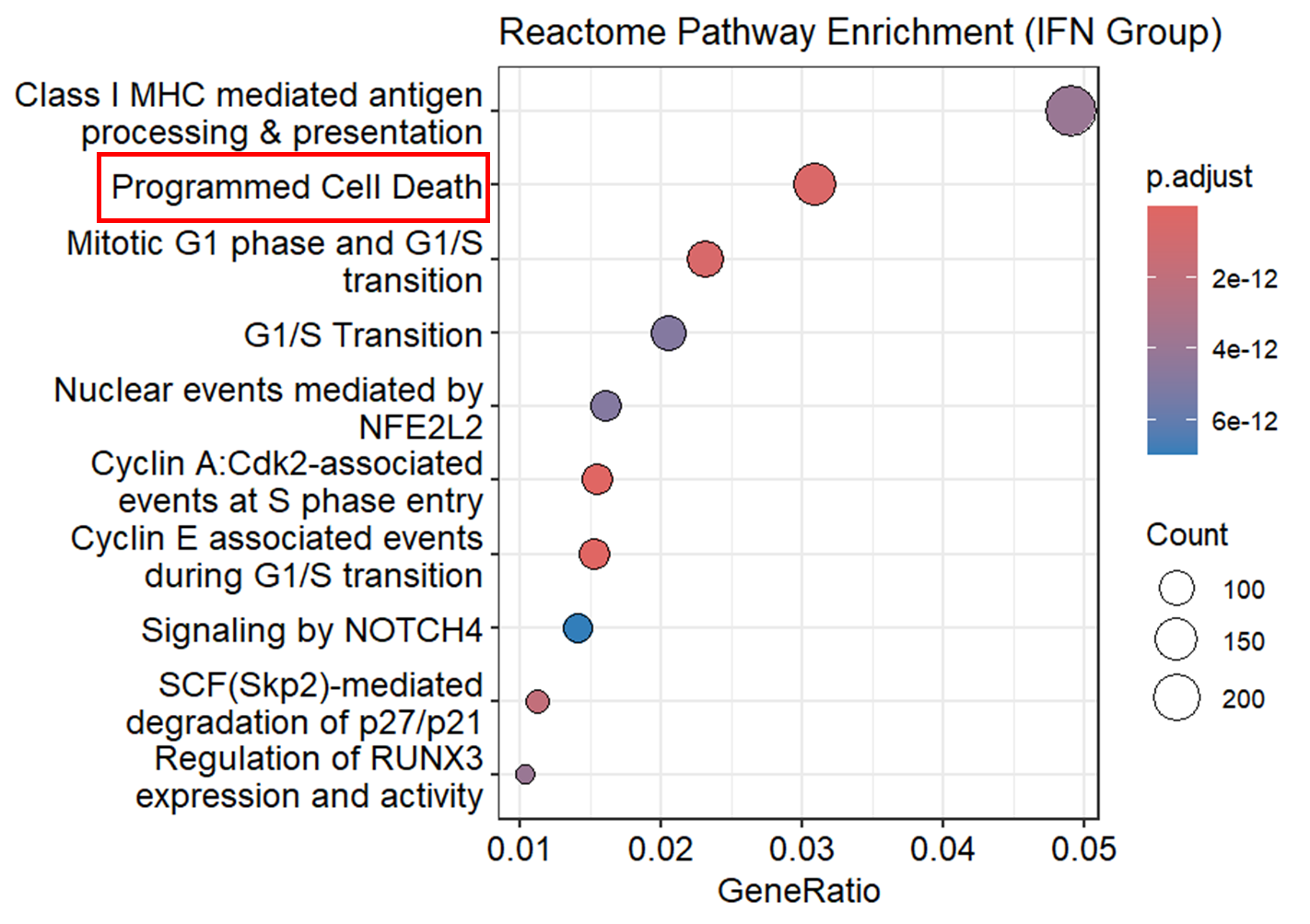

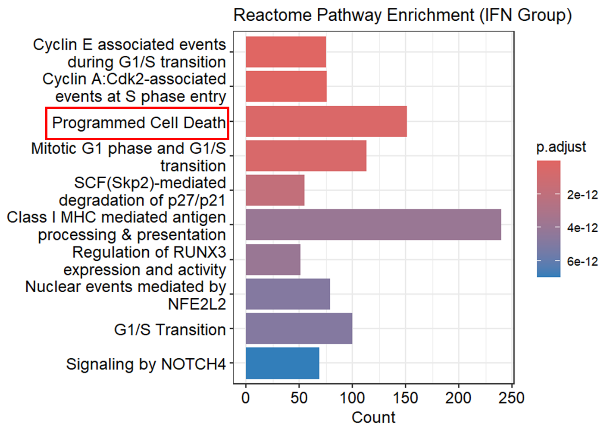

Top 20 REACTOME pathway

| **ID** | **Description** | **Gene**  **Ratio** | **Bg Ratio** | **Rich Factor** | **Fold Enrichment** | **z-Score** | **p-value** | **p. adjust** | **Gene ID** |
| --- | --- | --- | --- | --- | --- | --- | --- | --- | --- |
| R-HSA-69202 | Cyclin E associated events during G1/S transition | 75/4886 | 83/11146 | 0.90361 | 2.06134 | 8.57403 | 4.45E-19 | 7.21E-16 | geneID |
| R-HSA-69656 | Cyclin A:Cdk2-associated events at S phase entry | 76/4886 | 85/11146 | 0.89412 | 2.03967 | 8.50037 | 1.15E-18 | 9.34E-16 | PSMB8/PSMB9/PSME1/PSME2/PSMB10/PSMA4/PSMA1/PSMA6/MAX/UBB/PSMD11/PSMB7/UBC/PSMA3/PSMA2/LIN52/PSMC4/PSMA5/E2F5/PSMB2/PSMA7/PSMD3/PSMC1/PSMF1/PSMB4/PSMB1/CDKN1A/CDK2/CABLES1/PSMB3/MYC/PSMD13/PSMC6/WEE1/PSMB6/PSMD7/RBBP4/PSMD12/PSMD14/PSMB5/PSMD1/RBL2/TFDP2/CCNH/PSME3/PSMC2/PSMD2/CDC25A/PSMD6/PSMC3/RB1/E2F4/E2F1/CDK7/PSMD5/CCND1/PSMD9/CDK4/PSMD4/LIN37/LIN9/PTK6/CDKN1B/CCNE2/TFDP1/LIN54/SKP1/PSMC5/PSMD8/UBA52/SKP2/RPS27A/CKS1B/CCNE1/MNAT1 |
| R-HSA-5357801 | Programmed Cell Death | 151/4886 | 213/11146 | 0.70892 | 1.61720 | 8.03477 | 7.08E-16 | 3.82E-13 | PSMB8/PSMB9/PSME1/CDC25B/PSME2/PSMB10/PSMA4/PSMA1/PSMA6/MAX/UBB/PSMD11/PSMB7/UBC/PSMA3/PSMA2/LIN52/PSMC4/PSMA5/E2F5/PSMB2/PSMA7/PSMD3/PSMC1/PSMF1/PSMB4/PSMB1/CDKN1A/CDK2/CABLES1/PSMB3/MYC/PSMD13/PSMC6/WEE1/PSMB6/PSMD7/RBBP4/PSMD12/PSMD14/PSMB5/PSMD1/RBL2/TFDP2/CCNH/PSME3/PSMC2/PSMD2/CDC25A/PSMD6/PSMC3/RB1/E2F4/E2F1/CDK7/PSMD5/CCND1/PSMD9/CDK4/PSMD4/LIN37/LIN9/PTK6/CDKN1B/CCNE2/TFDP1/LIN54/SKP1/PSMC5/PSMD8/UBA52/SKP2/RPS27A/CKS1B/CCNE1/MNAT1 |
| R-HSA-453279 | Mitotic G1 phase and G1/S transition | 113/4886 | 149/11146 | 0.75839 | 1.73005 | 7.92567 | 1.29E-15 | 5.21E-13 | PSMB8/PSMB9/PSME1/IRF1/PSME2/PSMB10/CASP1/BAK1/PSMA4/PSMA1/PSMA6/GSDMD/UBB/PSMD11/PSMB7/PMAIP1/CASP8/UBC/PSMA3/TNFSF10/DYNLL2/BIRC3/PSMA2/CASP7/PSMC4/DAPK1/MLKL/STAT3/PSMA5/PSMB2/TICAM1/PSMA7/IRF2/CASP3/PSMD3/FAS/PSMC1/PSMF1/PSMB4/GSN/PPP3CC/PSMB1/TP63/CDKN2A/NMT1/PSMB3/AVEN/CYCS/PSMD13/PSMC6/PSMB6/PSMD7/STK24/RIPK1/PSMD12/UACA/PSMD14/PSMB5/CFLAR/HMGB1/PSMD1/PTK2/MAGED1/PLEC/YWHAB/CASP5/CASP6/ARHGAP10/BMF/TFDP2/BAD/APPL1/DAPK2/TJP2/PSME3/C1QBP/YWHAQ/PSMC2/PPP3R1/DNM1L/KPNA1/CASP9/PSMD2/TLR4/UBE2L3/SATB1/BID/PSMD6/CARD8/CLSPN/PSMC3/CDC37/STUB1/BIRC2/CASP4/OCLN/PAK2/E2F1/CHMP4B/IL18/KPNB1/DCC/TP73/PSMD5/HMGB2/DAPK3/FADD/PSMD9/PSMD4/UNC5B/DIABLO/TRAF2/LMNB1/YWHAZ/XIAP/FNTA/SFN/CHMP4A/IL1A/CHMP3/TFDP1/TNFRSF10B/CHMP4C/SDCBP/BBC3/DBNL/IL1B/ROCK1/BCL2L11/RIPK3/ADD1/YWHAG/OMA1/PSMC5/HSP90AA1/PSMD8/UNC5A/MAPK8/UBA52/PPP1R13B/OPA1/PDCD6IP/CHMP6/CHMP7/PRKCQ/BCAP31/RPS27A/APIP/OGT/CHMP2A/DFFB |
| R-HSA-187577 | SCF(Skp2)-mediated degradation of p27/p21 | 55/4886 | 60/11146 | 0.91667 | 2.09111 | 7.48663 | 5.65E-15 | 1.83E-12 | PSMB8/PSMB9/PSME1/PSME2/PSMB10/PSMA4/PSMA1/PSMA6/JAK2/MAX/UBB/LYN/PSMD11/PSMB7/UBC/PSMA3/PSMA2/LIN52/PSMC4/PSMA5/E2F5/PSMB2/PSMA7/PSMD3/PSMC1/PSMF1/PSMB4/PSMB1/CDKN1A/CDK2/CDKN2A/CABLES1/PSMB3/MYC/PPP2CA/PSMD13/CCND3/PSMC6/WEE1/PSMB6/PSMD7/RPA1/RBBP4/PSMD12/PSMD14/PSMB5/GMNN/MCM7/CCND2/E2F3/PSMD1/CDKN2D/RBL1/TYMS/RBL2/TFDP2/PPP2CB/CCNH/PSME3/PSMC2/MCM5/PSMD2/CDC25A/DYRK1A/TK1/PSMD6/POLE2/MCM8/FBXO5/PSMC3/RB1/POLA1/POLE4/E2F4/E2F1/CDK7/PSMD5/CCND1/POLE/PSMD9/MYBL2/CDK4/PSMD4/ORC2/MCM10/LIN37/LIN9/MCM2/PTK6/CDKN1B/CDC45/CCNE2/TFDP1/ORC5/POLE3/MCM4/LIN54/E2F2/SKP1/SRC/CDKN2C/PSMC5/PSMD8/CDC6/UBA52/ORC4/SKP2/RRM2/RPS27A/CDKN2B/CKS1B/CCNE1/MNAT1 |
| R-HSA-983169 | Class I MHC mediated antigen processing & presentation | 240/4886 | 381/11146 | 0.62992 | 1.43698 | 7.66744 | 1.60E-14 | 3.97E-12 | PSMB8/PSMB9/PSME1/PSME2/PSMB10/PSMA4/PSMA1/PSMA6/UBB/PSMD11/PSMB7/UBC/PSMA3/PSMA2/PSMC4/PSMA5/PSMB2/PSMA7/PSMD3/PSMC1/PSMF1/PSMB4/PSMB1/CDKN1A/CDK2/PSMB3/PSMD13/PSMC6/PSMB6/PSMD7/PSMD12/PSMD14/PSMB5/PSMD1/PSME3/PSMC2/PSMD2/PSMD6/PSMC3/PSMD5/CCND1/PSMD9/CDK4/PSMD4/PTK6/CDKN1B/CCNE2/SKP1/PSMC5/PSMD8/UBA52/SKP2/RPS27A/CKS1B/CCNE1 |
| R-HSA-8941858 | Regulation of RUNX3 expression and activity | 51/4886 | 55/11146 | 0.92727 | 2.11531 | 7.32520 | 1.71E-14 | 3.97E-12 | TAP1/PSMB8/PSMB9/HLA-E/TAP2/PSME1/B2M/PSME2/PSMB10/HLA-A/UBE2L6/HLA-C/HLA-B/PSMA4/PSMA1/HLA-F/DTX3L/PSMA6/RNF213/CTSS/UBB/PSMD11/PSMB7/FBXO6/TRIM21/ERAP2/FBXO7/UBR2/TRIM69/UBC/PSMA3/RNF114/CUL7/PSMA2/KCTD7/UBA6/PSMC4/CUL2/ITGB5/RBCK1/PSMA5/UBOX5/PSMB2/IKBKG/RNF19B/HERC6/TAPBP/PSMA7/UBA7/UBE2Z/PSMD3/KLHL22/SEC24C/PSMC1/PSMF1/PSMB4/ASB1/UBE2D3/ERAP1/PDIA3/PSMB1/VAMP3/MKRN1/KLHL5/FBXO2/ZNRF1/PSMB3/STX4/MYD88/FBXL5/PIK3C3/PSMD13/SEC31A/ITGAV/FBXO31/CDC27/PSMC6/RNF130/PSMB6/SOCS1/PSMD7/RNF34/CUL3/KLHL3/PSMD12/VHL/PSMD14/LMO7/PSMB5/RNF115/UBE2E3/FBXO21/TRIM11/HMGB1/FBXO10/PSMD1/UBE2M/SEC24A/KLHL21/ASB7/UBE3B/VAMP8/HERC1/FBXO4/UBE2Q1/PSME3/UBE2V2/ASB14/ATG14/PSMC2/HLA-G/RNF123/RNF217/RNF126/TRIP12/UBE2J2/SEC61B/PSMD2/TLR4/UBE2L3/HERC4/UBE2Q2/UBE2F/RNF6/LONRF1/PSMD6/DZIP3/HERC5/KEAP1/UBE2G1/PSMC3/CBLB/TRAF7/STUB1/CDC16/UBE2E1/RNF7/UBE2O/SOCS3/ASB8/ANAPC2/HECTD3/KLHL2/RBX1/FBXO41/RNF4/CALR/UBE2D2/RLIM/LRSAM1/SEC22B/UBE4A/FBXW7/RNF144B/IKBKB/SEC24B/LRRC41/PSMD5/HACE1/ASB6/UBE2N/PSMD9/RNF19A/LNPEP/FBXO44/KLHL13/UNKL/KCTD6/FBXL18/PSMD4/TRIM37/UBE2E2/SEC23A/UBE3D/CHUK/FBXO9/TRAIP/BLMH/NCF1/CYBB/DET1/FBXW2/S100A1/ANAPC10/ANAPC13/CDC34/RNF14/RNF138/TRIM32/FBXO22/SEC13/RCHY1/FBXW10/SAR1B/THOP1/FBXL20/SKP1/UBE2K/KBTBD8/FBXW11/HERC3/UBA3/ATG7/SPSB1/FBXL3/FBXO32/PSMC5/FBXL12/PSMD8/FBXW5/MYLIP/WSB1/UBR4/RNF41/UBA52/ZNRF2/TIRAP/UBA1/SKP2/KLHL25/ANAPC5/HSPA5/MIB2/MGRN1/BCAP31/CYBA/CCNF/FBXW9/RPS27A/ARIH2/UBA5/FBXL22/FBXL19/FBXL15/FBXL13/FBXL14/CDC26/SIAH1/FBXL8/RNF25 |
| R-HSA-9759194 | Nuclear events mediated by NFE2L2 | 79/4886 | 97/11146 | 0.81443 | 1.85789 | 7.49700 | 2.50E-14 | 4.87E-12 | PSMB8/PSMB9/PSME1/PSME2/PSMB10/PSMA4/PSMA1/PSMA6/UBB/PSMD11/PSMB7/UBC/PSMA3/PSMA2/PSMC4/PSMA5/PSMB2/PSMA7/PSMD3/PSMC1/PSMF1/PSMB4/PSMB1/CDKN2A/TGFB1/PSMB3/MDM2/PSMD13/PSMC6/PSMB6/PSMD7/PSMD12/PSMD14/PSMB5/PSMD1/PSME3/PSMC2/PSMD2/PSMD6/PSMC3/PSMD5/PSMD9/PSMD4/SRC/PSMC5/CBFB/PSMD8/UBA52/RUNX3/RPS27A/EP300 |
| R-HSA-69206 | G1/S Transition | 100/4886 | 131/11146 | 0.76336 | 1.74138 | 7.54077 | 2.71E-14 | 4.87E-12 | PSMB8/PSMB9/PSME1/PSME2/PSMB10/PSMA4/PSMA1/PSMA6/UBB/PSMD11/PSMB7/UBC/PSMA3/PSMA2/PSMC4/PSMA5/PSMB2/PSMA7/GSTA1/PSMD3/PSMC1/PSMF1/PSMB4/CCL2/PSMB1/CDKN2A/AREG/PSMB3/RELA/MYC/PSMD13/PSMC6/BACH1/PSMB6/ABCC3/PSMD7/SRXN1/PSMD12/PSMD14/PSMB5/PSMD1/GCLM/SQSTM1/ME1/GCLC/PSME3/PSMC2/TXN/TXNRD1/PSMD2/SP1/PRKAA2/PSMD6/KEAP1/MAFG/PSMC3/TALDO1/RBX1/SLC7A11/ATF4/NQO1/PSMD5/PSMD9/PSMD4/PGD/NFKB1/ABCC1/ABCF2/GSR/SKP1/G6PD/PSMC5/PSMD8/UBA52/TKT/RPS27A/MAFK/NOTCH1/EP300 |
| R-HSA-9013694 | Signaling by NOTCH4 | 69/4886 | 82/11146 | 0.84146 | 1.91956 | 7.38344 | 4.57E-14 | 6.99E-12 | PSMB8/PSMB9/PSME1/PSME2/PSMB10/PSMA4/PSMA1/PSMA6/MAX/UBB/PSMD11/PSMB7/UBC/PSMA3/PSMA2/LIN52/PSMC4/PSMA5/E2F5/PSMB2/PSMA7/PSMD3/PSMC1/PSMF1/PSMB4/PSMB1/CDKN1A/CDK2/CABLES1/PSMB3/MYC/PPP2CA/PSMD13/PSMC6/WEE1/PSMB6/PSMD7/RPA1/RBBP4/PSMD12/PSMD14/PSMB5/GMNN/MCM7/PSMD1/RBL1/TYMS/RBL2/TFDP2/PPP2CB/CCNH/PSME3/PSMC2/MCM5/PSMD2/CDC25A/TK1/PSMD6/POLE2/MCM8/FBXO5/PSMC3/RB1/POLA1/POLE4/E2F4/E2F1/CDK7/PSMD5/CCND1/POLE/PSMD9/CDK4/PSMD4/ORC2/MCM10/LIN37/LIN9/MCM2/PTK6/CDKN1B/CDC45/CCNE2/TFDP1/ORC5/POLE3/MCM4/LIN54/SKP1/PSMC5/PSMD8/CDC6/UBA52/ORC4/SKP2/RRM2/RPS27A/CKS1B/CCNE1/MNAT1 |
| R-HSA-69242 | S Phase | 118/4886 | 162/11146 | 0.72840 | 1.66162 | 7.49407 | 4.74E-14 | 6.99E-12 | PSMB8/PSMB9/PSME1/PSME2/PSMB10/PSMA4/PSMA1/PSMA6/UBB/PSMD11/PSMB7/UBC/PSMA3/PSMA2/APH1B/SMAD3/PSMC4/PSMA5/PSMB2/PSMA7/PSMD3/PSMC1/PSMF1/PSMB4/PSMB1/PSENEN/PSMB3/PSMD13/PSMC6/PSMB6/PSMD7/SNW1/PSMD12/PSMD14/PSMB5/PSMD1/PSEN2/RBPJ/APH1A/PSME3/PSMC2/PSMD2/PSMD6/PSMC3/JAG1/RBX1/NOTCH4/ADAM10/MAML3/FBXW7/NCSTN/PSMD5/PSMD9/PSEN1/PSMD4/NOTCH2/KAT2B/YWHAZ/SKP1/DLL4/PSMC5/PSMD8/UBA52/MAML1/MAML2/HES1/RPS27A/NOTCH1/EP300 |
| R-HSA-450408 | AUF1 (hnRNP D0) binds and destabilizes mRNA | 51/4886 | 56/11146 | 0.91071 | 2.07753 | 7.14148 | 1.10E-13 | 1.49E-11 | PSMB8/PSMB9/PSME1/CDC25B/PSME2/PSMB10/PSMA4/PSMA1/PSMA6/MAX/UBB/PSMD11/PSMB7/UBC/PSMA3/PSMA2/LIN52/PSMC4/PSMA5/E2F5/PSMB2/PSMA7/PSMD3/PSMC1/PSMF1/PSMB4/PSMB1/CDKN1A/CDK2/CABLES1/PSMB3/MYC/PSMD13/CDC27/PSMC6/WEE1/PSMB6/PSMD7/RPA1/RBBP4/PSMD12/PSMD14/PSMB5/GMNN/MCM7/PSMD1/RBL2/TFDP2/CCNH/PSME3/PSMC2/MCM5/RFC2/ANAPC16/POLD1/PSMD2/CDC25A/PDS5B/PSMD6/POLE2/LIG1/MCM8/PSMC3/RB1/CDC16/UBE2E1/POLA1/ANAPC2/GINS2/RBX1/FEN1/POLE4/E2F4/E2F1/CDK7/PSMD5/CCND1/POLE/PSMD9/CDK4/PSMD4/ORC2/RFC5/LIN37/LIN9/GINS4/POLD3/MCM2/PTK6/CDKN1B/CDC45/ANAPC10/CCNE2/TFDP1/ORC5/POLE3/ESCO2/MCM4/CDCA5/LIN54/RFC4/SKP1/RFC3/PSMC5/PSMD8/CDC6/UBA52/ORC4/GINS3/SKP2/ANAPC5/POLD4/RPS27A/CKS1B/CDC26/SMC1A/CCNE1/MNAT1 |
| R-HSA-9020702 | Interleukin-1 signaling | 89/4886 | 115/11146 | 0.77391 | 1.76546 | 7.28942 | 1.75E-13 | 2.18E-11 | PSMB8/PSMB9/PSME1/PSME2/PSMB10/PSMA4/PSMA1/PSMA6/UBB/PSMD11/PSMB7/HSPB1/UBC/PSMA3/PSMA2/PSMC4/PSMA5/PSMB2/PSMA7/PSMD3/PSMC1/PSMF1/PSMB4/PSMB1/HSPA1A/PSMB3/PSMD13/PSMC6/PSMB6/PSMD7/PSMD12/PSMD14/PSMB5/PSMD1/EIF4G1/PSME3/PSMC2/HSPA1B/PSMD2/PSMD6/PSMC3/PSMD5/PSMD9/PSMD4/PABPC1/PSMC5/PSMD8/UBA52/HSPA8/RPS27A/HNRNPD |
| R-HSA-8878159 | Transcriptional regulation by RUNX3 | 77/4886 | 96/11146 | 0.80208 | 1.82972 | 7.21301 | 2.53E-13 | 2.93E-11 | PSMB8/NLRC5/PSMB9/PSME1/PSME2/PSMB10/PSMA4/PSMA1/PSMA6/UBB/PSMD11/PSMB7/CASP8/UBC/PSMA3/PSMA2/NOD1/ALPK1/SAA1/PSMC4/USP18/IL1R2/PSMA5/PSMB2/IKBKG/PSMA7/TRAF6/PSMD3/PSMC1/PSMF1/PSMB4/APP/PSMB1/RIPK2/PSMB3/MYD88/RELA/IL1RAP/PSMD13/PSMC6/PSMB6/PSMD7/IRAK1/PSMD12/PSMD14/MAP3K8/PSMB5/PELI3/HMGB1/PSMD1/NKIRAS1/SQSTM1/NLRX1/PSME3/PSMC2/MAP3K3/PSMD2/PSMD6/IL1RN/PSMC3/RBX1/IKBKB/PSMD5/UBE2N/PSMD9/NKIRAS2/PSMD4/NFKB1/CHUK/TRAF2/LRRC14/MAP2K4/IL1A/MAP2K6/USP14/SKP1/IL1B/FBXW11/PSMC5/PSMD8/TAB3/UBA52/TAB2/MAP3K7/RPS27A/TOLLIP/IKBIP/TAB1/PELI2 |
| R-HSA-9010553 | Regulation of expression of SLITs and ROBOs | 122/4886 | 172/11146 | 0.70930 | 1.61807 | 7.21686 | 4.14E-13 | 4.47E-11 | PSMB8/PSMB9/PSME1/PSME2/PSMB10/PSMA4/PSMA1/PSMA6/UBB/PSMD11/PSMB7/UBC/PSMA3/PSMA2/SMAD3/PSMC4/TEAD2/PSMA5/PSMB2/PSMA7/PSMD3/PSMC1/PSMF1/PSMB4/PSMB1/CDKN1A/CDKN2A/TGFB1/PSMB3/BRD2/MDM2/MYC/PSMD13/PSMC6/PSMB6/PSMD7/SNW1/PSMD12/PSMD14/PSMB5/TCF7L1/PSMD1/RUNX1/RBPJ/YAP1/PSME3/FOXO3/PSMC2/PSMD2/PSMD6/PSMC3/ZFHX3/SMAD4/JAG1/MAML3/PSMD5/CCND1/TCF7L2/PSMD9/PSMD4/KAT2B/TEAD1/KRAS/SRC/BCL2L11/PSMC5/CBFB/PSMD8/UBA52/MAML1/RUNX3/TEAD4/MAML2/HES1/RPS27A/NOTCH1/EP300 |
| R-HSA-9604323 | Negative regulation of NOTCH4 signaling | 49/4886 | 54/11146 | 0.90741 | 2.06999 | 6.96307 | 4.81E-13 | 4.87E-11 | PSMB8/PSMB9/PSME1/PSME2/PSMB10/PSMA4/PSMA1/PSMA6/UBB/PSMD11/PSMB7/UBC/PSMA3/PSMA2/PSMC4/CUL2/PSMA5/PSMB2/PSMA7/PSMD3/PSMC1/PSMF1/PSMB4/PSMB1/MSI1/PSMB3/RPL3/PSMD13/PSMC6/RPS6/PSMB6/PSMD7/PSMD12/ETF1/PSMD14/RPL10A/PSMB5/RPL13A/RPL5/RPS3A/USP33/PSMD1/EIF4G1/RPL15/RPL7/RPL23/RPL31/RPL12/DAG1/UPF2/RPS3/PSME3/PSMC2/NCBP2/PSMD2/RPS4X/RPL26/RPL34/RPL7A/PSMD6/RPL6/PSMC3/RPS27/LDB1/RPL22/RPL30/RPL4/EIF4A3/RPL11/RPS9/RBX1/RPS15A/RPS7/RPS14/RPS8/RPL14/RPL10/RPLP0/PSMD5/GSPT2/RPL37A/PSMD9/RPL41/UPF3A/PSMD4/RPL39/ROBO3/RPL9/PABPC1/RPL35A/RPL8/RPL32/RPL36AL/RPL26L1/RPS23/RPS28/RPS15/RPL37/RPLP1/RPL24/RPS25/MAGOH/RPL13/RPS24/RPL18/RPL36/RPS10/RBM8A/LHX4/PSMC5/PSMD8/UBA52/RNPS1/HOXA2/RPS16/RPL27A/RPS5/FAU/RPS2/RPS27A/RPL29/RPL38 |
| R-HSA-376176 | Signaling by ROBO receptors | 148/4886 | 219/11146 | 0.67580 | 1.54164 | 7.15175 | 7.28E-13 | 6.94E-11 | PSMB8/PSMB9/PSME1/PSME2/PSMB10/PSMA4/PSMA1/PSMA6/UBB/PSMD11/PSMB7/UBC/PSMA3/PSMA2/PSMC4/PSMA5/PSMB2/PSMA7/PSMD3/PSMC1/PSMF1/PSMB4/PSMB1/PSMB3/PSMD13/PSMC6/PSMB6/PSMD7/PSMD12/PSMD14/PSMB5/PSMD1/PSME3/PSMC2/PSMD2/PSMD6/PSMC3/RBX1/NOTCH4/FBXW7/PSMD5/PSMD9/PSMD4/YWHAZ/SKP1/PSMC5/PSMD8/UBA52/RPS27A |
| R-HSA-109581 | Apoptosis | 126/4886 | 180/11146 | 0.70000 | 1.59685 | 7.13191 | 7.95E-13 | 7.16E-11 | PSMB8/PSMB9/PSME1/PSME2/PSMB10/PSMA4/PSMA1/PSMA6/UBB/PSMD11/PSMB7/SRGAP2/UBC/PSMA3/PSMA2/PSMC4/CUL2/PSMA5/PSMB2/PSMA7/PSMD3/PSMC1/PSMF1/PSMB4/GPC1/PSMB1/MSI1/PSMB3/RPL3/FLRT3/PSMD13/PSMC6/RPS6/PSMB6/PSMD7/PSMD12/PRKACB/ETF1/PSMD14/RPL10A/PSMB5/RPL13A/RPL5/RPS3A/USP33/PSMD1/EIF4G1/NELL2/RPL15/RPL7/RPL23/RPL31/CXCR4/ENAH/RPL12/DAG1/UPF2/SOS1/RPS3/PAK1/PSME3/AKAP5/PSMC2/NCBP2/PAK6/PSMD2/RPS4X/PFN2/RPL26/SRGAP1/RPL34/RPL7A/PSMD6/RPL6/PFN1/PSMC3/RPS27/LDB1/RPL22/RPL30/RPL4/EIF4A3/RPL11/RPS9/RBX1/RPS15A/RPS7/RPS14/PAK2/RPS8/RPL14/RPL10/DCC/RPLP0/PSMD5/GSPT2/RPL37A/PSMD9/RPL41/EVL/UPF3A/PSMD4/RPL39/ROBO3/CAP2/RPL9/PABPC1/RPL35A/RPL8/RPL32/RPL36AL/RPL26L1/RPS23/RPS28/RPS15/RPL37/RPLP1/RPL24/RPS25/MAGOH/RPL13/RPS24/RPL18/SRC/PRKCA/RPL36/CAP1/RPS10/RBM8A/LHX4/PSMC5/PSMD8/UBA52/ABL2/RNPS1/HOXA2/RPS16/RPL27A/RPS5/VASP/NRP1/FAU/PPP3CB/RPS2/RPS27A/RPL29/PRKACA/RPL38 |
| R-HSA-1168372 | Downstream signaling events of B Cell Receptor (BCR) | 68/4886 | 83/11146 | 0.81928 | 1.86894 | 7.01979 | 9.38E-13 | 8.00E-11 | PSMB8/PSMB9/PSME1/PSME2/PSMB10/BAK1/PSMA4/PSMA1/PSMA6/GSDMD/UBB/PSMD11/PSMB7/PMAIP1/CASP8/UBC/PSMA3/TNFSF10/DYNLL2/PSMA2/CASP7/PSMC4/DAPK1/STAT3/PSMA5/PSMB2/TICAM1/PSMA7/CASP3/PSMD3/FAS/PSMC1/PSMF1/PSMB4/GSN/PPP3CC/PSMB1/TP63/CDKN2A/NMT1/PSMB3/AVEN/CYCS/PSMD13/PSMC6/PSMB6/PSMD7/STK24/RIPK1/PSMD12/UACA/PSMD14/PSMB5/CFLAR/HMGB1/PSMD1/PTK2/MAGED1/PLEC/YWHAB/CASP6/ARHGAP10/BMF/TFDP2/BAD/APPL1/DAPK2/TJP2/PSME3/C1QBP/YWHAQ/PSMC2/PPP3R1/DNM1L/KPNA1/CASP9/PSMD2/TLR4/SATB1/BID/PSMD6/CARD8/CLSPN/PSMC3/BIRC2/OCLN/PAK2/E2F1/KPNB1/DCC/TP73/PSMD5/HMGB2/DAPK3/FADD/PSMD9/PSMD4/UNC5B/DIABLO/TRAF2/LMNB1/YWHAZ/XIAP/FNTA/SFN/TFDP1/TNFRSF10B/BBC3/DBNL/ROCK1/BCL2L11/ADD1/YWHAG/OMA1/PSMC5/PSMD8/UNC5A/MAPK8/UBA52/PPP1R13B/OPA1/PRKCQ/BCAP31/RPS27A/APIP/DFFB |
| R-HSA-169911 | Regulation of Apoptosis | 48/4886 | 53/11146 | 0.90566 | 2.06600 | 6.87229 | 1.00E-12 | 8.11E-11 | PSMB8/PSMB9/PSME1/PSME2/PSMB10/PSMA4/PSMA1/PSMA6/UBB/RASGRP3/PSMD11/PSMB7/CALM3/UBC/PSMA3/PSMA2/PSMC4/PSMA5/PSMB2/IKBKG/PSMA7/PSMD3/PSMC1/PSMF1/PSMB4/PSMB1/NFKBIE/PSMB3/RELA/PSMD13/PSMC6/PSMB6/PSMD7/PSMD12/PSMD14/PSMB5/NFATC2/PSMD1/BCL10/PSME3/PSMC2/PPP3R1/PSMD2/RASGRP1/PSMD6/PSMC3/NFATC1/IKBKB/PSMD5/MALT1/PSMD9/PSMD4/NFKB1/CHUK/PPP3CA/KRAS/CALM1/SKP1/FBXW11/REL/PSMC5/FKBP1A/PSMD8/HRAS/UBA52/PPP3CB/MAP3K7/RPS27A |

2. WikiPathways pathway analysis

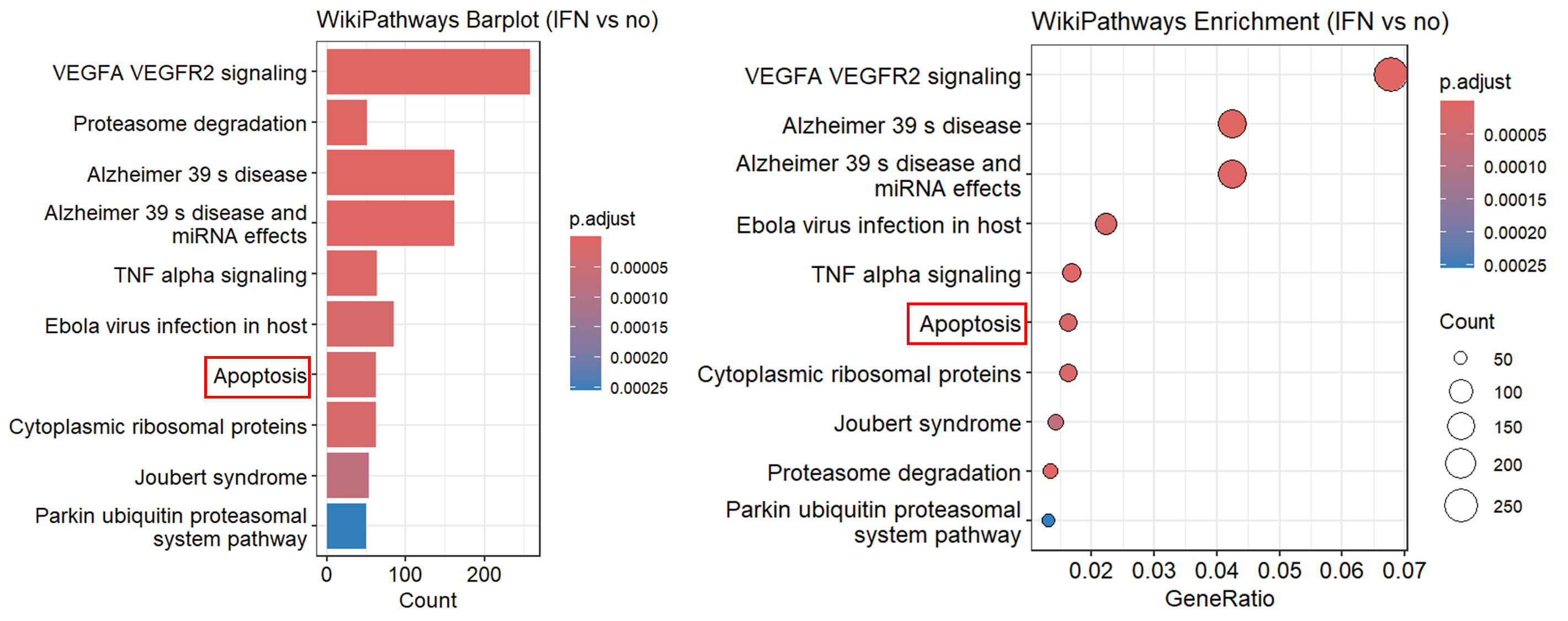

Top 20 WikiPathways signaling

| **ID** | **Description** | **Gene**  **Ratio** | **Bg Ratio** | **Rich**  **Factor** | **Fold**  **Enrichment** | **z-Score** | **p-value** | **p.**  **adjust** | **Gene ID** |
| --- | --- | --- | --- | --- | --- | --- | --- | --- | --- |
| WP3888 | VEGFA VEGFR2 signaling | 258/ 3807 | 432/ 8776 | 0.59722 | 1.37673 | 7.02860 | 1.92E-12 | 1.42E-09 | 6772/3383/4343/5717/3315/3301/57326/25865/3693/160/6774/6648/7170/8775/355/1666/6347/2817/5606/1267/3303/9325/2035/79094/114789/5335/170506/5954/9034/54205/4193/5970/154796/5515/3685/22820/55854/9564/6194/8650/84790/4646/9114/4673/9734/55379/4736/23521/11080/6125/8718/5329/6778/3146/2534/81873/4773/326624/1981/5747/51574/1796/6748/1827/10628/80031/6129/9513/1200/5034/8569/23215/6624/5605/781/2926/1397/4189/3725/1445/2746/5058/4929/2309/5601/3748/9146/7295/4641/8665/8754/5908/9252/84335/5562/2321/10758/8828/9588/32/1847/6154/857/2185/2113/2889/5228/5563/8027/2060/9365/79109/2130/219931/124491/10645/529/7430/9578/56905/7114/1385/1213/682/6729/5216/1398/6464/9451/5743/8575/4772/6902/4217/6747/182/4855/102/811/9261/100506658/7414/8867/5328/27289/10397/5062/468/5757/9645/813/4860/5295/5331/23476/595/8772/10525/1465/31/5710/6541/5226/7001/9209/11124/3688/4790/29109/1969/2308/253260/10963/23196/389/51278/5530/653361/1536/9344/26986/10818/5868/5829/8813/23597/2475/23753/23339/1982/9659/6416/274/5608/2081/382/409/10574/23189/6386/2152/56907/10061/3164/5780/867/8936/1977/5590/10056/8550/441478/8974/6093/2317/2876/23291/6714/5578/650/54505/54567/3320/4323/81567/5599/3265/4313/22926/2048/3490/7410/9100/154/7004/3418/4691/6418/8821/10014/4303/7086/29924/3939/83442/824/3486/7024/4633/56999/4627/51763/83473/4952/332/5170/5609/26058 |
| WP183 | Proteasome degradation | 51/ 3807 | 62/ 8776 | 0.82258 | 1.89624 | 6.19854 | 3.22E-10 | 1.19E-07 | 5696/5698/3133/5720/5721/5699/3105/3107/3106/5685/5682/3134/5687/7314/5717/5695/7316/5684/5683/5704/5686/5690/5688/7318/5709/5700/5692/7323/5689/5691/5719/5706/5694/5713/5718/5693/5707/10197/5701/3135/5708/9861/5702/7347/7322/5711/5715/5710/5705/5714/7317 |
| WP5124 | Alzheimer 39 s disease | 162/ 3807 | 263/ 8776 | 0.61597 | 1.41995 | 6.05222 | 1.26E-09 | 3.10E-07 | 5685/5682/5687/489/9776/5695/808/841/5684/7846/5683/83464/5610/1435/840/3667/5704/4843/5686/5690/5688/1454/836/5709/355/5700/5692/5533/351/1856/5689/823/55851/5691/54205/9706/5970/7417/5289/5719/776/5706/5694/4035/322/5713/27121/84790/5718/8324/7476/10213/5693/23621/54361/11047/5707/5664/91860/7280/3643/7976/51107/5605/572/5293/7124/5332/22863/5701/5534/5601/8408/842/5708/3831/8323/637/1855/1457/4040/9861/1452/7477/5702/8503/9451/292/5291/5743/1649/4217/1965/291/102/7473/7416/7132/468/64837/8313/3551/23385/5295/5331/84617/8772/5715/5663/3709/10383/3028/29982/5710/7494/4790/203068/1147/7186/5530/1536/6622/2475/55626/3552/2081/3845/801/23130/2906/10381/7419/2902/10297/3553/1460/80326/89953/8660/7481/5705/5714/369/5599/3710/3265/22926/4311/3708/10105/7855/348/55102/6868/3799/824/5532/79861/3416/488/2535/5609 |
| WP2059 | Alzheimer 39 s disease and miRNA effects | 162/ 3807 | 268/ 8776 | 0.60448 | 1.39346 | 5.72576 | 8.77E-09 | 1.62E-06 | 5685/5682/5687/489/9776/5695/808/841/5684/7846/5683/83464/5610/1435/840/3667/5704/4843/5686/5690/5688/1454/836/5709/355/5700/5692/5533/351/1856/5689/823/55851/5691/54205/9706/5970/7417/5289/5719/776/5706/5694/4035/322/5713/27121/84790/5718/8324/7476/10213/5693/23621/54361/11047/5707/5664/91860/7280/3643/7976/51107/5605/572/5293/7124/5332/22863/5701/5534/5601/8408/842/5708/3831/8323/637/1855/1457/4040/9861/1452/7477/5702/8503/9451/292/5291/5743/1649/4217/1965/291/102/7473/7416/7132/468/64837/8313/3551/23385/5295/5331/84617/8772/5715/5663/3709/10383/3028/29982/5710/7494/4790/203068/1147/7186/5530/1536/6622/2475/55626/3552/2081/3845/801/23130/2906/10381/7419/2902/10297/3553/1460/80326/89953/8660/7481/5705/5714/369/5599/3710/3265/22926/4311/3708/10105/7855/348/55102/6868/3799/824/5532/79861/3416/488/2535/5609 |
| WP231 | TNF alpha signaling | 64/ 3807 | 89/ 8776 | 0.71910 | 1.65769 | 5.45837 | 4.14E-08 | 6.13E-06 | 2752/841/330/7133/840/8517/836/4794/5606/29110/5515/8737/1326/8837/10131/6654/572/3725/7124/5790/5601/7295/842/4215/5708/9020/637/1457/11140/7128/4217/329/10010/7132/3551/6610/8772/8567/4214/4790/1147/56616/7186/5836/6416/5608/3845/8844/5590/6500/23291/11035/5966/3320/5599/257397/3265/56957/23118/117584/1535/6885/10454/5609 |
| WP4217 | Ebola virus infection in host | 85/ 3807 | 129/ 8776 | 0.65891 | 1.51895 | 5.19717 | 1.76E-07 | 2.11E-05 | 3133/6772/3105/3107/3108/3106/3134/3122/3115/3113/3480/3112/3109/7301/5610/3123/684/4240/2934/153090/29110/7150/65082/5970/7251/3685/27183/2348/57617/3672/55823/3119/7020/5293/2621/5058/708/3135/3111/3836/2318/7099/857/1212/2060/1211/219931/1213/8503/5291/8575/3655/9021/27072/1965/2316/3675/3127/3678/3661/5295/3117/3688/4790/3665/3384/558/389/3385/9367/5868/23339/60/7879/858/5922/2317/5966/4864/7410/8218/3120/6868/3118/2033 |
| WP254 | Apoptosis | 62/ 3807 | 88/ 8776 | 0.70455 | 1.62414 | 5.15041 | 2.27E-07 | 2.11E-05 | 3659/834/578/843/5366/841/3480/8743/330/4170/7133/840/8517/3660/836/355/4794/8626/1029/3663/54205/4193/5970/4609/8737/8837/8718/835/665/839/572/3725/7124/27242/842/637/3664/599/3070/329/837/7132/3551/3661/5295/7161/8772/4214/4790/3665/1147/56616/7186/331/666/6416/8795/27113/10018/9169/332/1677 |
| WP477 | Cytoplasmic ribosomal proteins | 62/ 3807 | 88/ 8776 | 0.70455 | 1.62414 | 5.15041 | 2.27E-07 | 2.11E-05 | 6122/6194/4736/23521/6125/6189/6138/6195/6129/9349/6160/6136/6188/6191/6154/6164/6130/6128/6232/6146/6156/6124/6135/6203/6210/6201/9801/6208/6202/9045/6134/6175/6168/6171/6196/6199/6170/6133/6165/6132/6161/6228/6234/6209/6167/6176/6152/6230/6137/6229/6141/25873/6204/7311/6217/6157/6193/2197/6187/6233/6159/6169 |
| WP4656 | Joubert syndrome | 54/ 3807 | 76/ 8776 | 0.71053 | 1.63792 | 4.88874 | 8.91E-07 | 7.33E-05 | 403/5108/57545/6469/79867/203286/582/5147/79600/26123/6102/4646/5158/51199/4644/4867/27077/1069/261734/55212/27241/1855/117177/9094/200728/5116/9738/2316/51684/4218/468/585/472/23322/142/80210/27130/402/51259/91147/2475/9731/129880/56623/10524/79848/22897/200894/10111/95681/80184/27031/65062/123016 |
| WP2359 | Parkin ubiquitin proteasomal system pathway | 50/ 3807 | 71/ 8776 | 0.70423 | 1.62340 | 4.61628 | 3.44E-06 | 2.55E-04 | 834/9246/5717/841/7846/5704/5709/5700/3306/3313/3303/5719/5706/5713/84790/5718/3308/10213/5707/7280/5701/118424/3304/5708/7332/3305/9861/7326/5702/10273/51182/55294/5711/84617/5715/25897/10383/5710/203068/6622/10381/5705/5714/7317/3309/3312/79861/8573/898/6477 |
| WP2004 | miR targeted genes in lymphocytes | 229/ 3807 | 424/ 8776 | 0.54009 | 1.24504 | 4.52695 | 4.21E-06 | 2.67E-04 | 79751/103/53/64866/4343/26064/7846/58533/4170/3667/160/51479/8943/7170/10241/57590/347902/3918/8546/9341/2580/1075/1026/1029/79665/5955/10423/27250/64422/2247/283237/9019/51616/10797/571/5887/253558/27230/3783/55503/84705/114793/1326/23621/5937/1871/8202/3475/2589/8774/2771/5878/10802/9513/7298/4552/781/3915/255520/51715/708/79969/10971/5236/2997/5534/7296/10672/9528/83858/1051/8161/3839/6675/5870/7127/51654/54964/5874/10449/391/57678/85440/27236/822/24145/29901/5925/55717/6902/79073/51014/5527/55660/3992/117145/7171/9497/7168/2820/83451/56262/4924/10036/56655/57532/124540/1284/23129/1869/372/5757/9354/285598/5479/4860/10270/1371/55225/595/8683/23224/4643/9076/8636/23589/8036/1465/10857/4774/8649/10282/51673/2673/1788/10484/3784/10611/55722/51125/558/4853/23423/5792/22862/64359/94081/83440/6509/23378/134266/5530/5725/29128/604/90522/80005/1027/23339/4144/84061/2948/4233/5510/8795/6386/7798/9522/3845/55347/5270/546/10061/10634/29099/4580/2483/1977/8801/594/84188/80335/23471/8974/9526/8744/23291/2539/6560/25828/10487/26973/64951/6576/3202/9319/865/29116/10558/23761/9296/27314/57552/9212/79602/9415/8301/6558/60559/4613/290/4691/10105/8487/83442/8829/4651/23647/5139/92935/79188/3159/488/8763/27243/54901/4851/8243/81034 |
| WP4197 | Immune response to tuberculosis | 20/ 3807 | 22/ 8776 | 0.90909 | 2.09566 | 4.50365 | 4.33E-06 | 2.67E-04 | 6890/5696/3659/6772/3437/8519/3430/10379/3717/6773/4599/8651/3434/3454/3455/4938/3460/3459/3716/5771 |
| WP5234 | Bardet Biedl syndrome | 59/ 3807 | 88/ 8776 | 0.67045 | 1.54555 | 4.50191 | 5.83E-06 | 3.32E-04 | 403/5311/9742/114327/23059/582/5147/6608/57560/167691/6102/23432/79659/51098/28981/1259/55081/54585/84314/5314/2736/79809/5727/51715/92482/55212/80258/27241/112752/11020/51684/57728/51626/374654/8239/4750/585/57539/80173/27130/55764/49855/26146/51259/91147/9731/129880/56623/22897/200894/22954/132320/201163/5310/95681/80184/27031/4952/123016 |
| WP1772 | Apoptosis modulation and signaling | 61/ 3807 | 92/ 8776 | 0.66304 | 1.52847 | 4.45997 | 7.05E-06 | 3.73E-04 | 834/578/843/8793/5366/841/9131/8743/330/4170/7133/840/7850/7189/836/355/8682/1029/3303/54205/4615/3654/8737/8837/8718/835/839/9531/90427/572/3725/842/9020/637/826/599/638/4217/329/837/7132/2021/3551/8772/4671/8567/5783/4790/56616/331/666/8795/27113/84883/10018/2353/5599/51651/54472/332/1677 |
| WP45 | G1 to S cell cycle control | 45/ 3807 | 64/ 8776 | 0.70313 | 1.62086 | 4.36322 | 1.13E-05 | 5.60E-04 | 1026/1017/1029/148327/4193/4609/896/7465/6117/4176/894/1871/1032/90993/7029/902/4174/993/1388/5427/1385/5925/1869/1022/10488/595/5426/472/84699/1019/4999/1647/4171/1027/8318/9134/7027/5001/4173/1870/1031/5000/1030/898/4331 |
| WP5039 | SARS CoV 2 innate immunity evasion and cell specific immune response | 46/ 3807 | 66/ 8776 | 0.69697 | 1.60667 | 4.33010 | 1.31E-05 | 5.76E-04 | 6772/8519/6373/6773/4599/841/3433/23367/4088/7706/79132/7189/6347/2920/29110/7040/7188/6352/8737/3454/7020/4283/3627/7124/9540/2921/3455/5473/682/3570/4928/6372/8480/10010/3066/164/3661/8772/4790/3665/3716/7186/59272/1437/57506/2033 |
| WP2003 | miR targeted genes in leukocytes | 79/ 3807 | 127/ 8776 | 0.62205 | 1.43396 | 4.31171 | 1.32E-05 | 5.76E-04 | 103/53/64866/7846/58533/4170/8943/7170/57590/2580/1075/10423/10797/3783/114793/1326/3475/2771/5878/10802/7298/10971/2997/7296/10672/1051/5870/7127/51654/54964/10449/7057/391/822/771/29901/55717/6902/79073/55660/3992/23657/56655/23129/372/5479/4860/8036/1465/8649/10611/51125/23423/94081/83440/6509/604/23339/5270/2483/23471/9526/8744/2539/25828/10487/26973/6576/10558/23761/9296/79602/290/10105/83442/79188/3159/488/8763 |
| WP179 | Cell cycle | 75/ 3807 | 120/ 8776 | 0.62500 | 1.44077 | 4.25522 | 1.71E-05 | 7.03E-04 | 994/4088/1875/1026/1017/1029/7040/4193/4609/10459/896/996/7465/1111/4176/894/1871/1032/7529/5933/5934/7029/902/10971/4174/7043/8379/7042/993/5925/8881/4089/29882/9978/1874/1869/3066/1022/595/472/1019/4999/1647/7272/4171/7534/545/2810/1027/8318/10393/25847/9134/7027/5001/4173/1870/6500/1031/7532/11200/990/699/5000/9700/6502/8556/51433/9184/995/7709/1030/2033/8243/898 |
| WP4666 | Hepatitis B infection | 91/ 3807 | 151/ 8776 | 0.60265 | 1.38924 | 4.22289 | 1.90E-05 | 7.20E-04 | 6772/843/3717/6773/64135/841/7046/6777/4088/6774/8517/148022/5582/7189/836/355/7098/1026/5606/29110/148327/7040/54205/4615/5970/4609/3654/537/3339/6778/4773/3454/7529/90993/5605/6654/572/3725/5293/7124/10971/5601/7043/842/7042/7099/1388/637/2185/6776/1385/8503/4776/5291/4772/4089/5600/468/10488/3551/3661/5295/8772/84699/4214/1643/4790/3665/1147/3716/7534/6416/5608/3845/9586/7419/6714/5578/369/2353/5599/3265/114609/23118/1642/57506/6885/332/10454/2033/5609 |
| WP1984 | Integrated breast cancer pathway | 92/ 3807 | 153/ 8776 | 0.60131 | 1.38615 | 4.21748 | 1.94E-05 | 7.20E-04 | 994/6772/578/4149/25780/5371/9958/841/7046/3667/80271/3612/4091/836/84640/1017/4092/60489/64781/4193/4609/7465/571/2956/1111/55526/5898/23077/8202/9175/2664/27005/1605/572/3725/57820/5058/10557/10600/842/993/637/6667/140739/580/1385/5925/641/4089/4436/5284/7249/1869/6790/1022/5888/7248/595/472/79877/8772/8061/1019/4664/4790/2308/1147/1647/3716/2355/2475/545/3845/659/23613/7035/26284/253959/4953/23411/11200/6597/369/10111/657/8846/79902/3156/466/10454/2033/8438 |

3. PathBank pathway analysis

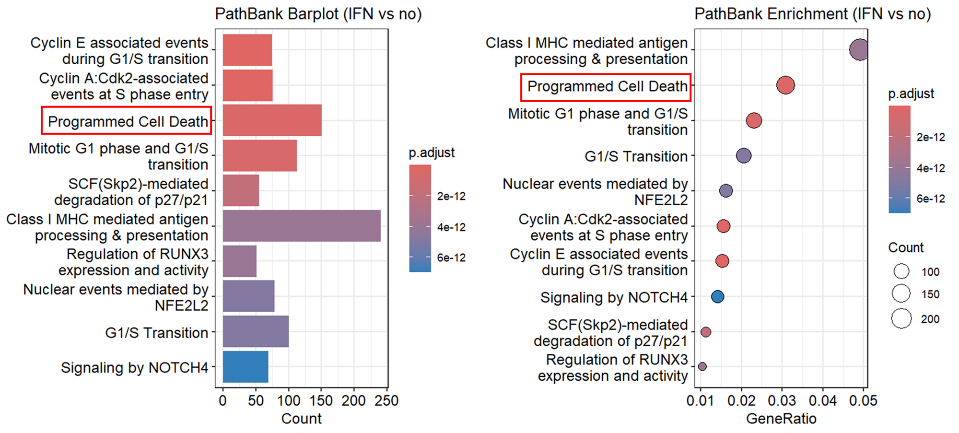

Top 20 PathBank pathway

| **ID** | **Description** | **Gene**  **Ratio** | **Bg Ratio** | **Rich Factor** | **Fold Enrichment** | **z-Score** | **p-value** | **p. adjust** | **Gene ID** |
| --- | --- | --- | --- | --- | --- | --- | --- | --- | --- |
| R-HSA-69202 | Cyclin E associated events during G1/S transition | 75/ 4886 | 83/ 11146 | 0.90361 | 2.06134 | 8.57403 | 4.45E-19 | 7.21E-16 | PSMB8/PSMB9/PSME1/PSME2/PSMB10/PSMA4/PSMA1/PSMA6/MAX/UBB/PSMD11/PSMB7/UBC/PSMA3/PSMA2/LIN52/PSMC4/PSMA5/E2F5/PSMB2/PSMA7/PSMD3/PSMC1/PSMF1/PSMB4/PSMB1/CDKN1A/CDK2/CABLES1/PSMB3/MYC/PSMD13/PSMC6/WEE1/PSMB6/PSMD7/RBBP4/PSMD12/PSMD14/PSMB5/PSMD1/RBL2/TFDP2/CCNH/PSME3/PSMC2/PSMD2/CDC25A/PSMD6/PSMC3/RB1/E2F4/E2F1/CDK7/PSMD5/CCND1/PSMD9/CDK4/PSMD4/LIN37/LIN9/PTK6/CDKN1B/CCNE2/TFDP1/LIN54/SKP1/PSMC5/PSMD8/UBA52/SKP2/RPS27A/CKS1B/CCNE1/MNAT1 |
| R-HSA-69656 | Cyclin A:Cdk2-associated events at S phase entry | 76/ 4886 | 85/ 11146 | 0.89412 | 2.03967 | 8.50037 | 1.15E-18 | 9.34E-16 | PSMB8/PSMB9/PSME1/CDC25B/PSME2/PSMB10/PSMA4/PSMA1/PSMA6/MAX/UBB/PSMD11/PSMB7/UBC/PSMA3/PSMA2/LIN52/PSMC4/PSMA5/E2F5/PSMB2/PSMA7/PSMD3/PSMC1/PSMF1/PSMB4/PSMB1/CDKN1A/CDK2/CABLES1/PSMB3/MYC/PSMD13/PSMC6/WEE1/PSMB6/PSMD7/RBBP4/PSMD12/PSMD14/PSMB5/PSMD1/RBL2/TFDP2/CCNH/PSME3/PSMC2/PSMD2/CDC25A/PSMD6/PSMC3/RB1/E2F4/E2F1/CDK7/PSMD5/CCND1/PSMD9/CDK4/PSMD4/LIN37/LIN9/PTK6/CDKN1B/CCNE2/TFDP1/LIN54/SKP1/PSMC5/PSMD8/UBA52/SKP2/RPS27A/CKS1B/CCNE1/MNAT1 |
| R-HSA-5357801 | Programmed Cell Death | 151/ 4886 | 213/ 11146 | 0.70892 | 1.61720 | 8.03477 | 7.08E-16 | 3.82E-13 | PSMB8/PSMB9/PSME1/IRF1/PSME2/PSMB10/CASP1/BAK1/PSMA4/PSMA1/PSMA6/GSDMD/UBB/PSMD11/PSMB7/PMAIP1/CASP8/UBC/PSMA3/TNFSF10/DYNLL2/BIRC3/PSMA2/CASP7/PSMC4/DAPK1/MLKL/STAT3/PSMA5/PSMB2/TICAM1/PSMA7/IRF2/CASP3/PSMD3/FAS/PSMC1/PSMF1/PSMB4/GSN/PPP3CC/PSMB1/TP63/CDKN2A/NMT1/PSMB3/AVEN/CYCS/PSMD13/PSMC6/PSMB6/PSMD7/STK24/RIPK1/PSMD12/UACA/PSMD14/PSMB5/CFLAR/HMGB1/PSMD1/PTK2/MAGED1/PLEC/YWHAB/CASP5/CASP6/ARHGAP10/BMF/TFDP2/BAD/APPL1/DAPK2/TJP2/PSME3/C1QBP/YWHAQ/PSMC2/PPP3R1/DNM1L/KPNA1/CASP9/PSMD2/TLR4/UBE2L3/SATB1/BID/PSMD6/CARD8/CLSPN/PSMC3/CDC37/STUB1/BIRC2/CASP4/OCLN/PAK2/E2F1/CHMP4B/IL18/KPNB1/DCC/TP73/PSMD5/HMGB2/DAPK3/FADD/PSMD9/PSMD4/UNC5B/DIABLO/TRAF2/LMNB1/YWHAZ/XIAP/FNTA/SFN/CHMP4A/IL1A/CHMP3/TFDP1/TNFRSF10B/CHMP4C/SDCBP/BBC3/DBNL/IL1B/ROCK1/BCL2L11/RIPK3/ADD1/YWHAG/OMA1/PSMC5/HSP90AA1/PSMD8/UNC5A/MAPK8/UBA52/PPP1R13B/OPA1/PDCD6IP/CHMP6/CHMP7/PRKCQ/BCAP31/RPS27A/APIP/OGT/CHMP2A/DFFB |
| R-HSA-453279 | Mitotic G1 phase and G1/S transition | 113/ 4886 | 149/ 11146 | 0.75839 | 1.73005 | 7.92567 | 1.29E-15 | 5.21E-13 | PSMB8/PSMB9/PSME1/PSME2/PSMB10/PSMA4/PSMA1/PSMA6/JAK2/MAX/UBB/LYN/PSMD11/PSMB7/UBC/PSMA3/PSMA2/LIN52/PSMC4/PSMA5/E2F5/PSMB2/PSMA7/PSMD3/PSMC1/PSMF1/PSMB4/PSMB1/CDKN1A/CDK2/CDKN2A/CABLES1/PSMB3/MYC/PPP2CA/PSMD13/CCND3/PSMC6/WEE1/PSMB6/PSMD7/RPA1/RBBP4/PSMD12/PSMD14/PSMB5/GMNN/MCM7/CCND2/E2F3/PSMD1/CDKN2D/RBL1/TYMS/RBL2/TFDP2/PPP2CB/CCNH/PSME3/PSMC2/MCM5/PSMD2/CDC25A/DYRK1A/TK1/PSMD6/POLE2/MCM8/FBXO5/PSMC3/RB1/POLA1/POLE4/E2F4/E2F1/CDK7/PSMD5/CCND1/POLE/PSMD9/MYBL2/CDK4/PSMD4/ORC2/MCM10/LIN37/LIN9/MCM2/PTK6/CDKN1B/CDC45/CCNE2/TFDP1/ORC5/POLE3/MCM4/LIN54/E2F2/SKP1/SRC/CDKN2C/PSMC5/PSMD8/CDC6/UBA52/ORC4/SKP2/RRM2/RPS27A/CDKN2B/CKS1B/CCNE1/MNAT1 |
| R-HSA-187577 | SCF(Skp2)-mediated degradation of p27/p21 | 55/ 4886 | 60/ 11146 | 0.91667 | 2.09111 | 7.48663 | 5.65E-15 | 1.83E-12 | PSMB8/PSMB9/PSME1/PSME2/PSMB10/PSMA4/PSMA1/PSMA6/UBB/PSMD11/PSMB7/UBC/PSMA3/PSMA2/PSMC4/PSMA5/PSMB2/PSMA7/PSMD3/PSMC1/PSMF1/PSMB4/PSMB1/CDKN1A/CDK2/PSMB3/PSMD13/PSMC6/PSMB6/PSMD7/PSMD12/PSMD14/PSMB5/PSMD1/PSME3/PSMC2/PSMD2/PSMD6/PSMC3/PSMD5/CCND1/PSMD9/CDK4/PSMD4/PTK6/CDKN1B/CCNE2/SKP1/PSMC5/PSMD8/UBA52/SKP2/RPS27A/CKS1B/CCNE1 |
| R-HSA-983169 | Class I MHC mediated antigen processing & presentation | 240/ 4886 | 381/ 11146 | 0.62992 | 1.43698 | 7.66744 | 1.60E-14 | 3.97E-12 | TAP1/PSMB8/PSMB9/HLA-E/TAP2/PSME1/B2M/PSME2/PSMB10/HLA-A/UBE2L6/HLA-C/HLA-B/PSMA4/PSMA1/HLA-F/DTX3L/PSMA6/RNF213/CTSS/UBB/PSMD11/PSMB7/FBXO6/TRIM21/ERAP2/FBXO7/UBR2/TRIM69/UBC/PSMA3/RNF114/CUL7/PSMA2/KCTD7/UBA6/PSMC4/CUL2/ITGB5/RBCK1/PSMA5/UBOX5/PSMB2/IKBKG/RNF19B/HERC6/TAPBP/PSMA7/UBA7/UBE2Z/PSMD3/KLHL22/SEC24C/PSMC1/PSMF1/PSMB4/ASB1/UBE2D3/ERAP1/PDIA3/PSMB1/VAMP3/MKRN1/KLHL5/FBXO2/ZNRF1/PSMB3/STX4/MYD88/FBXL5/PIK3C3/PSMD13/SEC31A/ITGAV/FBXO31/CDC27/PSMC6/RNF130/PSMB6/SOCS1/PSMD7/RNF34/CUL3/KLHL3/PSMD12/VHL/PSMD14/LMO7/PSMB5/RNF115/UBE2E3/FBXO21/TRIM11/HMGB1/FBXO10/PSMD1/UBE2M/SEC24A/KLHL21/ASB7/UBE3B/VAMP8/HERC1/FBXO4/UBE2Q1/PSME3/UBE2V2/ASB14/ATG14/PSMC2/HLA-G/RNF123/RNF217/RNF126/TRIP12/UBE2J2/SEC61B/PSMD2/TLR4/UBE2L3/HERC4/UBE2Q2/UBE2F/RNF6/LONRF1/PSMD6/DZIP3/HERC5/KEAP1/UBE2G1/PSMC3/CBLB/TRAF7/STUB1/CDC16/UBE2E1/RNF7/UBE2O/SOCS3/ASB8/ANAPC2/HECTD3/KLHL2/RBX1/FBXO41/RNF4/CALR/UBE2D2/RLIM/LRSAM1/SEC22B/UBE4A/FBXW7/RNF144B/IKBKB/SEC24B/LRRC41/PSMD5/HACE1/ASB6/UBE2N/PSMD9/RNF19A/LNPEP/FBXO44/KLHL13/UNKL/KCTD6/FBXL18/PSMD4/TRIM37/UBE2E2/SEC23A/UBE3D/CHUK/FBXO9/TRAIP/BLMH/NCF1/CYBB/DET1/FBXW2/S100A1/ANAPC10/ANAPC13/CDC34/RNF14/RNF138/TRIM32/FBXO22/SEC13/RCHY1/FBXW10/SAR1B/THOP1/FBXL20/SKP1/UBE2K/KBTBD8/FBXW11/HERC3/UBA3/ATG7/SPSB1/FBXL3/FBXO32/PSMC5/FBXL12/PSMD8/FBXW5/MYLIP/WSB1/UBR4/RNF41/UBA52/ZNRF2/TIRAP/UBA1/SKP2/KLHL25/ANAPC5/HSPA5/MIB2/MGRN1/BCAP31/CYBA/CCNF/FBXW9/RPS27A/ARIH2/UBA5/FBXL22/FBXL19/FBXL15/FBXL13/FBXL14/CDC26/SIAH1/FBXL8/RNF25 |
| R-HSA-8941858 | Regulation of RUNX3 expression and activity | 51/ 4886 | 55/ 11146 | 0.92727 | 2.11531 | 7.32520 | 1.71E-14 | 3.97E-12 | PSMB8/PSMB9/PSME1/PSME2/PSMB10/PSMA4/PSMA1/PSMA6/UBB/PSMD11/PSMB7/UBC/PSMA3/PSMA2/PSMC4/PSMA5/PSMB2/PSMA7/PSMD3/PSMC1/PSMF1/PSMB4/PSMB1/CDKN2A/TGFB1/PSMB3/MDM2/PSMD13/PSMC6/PSMB6/PSMD7/PSMD12/PSMD14/PSMB5/PSMD1/PSME3/PSMC2/PSMD2/PSMD6/PSMC3/PSMD5/PSMD9/PSMD4/SRC/PSMC5/CBFB/PSMD8/UBA52/RUNX3/RPS27A/EP300 |
| R-HSA-9759194 | Nuclear events mediated by NFE2L2 | 79/ 4886 | 97/ 11146 | 0.81443 | 1.85789 | 7.49700 | 2.50E-14 | 4.87E-12 | PSMB8/PSMB9/PSME1/PSME2/PSMB10/PSMA4/PSMA1/PSMA6/UBB/PSMD11/PSMB7/UBC/PSMA3/PSMA2/PSMC4/PSMA5/PSMB2/PSMA7/GSTA1/PSMD3/PSMC1/PSMF1/PSMB4/CCL2/PSMB1/CDKN2A/AREG/PSMB3/RELA/MYC/PSMD13/PSMC6/BACH1/PSMB6/ABCC3/PSMD7/SRXN1/PSMD12/PSMD14/PSMB5/PSMD1/GCLM/SQSTM1/ME1/GCLC/PSME3/PSMC2/TXN/TXNRD1/PSMD2/SP1/PRKAA2/PSMD6/KEAP1/MAFG/PSMC3/TALDO1/RBX1/SLC7A11/ATF4/NQO1/PSMD5/PSMD9/PSMD4/PGD/NFKB1/ABCC1/ABCF2/GSR/SKP1/G6PD/PSMC5/PSMD8/UBA52/TKT/RPS27A/MAFK/NOTCH1/EP300 |
| R-HSA-69206 | G1/S Transition | 100/ 4886 | 131/ 11146 | 0.76336 | 1.74138 | 7.54077 | 2.71E-14 | 4.87E-12 | PSMB8/PSMB9/PSME1/PSME2/PSMB10/PSMA4/PSMA1/PSMA6/MAX/UBB/PSMD11/PSMB7/UBC/PSMA3/PSMA2/LIN52/PSMC4/PSMA5/E2F5/PSMB2/PSMA7/PSMD3/PSMC1/PSMF1/PSMB4/PSMB1/CDKN1A/CDK2/CABLES1/PSMB3/MYC/PPP2CA/PSMD13/PSMC6/WEE1/PSMB6/PSMD7/RPA1/RBBP4/PSMD12/PSMD14/PSMB5/GMNN/MCM7/PSMD1/RBL1/TYMS/RBL2/TFDP2/PPP2CB/CCNH/PSME3/PSMC2/MCM5/PSMD2/CDC25A/TK1/PSMD6/POLE2/MCM8/FBXO5/PSMC3/RB1/POLA1/POLE4/E2F4/E2F1/CDK7/PSMD5/CCND1/POLE/PSMD9/CDK4/PSMD4/ORC2/MCM10/LIN37/LIN9/MCM2/PTK6/CDKN1B/CDC45/CCNE2/TFDP1/ORC5/POLE3/MCM4/LIN54/SKP1/PSMC5/PSMD8/CDC6/UBA52/ORC4/SKP2/RRM2/RPS27A/CKS1B/CCNE1/MNAT1 |
| R-HSA-9013694 | Signaling by NOTCH4 | 69/ 4886 | 82/ 11146 | 0.84146 | 1.91956 | 7.38344 | 4.57E-14 | 6.99E-12 | PSMB8/PSMB9/PSME1/PSME2/PSMB10/PSMA4/PSMA1/PSMA6/UBB/PSMD11/PSMB7/UBC/PSMA3/PSMA2/APH1B/SMAD3/PSMC4/PSMA5/PSMB2/PSMA7/PSMD3/PSMC1/PSMF1/PSMB4/PSMB1/PSENEN/PSMB3/PSMD13/PSMC6/PSMB6/PSMD7/SNW1/PSMD12/PSMD14/PSMB5/PSMD1/PSEN2/RBPJ/APH1A/PSME3/PSMC2/PSMD2/PSMD6/PSMC3/JAG1/RBX1/NOTCH4/ADAM10/MAML3/FBXW7/NCSTN/PSMD5/PSMD9/PSEN1/PSMD4/NOTCH2/KAT2B/YWHAZ/SKP1/DLL4/PSMC5/PSMD8/UBA52/MAML1/MAML2/HES1/RPS27A/NOTCH1/EP300 |
| R-HSA-69242 | S Phase | 118/ 4886 | 162/ 11146 | 0.72840 | 1.66162 | 7.49407 | 4.74E-14 | 6.99E-12 | PSMB8/PSMB9/PSME1/CDC25B/PSME2/PSMB10/PSMA4/PSMA1/PSMA6/MAX/UBB/PSMD11/PSMB7/UBC/PSMA3/PSMA2/LIN52/PSMC4/PSMA5/E2F5/PSMB2/PSMA7/PSMD3/PSMC1/PSMF1/PSMB4/PSMB1/CDKN1A/CDK2/CABLES1/PSMB3/MYC/PSMD13/CDC27/PSMC6/WEE1/PSMB6/PSMD7/RPA1/RBBP4/PSMD12/PSMD14/PSMB5/GMNN/MCM7/PSMD1/RBL2/TFDP2/CCNH/PSME3/PSMC2/MCM5/RFC2/ANAPC16/POLD1/PSMD2/CDC25A/PDS5B/PSMD6/POLE2/LIG1/MCM8/PSMC3/RB1/CDC16/UBE2E1/POLA1/ANAPC2/GINS2/RBX1/FEN1/POLE4/E2F4/E2F1/CDK7/PSMD5/CCND1/POLE/PSMD9/CDK4/PSMD4/ORC2/RFC5/LIN37/LIN9/GINS4/POLD3/MCM2/PTK6/CDKN1B/CDC45/ANAPC10/CCNE2/TFDP1/ORC5/POLE3/ESCO2/MCM4/CDCA5/LIN54/RFC4/SKP1/RFC3/PSMC5/PSMD8/CDC6/UBA52/ORC4/GINS3/SKP2/ANAPC5/POLD4/RPS27A/CKS1B/CDC26/SMC1A/CCNE1/MNAT1 |
| R-HSA-450408 | AUF1 (hnRNP D0) binds and destabilizes mRNA | 51/ 4886 | 56/ 11146 | 0.91071 | 2.07753 | 7.14148 | 1.10E-13 | 1.49E-11 | PSMB8/PSMB9/PSME1/PSME2/PSMB10/PSMA4/PSMA1/PSMA6/UBB/PSMD11/PSMB7/HSPB1/UBC/PSMA3/PSMA2/PSMC4/PSMA5/PSMB2/PSMA7/PSMD3/PSMC1/PSMF1/PSMB4/PSMB1/HSPA1A/PSMB3/PSMD13/PSMC6/PSMB6/PSMD7/PSMD12/PSMD14/PSMB5/PSMD1/EIF4G1/PSME3/PSMC2/HSPA1B/PSMD2/PSMD6/PSMC3/PSMD5/PSMD9/PSMD4/PABPC1/PSMC5/PSMD8/UBA52/HSPA8/RPS27A/HNRNPD |
| R-HSA-9020702 | Interleukin-1 signaling | 89/ 4886 | 115/ 11146 | 0.77391 | 1.76546 | 7.28942 | 1.75E-13 | 2.18E-11 | PSMB8/NLRC5/PSMB9/PSME1/PSME2/PSMB10/PSMA4/PSMA1/PSMA6/UBB/PSMD11/PSMB7/CASP8/UBC/PSMA3/PSMA2/NOD1/ALPK1/SAA1/PSMC4/USP18/IL1R2/PSMA5/PSMB2/IKBKG/PSMA7/TRAF6/PSMD3/PSMC1/PSMF1/PSMB4/APP/PSMB1/RIPK2/PSMB3/MYD88/RELA/IL1RAP/PSMD13/PSMC6/PSMB6/PSMD7/IRAK1/PSMD12/PSMD14/MAP3K8/PSMB5/PELI3/HMGB1/PSMD1/NKIRAS1/SQSTM1/NLRX1/PSME3/PSMC2/MAP3K3/PSMD2/PSMD6/IL1RN/PSMC3/RBX1/IKBKB/PSMD5/UBE2N/PSMD9/NKIRAS2/PSMD4/NFKB1/CHUK/TRAF2/LRRC14/MAP2K4/IL1A/MAP2K6/USP14/SKP1/IL1B/FBXW11/PSMC5/PSMD8/TAB3/UBA52/TAB2/MAP3K7/RPS27A/TOLLIP/IKBIP/TAB1/PELI2 |
| R-HSA-8878159 | Transcriptional regulation by RUNX3 | 77/ 4886 | 96/ 11146 | 0.80208 | 1.82972 | 7.21301 | 2.53E-13 | 2.93E-11 | PSMB8/PSMB9/PSME1/PSME2/PSMB10/PSMA4/PSMA1/PSMA6/UBB/PSMD11/PSMB7/UBC/PSMA3/PSMA2/SMAD3/PSMC4/TEAD2/PSMA5/PSMB2/PSMA7/PSMD3/PSMC1/PSMF1/PSMB4/PSMB1/CDKN1A/CDKN2A/TGFB1/PSMB3/BRD2/MDM2/MYC/PSMD13/PSMC6/PSMB6/PSMD7/SNW1/PSMD12/PSMD14/PSMB5/TCF7L1/PSMD1/RUNX1/RBPJ/YAP1/PSME3/FOXO3/PSMC2/PSMD2/PSMD6/PSMC3/ZFHX3/SMAD4/JAG1/MAML3/PSMD5/CCND1/TCF7L2/PSMD9/PSMD4/KAT2B/TEAD1/KRAS/SRC/BCL2L11/PSMC5/CBFB/PSMD8/UBA52/MAML1/RUNX3/TEAD4/MAML2/HES1/RPS27A/NOTCH1/EP300 |
| R-HSA-9010553 | Regulation of expression of SLITs and ROBOs | 122/ 4886 | 172/ 11146 | 0.70930 | 1.61807 | 7.21686 | 4.14E-13 | 4.47E-11 | PSMB8/PSMB9/PSME1/PSME2/PSMB10/PSMA4/PSMA1/PSMA6/UBB/PSMD11/PSMB7/UBC/PSMA3/PSMA2/PSMC4/CUL2/PSMA5/PSMB2/PSMA7/PSMD3/PSMC1/PSMF1/PSMB4/PSMB1/MSI1/PSMB3/RPL3/PSMD13/PSMC6/RPS6/PSMB6/PSMD7/PSMD12/ETF1/PSMD14/RPL10A/PSMB5/RPL13A/RPL5/RPS3A/USP33/PSMD1/EIF4G1/RPL15/RPL7/RPL23/RPL31/RPL12/DAG1/UPF2/RPS3/PSME3/PSMC2/NCBP2/PSMD2/RPS4X/RPL26/RPL34/RPL7A/PSMD6/RPL6/PSMC3/RPS27/LDB1/RPL22/RPL30/RPL4/EIF4A3/RPL11/RPS9/RBX1/RPS15A/RPS7/RPS14/RPS8/RPL14/RPL10/RPLP0/PSMD5/GSPT2/RPL37A/PSMD9/RPL41/UPF3A/PSMD4/RPL39/ROBO3/RPL9/PABPC1/RPL35A/RPL8/RPL32/RPL36AL/RPL26L1/RPS23/RPS28/RPS15/RPL37/RPLP1/RPL24/RPS25/MAGOH/RPL13/RPS24/RPL18/RPL36/RPS10/RBM8A/LHX4/PSMC5/PSMD8/UBA52/RNPS1/HOXA2/RPS16/RPL27A/RPS5/FAU/RPS2/RPS27A/RPL29/RPL38 |
| R-HSA-9604323 | Negative regulation of NOTCH4 signaling | 49/ 4886 | 54/ 11146 | 0.90741 | 2.06999 | 6.96307 | 4.81E-13 | 4.87E-11 | PSMB8/PSMB9/PSME1/PSME2/PSMB10/PSMA4/PSMA1/PSMA6/UBB/PSMD11/PSMB7/UBC/PSMA3/PSMA2/PSMC4/PSMA5/PSMB2/PSMA7/PSMD3/PSMC1/PSMF1/PSMB4/PSMB1/PSMB3/PSMD13/PSMC6/PSMB6/PSMD7/PSMD12/PSMD14/PSMB5/PSMD1/PSME3/PSMC2/PSMD2/PSMD6/PSMC3/RBX1/NOTCH4/FBXW7/PSMD5/PSMD9/PSMD4/YWHAZ/SKP1/PSMC5/PSMD8/UBA52/RPS27A |
| R-HSA-376176 | Signaling by ROBO receptors | 148/ 4886 | 219/ 11146 | 0.67580 | 1.54164 | 7.15175 | 7.28E-13 | 6.94E-11 | PSMB8/PSMB9/PSME1/PSME2/PSMB10/PSMA4/PSMA1/PSMA6/UBB/PSMD11/PSMB7/SRGAP2/UBC/PSMA3/PSMA2/PSMC4/CUL2/PSMA5/PSMB2/PSMA7/PSMD3/PSMC1/PSMF1/PSMB4/GPC1/PSMB1/MSI1/PSMB3/RPL3/FLRT3/PSMD13/PSMC6/RPS6/PSMB6/PSMD7/PSMD12/PRKACB/ETF1/PSMD14/RPL10A/PSMB5/RPL13A/RPL5/RPS3A/USP33/PSMD1/EIF4G1/NELL2/RPL15/RPL7/RPL23/RPL31/CXCR4/ENAH/RPL12/DAG1/UPF2/SOS1/RPS3/PAK1/PSME3/AKAP5/PSMC2/NCBP2/PAK6/PSMD2/RPS4X/PFN2/RPL26/SRGAP1/RPL34/RPL7A/PSMD6/RPL6/PFN1/PSMC3/RPS27/LDB1/RPL22/RPL30/RPL4/EIF4A3/RPL11/RPS9/RBX1/RPS15A/RPS7/RPS14/PAK2/RPS8/RPL14/RPL10/DCC/RPLP0/PSMD5/GSPT2/RPL37A/PSMD9/RPL41/EVL/UPF3A/PSMD4/RPL39/ROBO3/CAP2/RPL9/PABPC1/RPL35A/RPL8/RPL32/RPL36AL/RPL26L1/RPS23/RPS28/RPS15/RPL37/RPLP1/RPL24/RPS25/MAGOH/RPL13/RPS24/RPL18/SRC/PRKCA/RPL36/CAP1/RPS10/RBM8A/LHX4/PSMC5/PSMD8/UBA52/ABL2/RNPS1/HOXA2/RPS16/RPL27A/RPS5/VASP/NRP1/FAU/PPP3CB/RPS2/RPS27A/RPL29/PRKACA/RPL38 |
| R-HSA-109581 | Apoptosis | 126/ 4886 | 180/ 11146 | 0.70000 | 1.59685 | 7.13191 | 7.95E-13 | 7.16E-11 | PSMB8/PSMB9/PSME1/PSME2/PSMB10/BAK1/PSMA4/PSMA1/PSMA6/GSDMD/UBB/PSMD11/PSMB7/PMAIP1/CASP8/UBC/PSMA3/TNFSF10/DYNLL2/PSMA2/CASP7/PSMC4/DAPK1/STAT3/PSMA5/PSMB2/TICAM1/PSMA7/CASP3/PSMD3/FAS/PSMC1/PSMF1/PSMB4/GSN/PPP3CC/PSMB1/TP63/CDKN2A/NMT1/PSMB3/AVEN/CYCS/PSMD13/PSMC6/PSMB6/PSMD7/STK24/RIPK1/PSMD12/UACA/PSMD14/PSMB5/CFLAR/HMGB1/PSMD1/PTK2/MAGED1/PLEC/YWHAB/CASP6/ARHGAP10/BMF/TFDP2/BAD/APPL1/DAPK2/TJP2/PSME3/C1QBP/YWHAQ/PSMC2/PPP3R1/DNM1L/KPNA1/CASP9/PSMD2/TLR4/SATB1/BID/PSMD6/CARD8/CLSPN/PSMC3/BIRC2/OCLN/PAK2/E2F1/KPNB1/DCC/TP73/PSMD5/HMGB2/DAPK3/FADD/PSMD9/PSMD4/UNC5B/DIABLO/TRAF2/LMNB1/YWHAZ/XIAP/FNTA/SFN/TFDP1/TNFRSF10B/BBC3/DBNL/ROCK1/BCL2L11/ADD1/YWHAG/OMA1/PSMC5/PSMD8/UNC5A/MAPK8/UBA52/PPP1R13B/OPA1/PRKCQ/BCAP31/RPS27A/APIP/DFFB |
| R-HSA-1168372 | Downstream signaling events of B Cell Receptor (BCR) | 68/ 4886 | 83/ 11146 | 0.81928 | 1.86894 | 7.01979 | 9.38E-13 | 8.00E-11 | PSMB8/PSMB9/PSME1/PSME2/PSMB10/PSMA4/PSMA1/PSMA6/UBB/RASGRP3/PSMD11/PSMB7/CALM3/UBC/PSMA3/PSMA2/PSMC4/PSMA5/PSMB2/IKBKG/PSMA7/PSMD3/PSMC1/PSMF1/PSMB4/PSMB1/NFKBIE/PSMB3/RELA/PSMD13/PSMC6/PSMB6/PSMD7/PSMD12/PSMD14/PSMB5/NFATC2/PSMD1/BCL10/PSME3/PSMC2/PPP3R1/PSMD2/RASGRP1/PSMD6/PSMC3/NFATC1/IKBKB/PSMD5/MALT1/PSMD9/PSMD4/NFKB1/CHUK/PPP3CA/KRAS/CALM1/SKP1/FBXW11/REL/PSMC5/FKBP1A/PSMD8/HRAS/UBA52/PPP3CB/MAP3K7/RPS27A |
| R-HSA-169911 | Regulation of Apoptosis | 48/ 4886 | 53/ 11146 | 0.90566 | 2.06600 | 6.87229 | 1.00E-12 | 8.11E-11 | PSMB8/PSMB9/PSME1/PSME2/PSMB10/PSMA4/PSMA1/PSMA6/UBB/PSMD11/PSMB7/UBC/PSMA3/PSMA2/PSMC4/PSMA5/PSMB2/PSMA7/PSMD3/PSMC1/PSMF1/PSMB4/PSMB1/PSMB3/PSMD13/PSMC6/PSMB6/PSMD7/PSMD12/PSMD14/PSMB5/PSMD1/ARHGAP10/PSME3/PSMC2/PSMD2/PSMD6/PSMC3/PAK2/PSMD5/PSMD9/PSMD4/OMA1/PSMC5/PSMD8/UBA52/OPA1/RPS27A |

4. KEGG pathway analysis

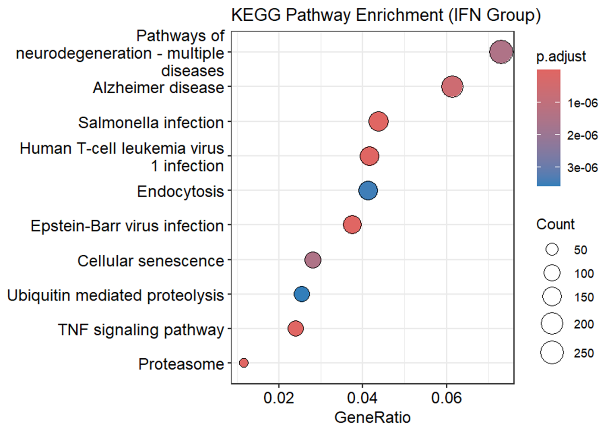

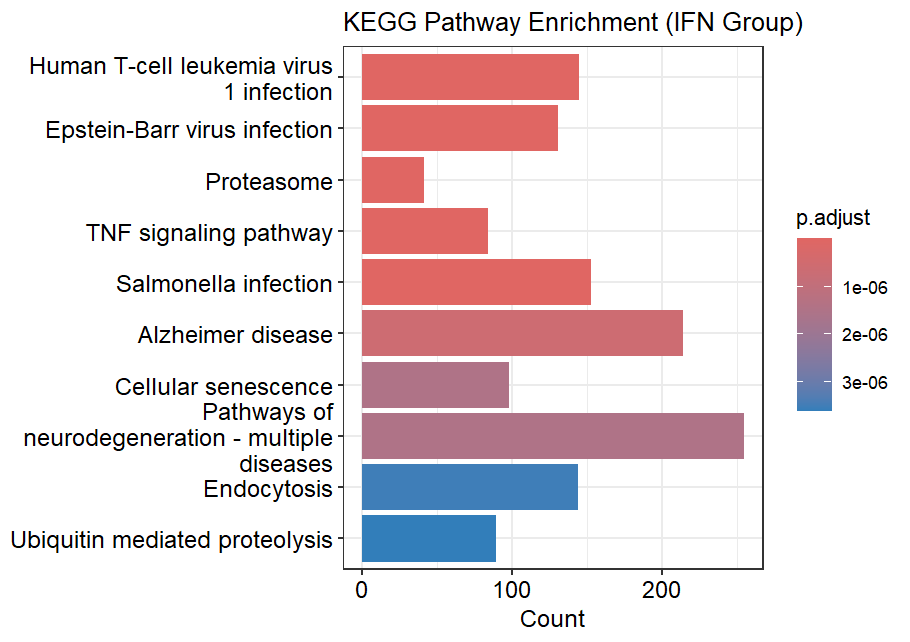

Top 20 KEGG pathway

| category | Sub-category | ID | Description | Gene  Ratio | Bg  Ratio | Rich actor | Fold Erichment | z-score | p-value | p. adjust | q-value | geneID |
| --- | --- | --- | --- | --- | --- | --- | --- | --- | --- | --- | --- | --- |
| Human Diseases | Infectious disease: viral | hsa05166 | Human T-cell leukemia virus 1 infection | 145/ 3489 | 224/ 8538 | 0.64732 | 1.58407 | 7.36354 | 2.53E-13 | 8.73E-11 | 5.56E-11 | HLA-E/B2M/IL15RA/HLA-A/HLA-C/HLA-DMA/HLA-B/HLA-F/ICAM1/HLA-DRA/HLA-DPB1/IL15/HLA-DPA1/HLA-DOB/TGFBR1/HLA-DMB/STAT5B/SMAD3/HLA-DRB1/IL1R2/IKBKG/PPP3CC/CDKN1A/CDK2/CDKN2A/CREB3L4/TGFB1/ETS2/ADCY3/TCF3/RELA/MYC/CD40/VDAC2/ITGB2/LCK/CCND3/ELK4/CDC27/PRKACB/CHEK1/NFYB/CCND2/HLA-DQB1/E2F3/NFATC2/CREB3L1/MAP2K2/JUN/PIK3CD/TNF/HLA-G/PPP3R1/MAPK9/HLA-DOA/TGFB3/MAD1L1/ANAPC16/TGFB2/MAP3K3/ZFP36/MAP3K14/ATF6B/GPS2/STAT5A/ETS1/LTBR/CREB1/PIK3R3/NFATC4/SLC25A5/PIK3CB/RB1/NFATC1/CDC16/SMAD4/POLB/ANAPC2/SLC25A4/CALR/BUB1B/HLA-DRB5/VDAC1/E2F1/ADCY6/TNFRSF1A/ATF4/CREB3/SLC2A1/IKBKB/PIK3R1/CCND1/ATM/CREB3L3/FOSL1/CDK4/TLN2/MAP3K1/HLA-DQA1/NFKB1/CHUK/JAK1/KAT2B/PPP3CA/IL2RG/XIAP/KAT5/ATR/ADCY4/ANAPC10/MAP2K4/ANAPC13/CCNE2/KRAS/CSF2/MAD2L1/CREB5/VDAC3/E2F2/CRTC2/DLG1/CDKN2C/CRTC1/TBP/CHEK2/FOS/MAPK8/HRAS/CRTC3/ESPL1/ADCY9/ANAPC5/BUB3/ADCY1/HLA-DQB2/NRP1/PPP3CB/CDKN2B/HLA-DQA2/PRKACA/TRRAP/CDC26/EP300/TBPL1/CCNE1 |
| Human Diseases | Infectious disease: viral | hsa05169 | Epstein-Barr virus infection | 131/ 3489 | 204/ 8538 | 0.64216 | 1.57143 | 6.86681 | 8.64E-12 | 1.28E-09 | 8.17E-10 | TAP1/HLA-E/TAP2/B2M/STAT1/HLA-A/HLA-C/HLA-DMA/HLA-B/BAK1/OAS2/HLA-F/ICAM1/OAS3/IRF9/HLA-DRA/ISG15/STAT2/HLA-DPB1/LYN/PSMD11/CASP8/HLA-DPA1/CIR1/HLA-DOB/HLA-DMB/EIF2AK2/PSMC4/HLA-DRB1/STAT3/IKBKG/TAPBP/TRAF6/CASP3/PSMD3/FAS/PSMC1/PDIA3/NFKBIE/CDKN1A/CDK2/MAP2K3/TBK1/CYCS/TRAF5/MYD88/MDM2/RELA/MYC/CD40/PSMD13/CCND3/PSMC6/PSMD7/IRAK1/RIPK1/SNW1/PSMD12/PSMD14/SIN3A/CCND2/HLA-DQB1/E2F3/ADRM1/PSMD1/ENTPD1/IFNAR1/RBPJ/CXCL10/JUN/PIK3CD/TNF/PSMC2/HLA-G/MAPK9/HLA-DOA/CD58/IFNAR2/CASP9/PSMD2/OAS1/MAP3K14/BID/PSMD6/PSMC3/PIK3R3/TNFAIP3/CD44/PIK3CB/RB1/CALR/HLA-DRB5/MAPK11/E2F1/HDAC2/IKBKB/IRF3/PIK3R1/CCND1/FADD/CDK4/PSMD4/HLA-DQA1/DDB2/NFKB1/IRF7/CHUK/GADD45A/JAK1/TRAF2/CDKN1B/MAP2K4/MAP2K6/CCNE2/SAP30L/E2F2/BCL2L11/PSMC5/PSMD8/MAPK8/SKP2/RUNX3/TAB2/HES1/HLA-DQB2/MAVS/MAP3K7/HLA-DQA2/TAB1/MAP2K7/CCNE1 |
| Genetic Information Processing | Folding, sorting and degradation | hsa03050 | Proteasome | 41/ 3489 | 46/ 8538 | 0.89130 | 2.18113 | 6.67685 | 1.11E-11 | 1.28E-09 | 8.17E-10 | PSMB8/PSMB9/PSME1/PSME2/PSMB10/PSMA4/PSMA1/PSMA6/PSMD11/PSMB7/PSMA3/POMP/PSMA2/PSMC4/PSMA5/PSMB2/PSMA7/PSMD3/PSMC1/PSMF1/PSMB4/PSMB1/PSMB3/PSMD13/PSMC6/PSMB6/PSMD7/PSMD12/PSMD14/PSMB5/ADRM1/PSMD1/PSME3/PSMC2/PSMD2/PSMD6/PSMC3/PSMD9/PSMD4/PSMC5/PSMD8 |
| Environmental Information Processing | Signal transduction | hsa04668 | TNF signaling pathway | 84/ 3489 | 119/ 8538 | 0.70588 | 1.72738 | 6.64208 | 3.88E-11 | 3.35E-09 | 2.14E-09 | IRF1/ICAM1/CASP10/CX3CL1/IL15/CASP8/CYLD/BIRC3/FRMD8/TNFRSF1B/CSF1/CASP7/MLKL/IKBKG/CASP3/FAS/CCL2/CXCL2/DAB2IP/MAP2K3/CREB3L4/TRAF5/RPS6KA4/RELA/CCL5/IL18R1/RIPK1/MAP3K8/CFLAR/RHBDF2/CREB3L1/CXCL10/JUN/PIK3CD/TNF/CCL20/DNM1L/MAPK9/CXCL3/RPS6KA5/MAP3K14/ATF6B/CEBPB/CREB1/PIK3R3/TNFAIP3/PIK3CB/PTGS2/MAP3K5/CXCL6/SOCS3/BIRC2/JAG1/MAPK11/TNFRSF1A/ATF4/CREB3/IKBKB/PIK3R1/FADD/CREB3L3/NFKB1/CHUK/TRAF2/XIAP/MAP2K4/MAP2K6/CSF2/CREB5/IL1B/RIPK3/JUNB/MMP14/FOS/MAPK8/TAB3/EDN1/TAB2/RHBDF1/ADAM17/PGAM5/MAP3K7/TAB1/MAP2K7 |
| Human Diseases | Infectious disease: bacterial | hsa05132 | Salmonella infection | 153/ 3489 | 251/ 8538 | 0.60956 | 1.49167 | 6.57223 | 6.03E-11 | 4.16E-09 | 2.65E-09 | CASP1/BAK1/GSDMD/CASP8/TNFSF10/TUBA1A/DYNLL2/BIRC3/NOD1/CASP7/MLKL/IKBKG/BRK1/TRAF6/CASP3/MAP2K3/EXOC2/RIPK2/VPS33A/CYCS/DYNLT1/MYD88/RELA/MYC/PIK3C3/ARF1/PKN1/IRAK1/RIPK1/MYL12A/TUBA1C/MYO6/DYNC2H1/MYL12B/RALA/ACTR10/ACTR3/VPS18/VPS11/ARPC5L/DYNC1LI1/TCF7L1/KPNA4/RAB5C/ACTR1B/TUBB2A/CASP5/DCTN6/MAP2K2/TLR5/RPS3/JUN/PIK3CD/TNF/ELMO2/ARL8A/PAK1/DYNC1H1/DYNLRB1/RILP/MAPK9/TXN/KPNA1/S100A10/KLC1/TLR4/ABI1/PFN2/KPNA3/RHOG/PFN1/PIK3CB/MYL9/VPS41/BIRC2/ARPC3/SNX9/FLNA/WASL/CYFIP2/CASP4/DYNC1LI2/MAPK11/DYNC2LI1/TNFRSF1A/DCTN4/KLC2/IL18/IKBKB/TCF7L2/TUBB6/FADD/CYTH4/NAIP/TUBB4B/AHNAK2/ACBD3/NCKAP1/M6PR/NFKB1/FHOD1/TUBB/CHUK/TRAF2/RAB9A/RAB5A/DYNLT3/VPS39/ARHGEF26/ACTB/EXOC7/RAB7A/STX10/MAP2K4/MAP2K6/TNFRSF10B/ARF6/SNX33/PODXL/FBXO22/ACTR2/TUBB3/SKP1/IL1B/KLC4/TXN2/RIPK3/AHNAK/CYTH3/HSP90AA1/FOS/ARL8B/MAPK8/TAB3/HRAS/TIRAP/NLRC4/DCTN3/PLEKHM2/FYCO1/TAB2/MYL5/KIF5B/TUBAL3/MAP3K7/ACTR1A/EXOC4/MYL2/ACTR3C/DYNC1I2/TAB1/WASF3/MAP2K7 |
| Human Diseases | Neurodegenerative disease | hsa05010 | Alzheimer disease | 214/ 3489 | 391/ 8538 | 0.54731 | 1.33934 | 5.70992 | 1.06E-08 | 6.07E-07 | 3.87E-07 | PSMA4/PSMA1/PSMA6/ATP2A3/ATG13/NDUFA9/PSMD11/PSMB7/CALM3/CASP8/PSMA3/TUBA1A/PSMA2/APH1B/EIF2AK2/CSF1/CASP7/IRS1/PSMC4/NOS2/NDUFS2/PSMA5/PSMB2/IKBKG/PSMA7/CSNK1E/CASP3/PSMD3/FAS/PSMC1/PSMB4/PPP3CC/APP/DVL2/PSMB1/UQCRFS1/CAPN1/SLC39A1/PSENEN/PSMB3/CYCS/NDUFAB1/ULK2/RELA/VDAC2/PIK3C3/PSMD13/CACNA1D/SDHA/PSMC6/PSMB6/LRP1/CYC1/APBB1/PSMD7/DKK4/TUBA1C/PSMD12/FZD7/WNT7A/NDUFV2/PSMD14/PSMB5/BACE1/WNT4/ADRM1/PSMD1/SDHB/SLC39A11/PSEN2/CALML4/NDUFS3/BACE2/TUBB2A/INSR/FZD3/NDUFS6/NDUFV1/APH1A/MAP2K2/BAD/PIK3CD/NDUFB5/TNF/SLC39A14/PLCB4/ATG14/NDUFS7/UQCRC1/PSMC2/SLC39A13/PPP3R1/MAPK9/COX5A/NDUFB3/ULK1/CASP9/PSMD2/KLC1/FZD6/BID/DVL1/NDUFB7/CSNK2A1/SLC11A2/LRP6/PSMD6/UQCRC2/SLC39A10/CSNK1A1/WNT7B/PSMC3/PIK3R3/EIF2AK3/SLC25A5/PIK3CB/PTGS2/DDIT3/MAP3K5/EIF2S1/NDUFS1/COX8A/SLC25A4/NDUFB10/ADAM10/NDUFA10/WNT3/VDAC1/COX7B/TNFRSF1A/ATF4/KLC2/AXIN2/IKBKB/NCSTN/PIK3R1/PLCB3/TUBB6/FADD/PSMD9/NDUFB9/PSEN1/ITPR2/NDUFC2/TUBB4B/HSD17B10/NRBF2/PSMD4/XBP1/NFKB1/TUBB/CHUK/WNT9A/COX7A2L/NDUFA1/TRAF2/PPP3CA/CYBB/SNCA/NDUFB8/SDHC/MTOR/AMBRA1/IL1A/ERN1/SDHD/NDUFS8/KRAS/CALM1/ATG2A/GRIN2D/TUBB3/VDAC3/NDUFA8/GRIN1/APC2/NDUFB4/IL1B/CSNK2B/WNT10A/KLC4/IRS2/WNT11/PSMC5/PSMD8/ARAF/FZD4/MAPK8/ITPR3/HRAS/ATF6/MME/ITPR1/UQCRH/PPIF/COX4I1/FZD5/APOE/ATG2B/ADAM17/KIF5B/CAPN2/PPP3CB/TUBAL3/IDE/NDUFA5/NDUFV3/SLC39A5/ATP2A2/FZD2/COX7C/MAP2K7/SLC39A3/UQCR10 |
| Cellular Processes | Cell growth and death | hsa04218 | Cellular senescence | 98/ 3489 | 157/ 8538 | 0.62420 | 1.52750 | 5.54531 | 3.21E-08 | 1.48E-06 | 9.42E-07 | HLA-E/HLA-A/HLA-C/HLA-B/HLA-F/CALM3/TGFBR1/LIN52/SMAD3/NBN/E2F5/PPP3CC/CDKN1A/CDK2/MAP2K3/CDKN2A/CAPN1/TGFB1/MDM2/RELA/MYC/VDAC2/CACNA1D/CCND3/RBBP4/RAD9A/CHEK1/TRPV4/CCND2/E2F3/NFATC2/CALML4/SQSTM1/RBL1/MAP2K2/RBL2/PIK3CD/FOXO3/HLA-G/PPP3R1/TGFB3/ZFP36L1/TGFB2/TRAF3IP2/CDC25A/ETS1/TRPM7/PIK3R3/NFATC4/SLC25A5/PIK3CB/RB1/NFATC1/SLC25A4/MAPKAPK2/TSC2/MAPK11/E2F4/MRAS/VDAC1/E2F1/TSC1/PIK3R1/RAD1/CCND1/ATM/MYBL2/ITPR2/CDK4/NFKB1/FOXO1/LIN37/LIN9/GADD45A/PPP3CA/MTOR/ATR/IL1A/MAP2K6/CCNE2/KRAS/CALM1/VDAC3/LIN54/E2F2/FBXW11/SIRT1/CHEK2/RAD50/ITPR3/HRAS/PPP1CC/ITPR1/CAPN2/IGFBP3/PPP3CB/CDKN2B/CCNE1 |
| Human Diseases | Neurodegenerative disease | hsa05022 | Pathways of neurodegeneration - multiple diseases | 255/ 3489 | 483/ 8538 | 0.52795 | 1.29196 | 5.49112 | 3.42E-08 | 1.48E-06 | 9.42E-07 | UBE2L6/BAK1/PSMA4/PSMA1/PSMA6/ATP2A3/ATG13/NDUFA9/UBB/PSMD11/PSMB7/CALM3/CASP8/OPTN/UBC/PSMA3/TUBA1A/PSMA2/TNFRSF1B/CSF1/CASP7/PSMC4/NOS2/NDUFS2/MFN2/PSMA5/PSMB2/SOD1/PSMA7/PRKCG/CSNK1E/UBA7/CASP3/PSMD3/FAS/PSMC1/PSMB4/PPP3CC/APP/HIP1/SPG11/DVL2/PSMB1/UQCRFS1/MAP2K3/TBK1/CAPN1/PLCG1/PSMB3/CYCS/NDUFAB1/FIG4/ULK2/RELA/VDAC2/PIK3C3/PSMD13/CACNA1D/SDHA/PSMC6/PSMB6/CYC1/PSMD7/DKK4/TUBA1C/TOMM40L/DNAL4/GPX8/PSMD12/CAT/FZD7/HTT/WNT7A/NDUFV2/PSMD14/ACTR10/PSMB5/WNT4/DERL1/TRAP1/IFT57/ADRM1/GPX2/PSMD1/SDHB/DNALI1/PSEN2/ACTR1B/ATXN2L/CALML4/DNAH12/NDUFS3/SQSTM1/TUBB2A/FZD3/DCTN6/NDUFS6/NDUFV1/MAP2K2/BAD/GPX3/DNAL1/NDUFB5/TNF/PLCB4/MAP3K10/ATG14/NDUFS7/UQCRC1/PSMC2/PPP3R1/MAPK9/COX5A/NDUFB3/ULK1/UBE2J2/STX1A/CASP9/PSMD2/KLC1/UBE2L3/FZD6/BID/DVL1/NDUFB7/SMCR8/CSNK2A1/LRP6/PSMD6/UQCRC2/SPTBN2/DNAH6/CSNK1A1/WNT7B/UBE2G1/PSMC3/EIF2AK3/SLC25A5/PTGS2/DDIT3/DNAH5/MAP3K5/EIF2S1/ALS2/NDUFS1/COX8A/RAB8A/SLC25A4/NDUFB10/NDUFA10/TANK/WNT3/MAPK11/VDAC1/DNAH2/COX7B/TNFRSF1A/ATF4/DCTN4/KLC2/AXIN2/DNAH11/PLCB3/VCP/TUBB6/FADD/PSMD9/NDUFB9/MAP1LC3B/PSEN1/ATXN3/ITPR2/NDUFC2/TUBB4B/HSD17B10/NRBF2/PSMD4/XBP1/NFKB1/TUBB/WNT9A/COX7A2L/NDUFA1/RAB1A/TRAF2/PPP3CA/CYBB/CAMK2G/SNCA/RAB5A/NDUFB8/SDHC/MTOR/AMBRA1/IL1A/MAP2K6/ERN1/SDHD/NDUFS8/C9orf72/KRAS/CALM1/ATG2A/PRNP/GRIN2D/CAMK2D/MFN1/TUBB3/VDAC3/NDUFA8/GRIN1/APC2/NDUFB4/IL1B/GPX1/CSNK2B/PRKCA/WNT10A/KLC4/WNT11/PSMC5/PSMD8/ARAF/FZD4/MAPK8/ITPR3/HRAS/UBA52/UBA1/ATF6/CCS/TOMM40/DCTN3/HSPA5/WDR41/ITPR1/UQCRH/PPIF/COX4I1/FZD5/ATG2B/KIF5B/CAPN2/PPP3CB/TUBAL3/ACTR1A/DNAH9/RPS27A/NDUFA5/NDUFV3/ATP2A2/FZD2/COX7C/MAP2K7/UQCR10 |
| Cellular Processes | Transport and catabolism | hsa04144 | Endocytosis | 144/ 3489 | 252/ 8538 | 0.57143 | 1.39835 | 5.33577 | 9.12E-08 | 3.50E-06 | 2.23E-06 | HLA-E/HLA-A/HLA-C/HLA-B/HLA-F/GIT2/RAB31/PML/ARF3/IGF1R/TGFBR1/IQSEC1/SNX6/SMAD3/ARRB1/AP2A1/EPN3/DNM3/TRAF6/FGFR2/EHD2/FGFR3/VPS37A/PIP5K1B/ARAP3/DNM1/HSPA1A/MDM2/RAB10/TSG101/FGFR4/TFRC/ARF1/VPS4A/FOLR1/PSD3/ARFGAP1/SNX2/GIT1/AGAP2/ACTR3/SH3GLB1/SH3KBP1/CHMP5/ARPC5L/SMAP2/SNX3/PARD6G/ARFGAP3/EHD1/RAB5C/RAB11B/SNX32/CXCR4/AP2A2/AGAP1/RAB11FIP3/ARF5/VPS37B/PARD6A/HLA-G/HGS/PLD2/ARAP2/RAB35/HSPA1B/SNX1/CAV1/CHMP1A/HSPA1L/CLTB/VPS4B/STAM/EPS15/CLTA/GRK5/GRK6/LDLRAP1/CLTC/PARD6B/CBLB/CAPZA1/ARFGAP2/ARPC3/WASL/RAB8A/CHMP4B/IST1/STAM2/SH3GLB2/CYTH4/RAB11FIP2/ASAP2/ACAP2/EPS15L1/SNX12/RAB5A/IL2RG/ZFYVE9/RAB7A/CHMP4A/CAV2/BIN1/CHMP3/ARF6/ARRB2/CHMP4C/SNX5/VTA1/WIPF2/ASAP1/ACTR2/CBL/PIP5K1C/PRKCZ/VPS26A/VPS35/USP8/SRC/ARFGEF1/CYTH3/EHD4/VPS45/VPS28/HRAS/RNF41/ACAP1/PDCD6IP/CLTCL1/RAB11FIP5/PARD3/CHMP6/HSPA8/DAB2/GBF1/CHMP7/EPN1/KIF5B/AGAP3/ACTR3C/ASAP3/WIPF3/CHMP2A/ZFYVE27 |
| Genetic Information Processing | Folding, sorting and degradation | hsa04120 | Ubiquitin mediated proteolysis | 89/ 3489 | 142/ 8538 | 0.62676 | 1.53376 | 5.33154 | 1.04E-07 | 3.60E-06 | 2.29E-06 | UBE2L6/UBB/PML/RHOBTB2/UBC/BIRC3/CUL7/UBA6/CUL2/UBOX5/UBA7/TRAF6/UBE2Z/PIAS3/UBE2D3/MID1/FBXO2/NHLRC1/MDM2/CDC27/SOCS1/CUL3/WWP2/VHL/UBE2E3/SYVN1/UBE2M/UBE3B/HERC1/FBXO4/UBE2Q1/TRIP12/UBE2J2/ANAPC16/UBE2L3/HERC4/UBE2Q2/UBE2F/RHOBTB1/KEAP1/UBE2G1/CBLB/UBE4B/STUB1/CDC16/UBE2E1/RNF7/UBE2O/SOCS3/ANAPC2/BIRC2/RBX1/SAE1/UBE2D2/UBE4A/FBXW7/UBE2N/FANCL/KLHL13/TRIM37/MAP3K1/DDB2/UBE2E2/PIAS4/DET1/XIAP/ANAPC10/ANAPC13/CDC34/TRIM32/RCHY1/CBL/SKP1/BIRC6/UBE2K/FBXW11/HERC3/UBA3/UBA2/UBA52/UBA1/SKP2/ANAPC5/DDB1/MGRN1/RPS27A/PIAS2/CDC26/SIAH1 |
| Genetic Information Processing | Folding, sorting and degradation | hsa04141 | Protein processing in endoplasmic reticulum | 103/ 3489 | 170/ 8538 | 0.60588 | 1.48267 | 5.28398 | 1.30E-07 | 4.08E-06 | 2.60E-06 | BAK1/FBXO6/DNAJA1/DNAJA2/BAG1/SIL1/NPLOC4/DNAJC1/SEC24C/UBE2D3/PDIA3/CAPN1/FBXO2/HSPA1A/YOD1/UBXN4/PREB/SVIP/SEC31A/RAD23B/LMAN2/DERL1/UBQLN1/SYVN1/PLAA/SSR4/SEC24A/PRKCSH/P4HB/SSR2/DNAJB11/MAPK9/NGLY1/UBE2J2/HSPA1B/SEC61B/NSFL1C/EDEM2/ATF6B/GANAB/HSPA1L/SEC23B/UBQLN2/UBXN6/UBE2G1/MBTPS1/SSR1/UBE4B/EIF2AK3/DNAJC5/STUB1/DDIT3/RRBP1/MAP3K5/DERL2/DNAJB2/EIF2S1/SSR3/RBX1/CALR/UBE2D2/ATF4/BAG2/SEC24B/MAN1B1/VCP/HYOU1/ATXN3/UGGT2/XBP1/SEC23A/DNAJB12/HSPBP1/TRAF2/SEL1L/DNAJC3/MAGT1/CKAP4/ERN1/SEC63/SAR1A/ERP29/SEC13/SAR1B/SKP1/TRAM1/DAD1/DERL3/UBXN2A/HSP90AA1/TXNDC5/MAPK8/PPP1R15A/ATF6/HSPA5/UBQLN4/HSPA8/UGGT1/CAPN2/BCAP31/OSTC/MAP2K7/MBTPS2 |
| Human Diseases | Infectious disease: parasitic | hsa05145 | Toxoplasmosis | 73/ 3489 | 112/ 8538 | 0.65179 | 1.59500 | 5.26883 | 1.51E-07 | 4.34E-06 | 2.77E-06 | STAT1/HLA-DMA/HLA-DRA/JAK2/HLA-DPB1/CASP8/HLA-DPA1/BIRC3/HLA-DOB/HLA-DMB/NOS2/HLA-DRB1/STAT3/IKBKG/TRAF6/CASP3/LAMC2/MAP2K3/CIITA/LAMB2/HSPA1A/TGFB1/CYCS/MYD88/RELA/CD40/SOCS1/IRAK1/ALOX5/HLA-DQB1/GNAI2/BAD/LAMC1/TNF/MAPK9/HLA-DOA/TGFB3/HSPA1B/CASP9/TGFB2/TLR4/IFNGR2/IFNGR1/HSPA1L/LAMA5/ITGA6/BIRC2/HLA-DRB5/MAPK11/TNFRSF1A/LAMB3/IKBKB/GNAI3/HLA-DQA1/ITGB1/NFKB1/CHUK/JAK1/LAMA1/XIAP/MAP2K6/IL12A/MAPK8/LAMA4/TAB2/HSPA8/PPIF/HLA-DQB2/MAP3K7/LAMA3/HLA-DQA2/TAB1/PDPK1 |
| Human Diseases | Infectious disease: viral | hsa05161 | Hepatitis B | 98/ 3489 | 163/ 8538 | 0.60123 | 1.47127 | 5.04982 | 4.39E-07 | 1.05E-05 | 6.68E-06 | STAT1/CASP10/JAK2/STAT2/IFIH1/CASP8/TGFBR1/STAT5B/SMAD3/STAT3/IKBKG/TICAM1/PRKCG/TRAF6/CASP3/FAS/TLR3/CDKN1A/CDK2/MAP2K3/TBK1/CREB3L4/TGFB1/CYCS/MYD88/RELA/MYC/IRAK1/ATP6AP1/HSPG2/STAT6/E2F3/NFATC2/IFNAR1/YWHAB/CREB3L1/MAP2K2/SOS1/BAD/JUN/PIK3CD/TNF/YWHAQ/MAPK9/TGFB3/CASP9/TGFB2/TLR4/ATF6B/BID/PTK2B/STAT5A/CREB1/PIK3R3/NFATC4/PIK3CB/RB1/NFATC1/SMAD4/MAPK11/E2F1/ATF4/CREB3/IKBKB/IRF3/PIK3R1/FADD/CREB3L3/MAP3K1/DDB2/NFKB1/IRF7/CHUK/JAK1/YWHAZ/MAP2K4/MAP2K6/CCNE2/KRAS/CREB5/VDAC3/E2F2/SRC/PRKCA/ARAF/FOS/MAPK8/HRAS/TIRAP/TAB2/DDB1/MAVS/MAP3K7/BIRC5/TAB1/EP300/MAP2K7/CCNE1 |
| Cellular Processes | Transport and catabolism | hsa04140 | Autophagy - animal | 101/ 3489 | 169/ 8538 | 0.59763 | 1.46248 | 5.04777 | 4.41E-07 | 1.05E-05 | 6.68E-06 | ATG13/UBB/OPTN/IGF1R/WDR45/UBC/IRS1/DAPK1/CALCOCO2/TRAF6/YKT6/RRAGC/TBK1/RRAGB/GABARAP/ATG3/VPS33A/ULK2/PIK3C3/PPP2CA/MTMR14/PRKACB/STX17/IGBP1/CFLAR/VPS18/SH3GLB1/HMGB1/VPS11/TAX1BP1/SQSTM1/RRAGD/VAMP8/MAP2K2/ATG16L1/BAD/PIK3CD/DAPK2/PPP2CB/ATG14/ATG4A/MAPK9/ULK1/AKT1S1/PRKAA1/WDFY3/SMCR8/PRKAA2/CAMKK2/PIK3R3/EIF2AK3/PIK3CB/ATG12/VPS41/EIF2S1/RAB8A/UVRAG/LAMP2/TSC2/TANK/MRAS/HIF1A/TSC1/PIK3R1/DAPK3/VMP1/MAP1LC3B/RAB33B/NRBF2/RPS6KB2/GORASP1/RAB1A/MTOR/VPS39/EIF2AK4/RAB7A/AMBRA1/ERN1/C9orf72/KRAS/ATG2A/BIRC6/CTSD/ATG4B/ATG7/IRS2/MAPK8/HRAS/ATG16L2/UBA52/RRAGA/WDR41/ITPR1/PRKCQ/ATG2B/MAP3K7/ATG10/RPS27A/LAMP1/PRKACA/PDPK1 |
| Human Diseases | Neurodegenerative disease | hsa05014 | Amyotrophic lateral sclerosis | 198/ 3489 | 371/ 8538 | 0.53369 | 1.30601 | 5.00947 | 4.55E-07 | 1.05E-05 | 6.68E-06 | CASP1/PSMA4/PSMA1/PSMA6/ATG13/NDUFA9/PSMD11/PSMB7/OPTN/PSMA3/TUBA1A/PSMA2/TNFRSF1B/PSMC4/NOS2/NDUFS2/ANG/PSMA5/PSMB2/SOD1/PSMA7/CASP3/PSMD3/PSMC1/PSMB4/PPP3CC/SPG11/SETX/PSMB1/UQCRFS1/MAP2K3/TBK1/PSMB3/CYCS/NDUFAB1/FIG4/ULK2/PIK3C3/PSMD13/SDHA/PSMC6/PSMB6/CYC1/PSMD7/TUBA1C/TOMM40L/DNAL4/GPX8/PSMD12/CAT/NDUFV2/PSMD14/ACTR10/PSMB5/DERL1/UBQLN1/ADRM1/GPX2/PSMD1/SDHB/DNALI1/ACTR1B/ATXN2L/NUP43/DNAH12/NDUFS3/SQSTM1/TUBB2A/DCTN6/NDUFS6/NDUFV1/BAD/NUP210/GPX3/DNAL1/NDUFB5/TNF/ATG14/NDUFS7/UQCRC1/PSMC2/PPP3R1/COX5A/NDUFB3/ULK1/NRG4/CASP9/PSMD2/KLC1/PFN2/BID/NDUFB7/NUP50/SMCR8/UBQLN2/PSMD6/HNRNPA1/UQCRC2/DNAH6/PFN1/PSMC3/ANXA7/EIF2AK3/DDIT3/NRG1/DNAH5/MAP3K5/NUP98/EIF2S1/ALS2/NDUFS1/COX8A/RAB8A/RAE1/NDUFB10/NDUFA10/TANK/MAPK11/VDAC1/DNAH2/COX7B/TNFRSF1A/ATF4/DCTN4/KLC2/DNAH11/CHCHD10/VCP/TUBB6/PSMD9/NDUFB9/MAP1LC3B/NDUFC2/TUBB4B/NRBF2/PSMD4/XBP1/TUBB/NUP153/COX7A2L/NDUFA1/RAB1A/TRAF2/PPP3CA/RAB5A/NDUFB8/SDHC/MTOR/ANXA11/ACTB/AMBRA1/MAP2K6/ERN1/SDHD/NDUFS8/C9orf72/SEC13/ATG2A/GRIN2D/TUBB3/NDUFA8/GRIN1/NDUFB4/SEH1L/GPX1/KLC4/PSMC5/PSMD8/NUP205/ITPR3/ATF6/CCS/TOMM40/HNRNPA2B1/DCTN3/NUP188/HSPA5/SRSF3/NUP85/NUP35/UBQLN4/WDR41/UQCRH/COX4I1/ATG2B/KIF5B/ALYREF/PPP3CB/NUP214/TUBAL3/ACTR1A/DNAH9/NDUFA5/NDUFV3/NXF1/POM121C/COX7C/UQCR10 |
| Human Diseases | Infectious disease: viral | hsa05165 | Human papillomavirus infection | 179/ 3489 | 333/ 8538 | 0.53754 | 1.31542 | 4.88057 | 8.82E-07 | 1.90E-05 | 1.21E-05 | HLA-E/IRF1/STAT1/HLA-A/HLA-C/HLA-B/BAK1/HLA-F/IRF9/ISG15/STAT2/MX1/CASP8/ATP6V1B2/EIF2AK2/ITGB5/TCIRG1/IKBKG/TICAM1/CASP3/FAS/PSMC1/LAMC2/DVL2/TLR3/TUBG2/CDKN1A/CDK2/TBK1/LAMB2/CREB3L4/COL6A1/MX2/MDM2/RELA/PPP2CA/OASL/ITGAV/LFNG/CCND3/ATP6V0B/ATP6V0D1/ATP6AP1/PRKACB/PPP2R3A/FZD7/WNT7A/ATP6V1H/WNT4/CCND2/VTN/ITGA1/TCF7L1/PARD6G/PTK2/MAGI1/RFNG/IFNAR1/RBPJ/RBL1/FZD3/CREB3L1/COL9A2/MAP2K2/SOS1/RBL2/BAD/LAMC1/NOTCH3/PIK3CD/PPP2CB/TNF/ATP6V0C/ATP6V0E2/PARD6A/HLA-G/IFNAR2/FZD6/DVL1/THBS1/ATP6V1E1/LAMA5/CSNK1A1/CREB1/WNT7B/PARD6B/PIK3R3/PIK3CB/RB1/PTGS2/ITGA6/ATP6V1A/PPP2R5C/JAG1/ITGA3/NOTCH4/ITGA9/TSC2/WNT3/ATP6V0A2/COL4A2/E2F1/MAML3/TNFRSF1A/HDAC2/ITGA5/ITGB8/LAMB3/ATP6V1D/CREB3/AXIN2/IKBKB/IRF3/TSC1/PIK3R1/HES2/CCND1/TCF7L2/ATM/FADD/ITGB6/PSEN1/CREB3L3/CDK4/ATP6V1G1/RPS6KB2/ITGB1/NFKB1/FOXO1/CHUK/NOTCH2/WNT9A/JAK1/ITGA7/LAMA1/PXN/MTOR/ATR/CDKN1B/CCNE2/ITGB7/KRAS/GNAS/CREB5/ITGB4/APC2/PRKCZ/CHAD/DLG1/WNT10A/HES6/THBS3/WNT11/TBP/CRB3/FZD4/UBR4/HRAS/ATP6V1F/TADA3/CHD4/MAML1/LAMA4/COMP/PPP2R2D/MAML2/HES1/PARD3/LLGL2/FZD5/BCAP31/ITGA2B/LAMA3/PRKACA/FZD2/NOTCH1/EP300/TBPL1/CCNE1 |
| Human Diseases | Neurodegenerative disease | hsa05016 | Huntington disease | 168/ 3489 | 311/ 8538 | 0.54019 | 1.32192 | 4.80732 | 1.28E-06 | 2.42E-05 | 1.55E-05 | PSMA4/PSMA1/PSMA6/TGM2/ATG13/NDUFA9/PSMD11/PSMB7/CASP8/PSMA3/TUBA1A/PSMA2/PSMC4/NDUFS2/PPARG/AP2A1/PSMA5/SOD2/PSMB2/SOD1/PSMA7/CASP3/PSMD3/PSMC1/PSMB4/HIP1/PSMB1/UQCRFS1/CREB3L4/PSMB3/CYCS/NDUFAB1/ULK2/VDAC2/PIK3C3/PSMD13/SDHA/SLC1A3/PSMC6/PSMB6/CYC1/PSMD7/POLR2J/TUBA1C/DNAL4/GPX8/PSMD12/HTT/NDUFV2/PSMD14/SIN3A/ACTR10/PSMB5/IFT57/ADRM1/GPX2/PSMD1/SDHB/DNALI1/ACTR1B/DNAH12/NDUFS3/TUBB2A/AP2A2/DCTN6/NDUFS6/NDUFV1/CREB3L1/GPX3/DNAL1/NDUFB5/PLCB4/MAP3K10/ATG14/NDUFS7/UQCRC1/PSMC2/MAPK9/COX5A/NDUFB3/ULK1/STX1A/CASP9/PSMD2/KLC1/SP1/CLTB/NDUFB7/PSMD6/UQCRC2/CLTA/DNAH6/CREB1/CLTC/PSMC3/SLC25A5/DNAH5/MAP3K5/POLR2I/NDUFS1/COX8A/SLC25A4/NDUFB10/NDUFA10/VDAC1/DNAH2/COX7B/HDAC2/DCTN4/KLC2/CREB3/DNAH11/PLCB3/TUBB6/PSMD9/NDUFB9/CREB3L3/NDUFC2/TUBB4B/NRBF2/PSMD4/TUBB/COX7A2L/NDUFA1/TRAF2/NDUFB8/SDHC/MTOR/AMBRA1/ERN1/SDHD/NDUFS8/BBC3/ATG2A/CREB5/TUBB3/VDAC3/NDUFA8/GRIN1/NDUFB4/GPX1/KLC4/PSMC5/TBP/PSMD8/MAPK8/POLR2B/POLR2A/DCTN3/CLTCL1/ITPR1/UQCRH/PPIF/COX4I1/ATG2B/KIF5B/TUBAL3/ACTR1A/DNAH9/NDUFA5/NDUFV3/POLR2E/POLR2J3/EP300/COX7C/TBPL1/MAP2K7/UQCR10 |
| Human Diseases | Neurodegenerative disease | hsa05012 | Parkinson disease | 149/ 3489 | 271/ 8538 | 0.54982 | 1.34546 | 4.80412 | 1.33E-06 | 2.42E-05 | 1.55E-05 | UBE2L6/PSMA4/PSMA1/PSMA6/NDUFA9/UBB/PSMD11/PSMB7/CALM3/UBC/PSMA3/TUBA1A/PSMA2/PSMC4/NDUFS2/MFN2/PSMA5/PSMB2/SOD1/PSMA7/UBA7/CASP3/PSMD3/PSMC1/PSMB4/PSMB1/UQCRFS1/SLC39A1/PLCG1/PSMB3/CYCS/NDUFAB1/VDAC2/PSMD13/SDHA/PSMC6/PSMB6/CYC1/PSMD7/TUBA1C/DUSP1/PSMD12/PRKACB/NDUFV2/PSMD14/PSMB5/TRAP1/ADRM1/PSMD1/SDHB/GNAI2/SLC39A11/CALML4/NDUFS3/TUBB2A/NDUFS6/NDUFV1/NDUFB5/ADORA2A/SLC39A14/DRD1/NDUFS7/UQCRC1/PSMC2/SLC39A13/MAPK9/COX5A/TXN/NDUFB3/UBE2J2/CASP9/PSMD2/KLC1/UBE2L3/NDUFB7/SLC11A2/PSMD6/UQCRC2/SLC39A10/KEAP1/UBE2G1/PSMC3/EIF2AK3/SLC25A5/DDIT3/MAP3K5/EIF2S1/NDUFS1/COX8A/SLC25A4/NDUFB10/NDUFA10/VDAC1/COX7B/ATF4/KLC2/TUBB6/PSMD9/GNAI3/NDUFB9/ITPR2/NDUFC2/TUBB4B/PSMD4/XBP1/TUBB/COX7A2L/NDUFA1/CAMK2G/SNCA/NDUFB8/SDHC/ERN1/MAOB/SDHD/NDUFS8/CALM1/GNAS/CAMK2D/MFN1/TUBB3/VDAC3/NDUFA8/NDUFB4/KLC4/TXN2/PSMC5/PSMD8/MAPK8/ITPR3/UBA52/UBA1/ATF6/MAOA/HSPA5/ITPR1/UQCRH/PPIF/COX4I1/KIF5B/TUBAL3/RPS27A/NDUFA5/NDUFV3/PRKACA/SLC39A5/COX7C/SLC39A3/UQCR10 |
| Human Diseases | Infectious disease: viral | hsa05168 | Herpes simplex virus 1 infection | 106/ 3489 | 182/ 8538 | 0.58242 | 1.42525 | 4.82033 | 1.33E-06 | 2.42E-05 | 1.55E-05 | TAP1/HLA-E/TAP2/B2M/STAT1/HLA-A/HLA-C/HLA-DMA/HLA-B/BAK1/CD74/OAS2/HLA-F/SP100/OAS3/IRF9/HLA-DRA/JAK2/STAT2/IFIH1/HLA-DPB1/PML/ZC3HAV1/CASP8/HLA-DPA1/BIRC3/HLA-DOB/HLA-DMB/EIF2AK2/HLA-DRB1/BST2/IKBKG/TAPBP/TICAM1/TRAF6/CASP3/FAS/CCL2/PDIA3/TLR3/TBK1/CYCS/TRAF5/MYD88/RELA/CCL5/IRAK1/HLA-DQB1/IFNAR1/BAD/PIK3CD/TNF/HLA-G/HLA-DOA/IFNAR2/CASP9/C3/OAS1/IFNGR2/IFNGR1/BID/RNASEL/PIK3R3/EIF2AK3/PIK3CB/SOCS3/EIF2S1/BIRC2/CALR/TSC2/HLA-DRB5/TNFRSF1A/ITGA5/IKBKB/IRF3/TSC1/PIK3R1/SRSF9/FADD/HLA-DQA1/NFKB1/IRF7/CHUK/SRSF8/JAK1/TRAF2/TNFRSF14/MTOR/EIF2B2/POU2F1/EIF2AK4/IL12A/IL1B/EIF2AK1/SRC/PPP1CC/TAB2/SRSF3/HLA-DQB2/MAVS/ALYREF/MAP3K7/HLA-DQA2/SRSF1/NXF1/TAB1 |
| Human Diseases | Neurodegenerative disease | hsa05017 | Spinocerebellar ataxia | 87/ 3489 | 144/ 8538 | 0.60417 | 1.47847 | 4.81338 | 1.45E-06 | 2.49E-05 | 1.59E-05 | PSMA4/PSMA1/PSMA6/ATP2A3/ATG13/PSMD11/PSMB7/PSMA3/PSMA2/PSMC4/PSMA5/PSMB2/PSMA7/PRKCG/PSMD3/PSMC1/PSMB4/PSMB1/GTF2B/PSMB3/CYCS/ULK2/VDAC2/PIK3C3/PSMD13/PSMC6/PSMB6/PSMD7/PSMD12/PSMD14/PSMB5/ADRM1/PSMD1/ATXN2L/RBPJ/PIK3CD/PLCB4/ATG14/PSMC2/MAPK9/KCNC3/ULK1/PSMD2/SP1/PSMD6/SPTBN2/PSMC3/PIK3R3/SLC25A5/PIK3CB/MAP3K5/SLC25A4/VDAC1/PIK3R1/PLCB3/PSMD9/ATXN3/ITPR2/NRBF2/PSMD4/XBP1/TRAF2/MTOR/NOP56/KAT5/GRIN3A/AMBRA1/ERN1/ATG2A/GRIN2D/VDAC3/GRIN1/RORA/PRKCA/OMA1/PSMC5/TBP/PSMD8/MAPK8/PUM1/ITPR3/OPA1/ITPR1/PPIF/ATG2B/ATP2A2/TBPL1 |

5. Gene Set Enrichment Analysis (GSEA) : GO BP (Biological Process)

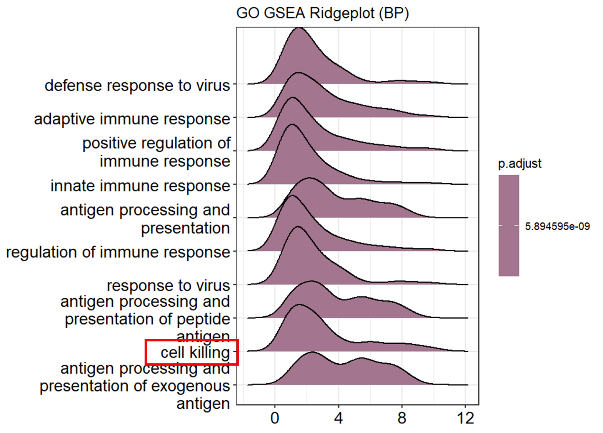

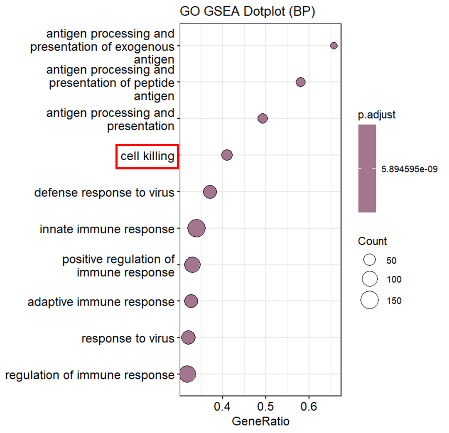

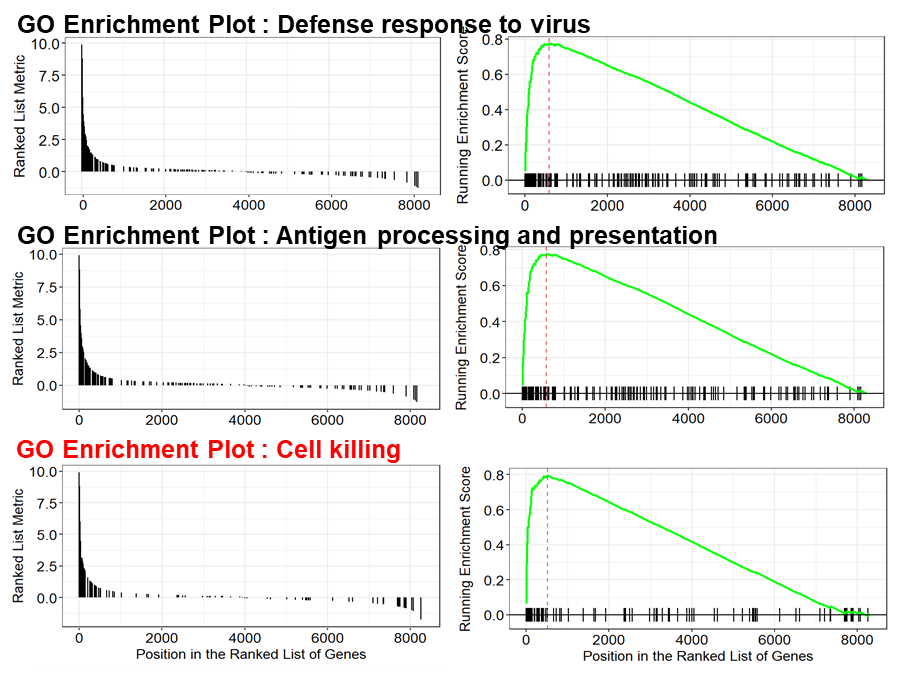

Top 20 enriched GO biological process (BP) term

| **ID** | **Description** | **set Size** | **ES** | **NES** | **p-value** | **p.adjust** | **q-value** | **leading edge** | **Core enrichment** |
| --- | --- | --- | --- | --- | --- | --- | --- | --- | --- |
| GO:0051607 | defense response to virus | 186 | 0.77563 | 2.89360 | 1.00E-10 | 5.89E-09 | 4.73E-09 | tags=37%, list=7%, signal=35% | GBP5/CXCL9/CXCL10/IFI44L/BST2/OAS2/IFITM1/GBP1/NLRC5/ZBP1/FGL2/CD40/APOBEC3G/IFI6/RSAD2/IFIT3/ISG15/GBP2/MX1/STAT1/OAS3/CASP1/IRF1/SAMHD1/IFITM3/IFIT1/PARP9/DDX60/AIM2/OASL/IFIT5/IFI27/MX2/RTP4/USP18/IFIH1/IFI16/APOBEC3D/ISG20/IFIT2/STAT2/PMAIP1/IRF9/OAS1/DDX60L/DTX3L/NLRP6/TRIM22/IL15/GBP3/MICB/MLKL/ZNFX1/TRIM31/TRIM21/DHX58/HERC5/EIF2AK2/IFITM2/APOBEC3H/TLR3/PML/MOV10/ADAR/APOBEC3F/ZC3HAV1/TRIM56/ZDHHC11/TRIM38 |
| GO:0002250 | adaptive immune response | 202 | 0.75353 | 2.83280 | 1.00E-10 | 5.89E-09 | 4.73E-09 | tags=33%, list=6%, signal=31% | SERPING1/HLA-DPA1/HLA-DPB1/HLA-DOA/HLA-DRA/HLA-DQA1/C1S/HLA-DRB1/HLA-DRB5/HLA-DQB1/CD74/HLA-DQA2/TNFSF13B/HLA-DQB2/HLA-DMB/CD274/C2/CD40/C1R/HLA-DMA/RSAD2/CFI/HLA-F/HLA-B/CLC/HLA-DOB/LAMP3/TAP1/ICAM1/IRF1/BTN3A1/HLA-E/SECTM1/HLA-C/TAP2/IL18BP/B2M/BTN3A3/HLA-A/PRDM1/BTN3A2/ASCL2/CD7/C3/JAK2/CTSS/ADA/IL12A/ERAP2/MICB/LAT2/PRKCQ/HLA-G/ALCAM/TNFRSF1B/IL18R1/C8G/CD55/IL6R/FCER1G/PARP3/RNF19B/IL20RB/SOCS3/CEACAM1/RIPK2 |
| GO:0050778 | positive regulation of immune response | 348 | 0.70287 | 2.80135 | 1.00E-10 | 5.89E-09 | 4.73E-09 | tags=33%, list=10%, signal=31% | GBP5/SERPING1/IDO1/HLA-DPA1/HLA-DPB1/HLA-DOA/HLA-DRA/HLA-DQA1/C1S/HLA-DRB1/CFH/HLA-DRB5/HLA-DQB1/CD74/HLA-DQA2/TNFSF13B/HLA-DQB2/HLA-DMB/CD274/GBP1/NLRC5/C2/ZBP1/CD40/C1R/HLA-DMA/CD38/RSAD2/CFI/HLA-F/HLA-B/HLA-DOB/GBP2/SLC15A3/OAS3/CASP1/CD5L/IRF1/BTN3A1/BTN1A1/PARP9/HLA-E/DDX60/SECTM1/HLA-C/TAP2/B2M/IFI35/BTN3A3/HLA-A/AIM2/OASL/BTN3A2/IFIH1/CD7/IFI16/CFB/C3/BIRC3/BTN2A2/EREG/OAS1/JAK2/CTSS/NLRP6/ADA/PDE4B/IL15/NMI/IL12A/PIK3AP1/MICB/ZNFX1/TRIM31/LAT2/PRKCQ/HLA-G/BANK1/DHX58/LGALS9/EIF2AK2/IL18R1/C8G/CD55/FCER1G/CCL5/TLR3/ALPK1/RBCK1/CYLD/NR1H4/CEACAM1/RIPK2/CD47/ZC3HAV1/TRIL/TRIM56/TRIM15/OTUD4/LYN/RASGRP1/WDFY1/PRKD2/TICAM1/NLRC3/PLSCR1/RNF31/S100A14/NOD1/TRIM25/UBR2/USP15/POLR3G/HSPD1/IRF7 |
| GO:0045087 | innate immune response | 441 | 0.68968 | 2.80120 | 1.00E-10 | 5.89E-09 | 4.73E-09 | tags=34%, list=10%, signal=32% | GBP5/SERPING1/HLA-DPA1/CXCL10/GBP4/C1S/CFH/CIITA/BST2/CD74/OAS2/CD274/IFITM1/GBP1/NLRC5/C2/ZBP1/NOS2/UBE2L6/CD40/C1R/APOBEC3G/IFI6/RSAD2/CFI/IFIT3/HLA-F/HLA-B/UBD/ISG15/GBP2/MX1/STAT1/SLC15A3/OAS3/CASP1/TRIM17/IRF1/SAMHD1/IFITM3/IFIT1/PARP9/HLA-E/DDX60/APOL1/GBP6/HLA-C/NCF1/B2M/IFI35/TRIM40/HLA-A/PRDM1/AIM2/OASL/IFIT5/IFI27/MX2/USP18/TRIM69/IFIH1/IFI16/APOBEC3D/ISG20/IFIT2/PARP14/STAT2/ASS1/CFB/C3/BIRC3/EREG/OAS1/DTX3L/JAK2/CTSS/NLRP6/TRIM22/IRF8/NMI/IL12A/NUB1/PIK3AP1/SAMD9/GBP3/CSF1/ZNFX1/TRIM31/TRIM21/HLA-G/DHX58/SP100/TRAFD1/LGALS9/HERC5/GSDMD/EIF2AK2/IFITM2/APOBEC3H/C8G/CD55/FCER1G/CCL5/TLR3/PML/ADAR/CSF1R/ALPK1/RNF19B/OPTN/LCN2/HK1/CITED1/CYLD/NR1H4/UBA7/CEACAM1/APOBEC3F/RIPK2/CASP8/CD47/ZC3HAV1/TRIL/KYNU/TRIM56/TRIM14/ZDHHC11/TRIM38/GCH1/TRIM26/TRIM15/OTUD4/LYN/RASGRP1/WDFY1/CALCOCO2/USP38/TICAM1/IFNE/NLRC3/PLSCR1/S100A14/NOD1/TRIM25/RAB43/USP15/ENDOD1/POLR3G/HSPD1/IRF7 |
| GO:0019882 | antigen processing and presentation | 67 | 0.86248 | 2.79827 | 1.00E-10 | 5.89E-09 | 4.73E-09 | tags=49%, list=4%, signal=47% | HLA-DPA1/HLA-DPB1/HLA-DOA/HLA-DRA/HLA-DQA1/HLA-DRB1/HLA-DRB5/HLA-DQB1/CD74/HLA-DQA2/HLA-DQB2/HLA-DMB/FGL2/HLA-DMA/HLA-F/HLA-B/IFI30/HLA-DOB/TAP1/ICAM1/HLA-E/HLA-C/TAP2/B2M/HLA-A/PSMB8/PSME1/CTSS/ERAP2/LGMN/TAPBPL/MICB/HLA-G |
| GO:0050776 | regulation of immune response | 424 | 0.69096 | 2.79299 | 1.00E-10 | 5.89E-09 | 4.73E-09 | tags=32%, list=10%, signal=30% | GBP5/SERPING1/IDO1/HLA-DPA1/HLA-DPB1/HLA-DOA/HLA-DRA/HLA-DQA1/C1S/HLA-DRB1/CFH/HLA-DRB5/HLA-DQB1/CD74/HLA-DQA2/TNFSF13B/HLA-DQB2/HLA-DMB/CD274/GBP1/NLRC5/C2/ZBP1/FGL2/CD40/C1R/HLA-DMA/CD38/RSAD2/CFI/HLA-F/HLA-B/CLC/ISG15/HLA-DOB/GBP2/SLC15A3/OAS3/CASP1/CD5L/IRF1/SAMHD1/BTN3A1/BTN1A1/PARP9/HLA-E/DDX60/SECTM1/HLA-C/NCF1/TAP2/B2M/IFI35/BTN3A3/HLA-A/AIM2/OASL/BTN3A2/USP18/ASCL2/IFIH1/CD7/IFI16/PARP14/STAT2/CFB/C3/BIRC3/BTN2A2/EREG/OAS1/JAK2/CTSS/NLRP6/ADA/PDE4B/IL15/NMI/IL12A/PIK3AP1/MICB/ZNFX1/TRIM31/LAT2/PRKCQ/TRIM21/HLA-G/BANK1/DHX58/TRAFD1/LGALS9/TNFRSF1B/EIF2AK2/IL18R1/C8G/CD55/FCER1G/PARP3/CCL5/TLR3/ADAR/ALPK1/IL20RB/RBCK1/CYLD/NR1H4/CEACAM1/RIPK2/CASP8/CD47/ZC3HAV1/PSMA1/FGL1/TRIL/TRIM56/TRIM15/OTUD4/LYN/RASGRP1/WDFY1/USP38/PRKD2/TICAM1/NLRC3/SPNS2/PLSCR1/RNF31/S100A14/NOD1/TRIM25/UBR2/USP15/POLR3G/HSPD1/IRF7 |
| GO:0009615 | response to virus | 237 | 0.72980 | 2.79010 | 1.00E-10 | 5.89E-09 | 4.73E-09 | tags=32%, list=7%, signal=31% | GBP5/CXCL9/CXCL10/IFI44L/BST2/OAS2/IFITM1/GBP1/NLRC5/ZBP1/FGL2/IFI44/CD40/APOBEC3G/IFI6/RSAD2/IFIT3/CCL22/ISG15/GBP2/MX1/STAT1/OAS3/CASP1/IRF1/SAMHD1/IFITM3/IFIT1/PARP9/DDX60/AIM2/OASL/IFIT5/IFI27/MX2/RTP4/USP18/IFIH1/IFI16/APOBEC3D/ISG20/IFIT2/STAT2/PMAIP1/IRF9/OAS1/DDX60L/DTX3L/JAK2/NLRP6/TRIM22/IL15/NMI/IL12A/GBP3/MICB/MLKL/ZNFX1/TRIM31/TRIM21/DHX58/LGALS9/HERC5/EIF2AK2/IFITM2/APOBEC3H/CCL5/TLR3/PML/MOV10/ADAR/APOBEC3F/ZC3HAV1/TRIM56/ZDHHC11/TRIM38 |
| GO:0048002 | antigen processing and presentation of peptide antigen | 50 | 0.89400 | 2.75751 | 1.00E-10 | 5.89E-09 | 4.73E-09 | tags=58%, list=4%, signal=56% | HLA-DPA1/HLA-DPB1/HLA-DOA/HLA-DRA/HLA-DQA1/HLA-DRB1/HLA-DRB5/HLA-DQB1/CD74/HLA-DQA2/HLA-DQB2/HLA-DMB/HLA-DMA/HLA-F/HLA-B/IFI30/HLA-DOB/TAP1/HLA-E/HLA-C/TAP2/B2M/HLA-A/CTSS/ERAP2/LGMN/TAPBPL/MICB/HLA-G |
| GO:0001906 | cell killing | 95 | 0.79588 | 2.74024 | 1.00E-10 | 5.89E-09 | 4.73E-09 | tags=41%, list=6%, signal=39% | GBP5/CXCL9/PLA2G2A/CXCL10/HLA-DRA/HLA-DRB1/CFH/CXCL11/GBP1/NOS2/CCL25/HLA-F/CCL22/HLA-B/GBP2/CD5L/ICAM1/HLA-E/DAO/APOL1/HLA-C/TAP2/B2M/HLA-A/C3/CXCL14/NLRP6/IL12A/GSDMB/GBP3/MICB/HLA-G/LGALS9/CXCL2/C8G/CD55/RNF19B/CEACAM1/MUC7 |
| GO:0019884 | antigen processing and presentation of exogenous antigen | 35 | 0.92818 | 2.73032 | 1.00E-10 | 5.89E-09 | 4.73E-09 | tags=66%, list=4%, signal=64% | HLA-DPA1/HLA-DPB1/HLA-DOA/HLA-DRA/HLA-DQA1/HLA-DRB1/HLA-DRB5/HLA-DQB1/CD74/HLA-DQA2/HLA-DQB2/HLA-DMB/HLA-DMA/HLA-F/IFI30/HLA-DOB/HLA-E/TAP2/B2M/HLA-A/PSME1/CTSS/LGMN |
| GO:1903900 | regulation of viral life cycle | 82 | 0.79842 | 2.68082 | 1.00E-10 | 5.89E-09 | 4.73E-09 | tags=46%, list=8%, signal=43% | HLA-DRB1/CIITA/BST2/CD74/OAS2/IFITM1/APOBEC3G/RSAD2/ISG15/LAMP3/MX1/OAS3/IFITM3/IFIT1/OASL/IFIT5/IL32/IFIH1/IFI16/APOBEC3D/ISG20/OAS1/PROX1/TRIM22/ZNFX1/TRIM31/TRIM21/LGALS9/EIF2AK2/IFITM2/APOBEC3H/CCL5/ADAR/APOBEC3F/ZC3HAV1/TRIM38/TRIM26/TRIM15 |
| GO:0050792 | regulation of viral process | 95 | 0.77729 | 2.67623 | 1.00E-10 | 5.89E-09 | 4.73E-09 | tags=43%, list=8%, signal=40% | HLA-DRB1/CIITA/BST2/CD74/OAS2/IFITM1/APOBEC3G/RSAD2/ISG15/LAMP3/MX1/STAT1/OAS3/IFITM3/IFIT1/OASL/IFIT5/IL32/IFIH1/IFI16/APOBEC3D/ISG20/OAS1/PROX1/TRIM22/ZNFX1/TRIM31/TRIM21/SP100/LGALS9/EIF2AK2/IFITM2/APOBEC3H/CCL5/ADAR/APOBEC3F/ZC3HAV1/TRIM14/TRIM38/TRIM26/TRIM15 |
| GO:0002478 | antigen processing and presentation of exogenous peptide antigen | 31 | 0.93800 | 2.66555 | 1.00E-10 | 5.89E-09 | 4.73E-09 | tags=71%, list=4%, signal=69% | HLA-DPA1/HLA-DPB1/HLA-DOA/HLA-DRA/HLA-DQA1/HLA-DRB1/HLA-DRB5/HLA-DQB1/CD74/HLA-DQA2/HLA-DQB2/HLA-DMB/HLA-DMA/HLA-F/IFI30/HLA-DOB/HLA-E/TAP2/B2M/HLA-A/CTSS/LGMN |
| GO:0048525 | negative regulation of viral process | 64 | 0.82522 | 2.65700 | 1.00E-10 | 5.89E-09 | 4.73E-09 | tags=55%, list=8%, signal=51% | CIITA/BST2/CD74/OAS2/IFITM1/APOBEC3G/RSAD2/ISG15/MX1/STAT1/OAS3/IFITM3/IFIT1/OASL/IFIT5/IL32/IFIH1/IFI16/APOBEC3D/ISG20/OAS1/PROX1/ZNFX1/TRIM31/TRIM21/SP100/EIF2AK2/IFITM2/APOBEC3H/CCL5/APOBEC3F/ZC3HAV1/TRIM14/TRIM26/TRIM15 |
| GO:0034341 | response to type II interferon | 72 | 0.80807 | 2.65559 | 1.00E-10 | 5.89E-09 | 4.73E-09 | tags=49%, list=7%, signal=46% | GBP5/HLA-DPA1/GBP4/CIITA/BST2/CD74/IFITM1/GBP1/NLRC5/NOS2/CD40/UBD/GBP2/STAT1/CASP1/IRF1/IFITM3/PARP9/GBP6/PARP14/ASS1/JAK2/IRF8/NUB1/GBP3/TRIM21/SP100/LGALS9/IFITM2/CCL5/TLR3/CITED1/CD47/KYNU/GCH1 |
| GO:0002460 | adaptive immune response based on somatic recombination of immune receptors built from immunoglobulin superfamily domains | 147 | 0.72081 | 2.63676 | 1.00E-10 | 5.89E-09 | 4.73E-09 | tags=31%, list=6%, signal=29% | SERPING1/HLA-DRA/C1S/HLA-DRB1/CD74/TNFSF13B/CD274/C2/CD40/C1R/RSAD2/CFI/HLA-F/HLA-B/CLC/ICAM1/HLA-E/SECTM1/HLA-C/TAP2/IL18BP/B2M/BTN3A3/HLA-A/BTN3A2/ASCL2/CD7/C3/JAK2/ADA/IL12A/MICB/PRKCQ/HLA-G/TNFRSF1B/IL18R1/C8G/CD55/IL6R/FCER1G/PARP3/IL20RB/SOCS3/CEACAM1/RIPK2 |
| GO:0002684 | positive regulation of immune system process | 495 | 0.64177 | 2.63082 | 1.00E-10 | 5.89E-09 | 4.73E-09 | tags=23%, list=6%, signal=23% | GBP5/SERPING1/IDO1/HLA-DPA1/HLA-DPB1/CXCL10/HLA-DOA/HLA-DRA/HLA-DQA1/C1S/HLA-DRB1/CFH/HLA-DRB5/HLA-DQB1/CD74/HLA-DQA2/TNFSF13B/HLA-DQB2/HLA-DMB/CD274/GBP1/NLRC5/C2/ZBP1/NOS2/CD40/C1R/HLA-DMA/CX3CL1/CD38/RSAD2/CFI/HLA-F/HLA-B/HLA-DOB/GBP2/SLC15A3/OAS3/CASP1/IL7/CD5L/SOCS1/ICAM1/IRF1/BTN3A1/BTN1A1/PARP9/HLA-E/DDX60/SECTM1/HLA-C/TAP2/B2M/IFI35/BTN3A3/HLA-A/AIM2/OASL/BTN3A2/IL15RA/ASCL2/IFIH1/CD7/IFI16/CFB/C3/BIRC3/BTN2A2/EREG/OAS1/CAMK1D/JAK2/CTSS/NLRP6/ADA/PDE4B/IL15/NMI/IL12A/PIK3AP1/DPP4/LGMN/MICB/MDK/CSF1/ZNFX1/TRIM31/LAT2/PRKCQ/HLA-G/BANK1/DHX58/ITGA2B/LGALS9/EIF2AK2/IL18R1/GPR68/C8G/CD55/IL6R/FCER1G/CCL5/TLR3/CSF1R/ALPK1/HK1/RBCK1/CYLD/NR1H4/CEACAM1/RIPK2/CASP8/CD47 |
| GO:0002831 | regulation of response to biotic stimulus | 276 | 0.66679 | 2.59600 | 1.00E-10 | 5.89E-09 | 4.73E-09 | tags=33%, list=10%, signal=31% | GBP5/SERPING1/HLA-DRB1/CFH/CD274/NLRC5/ZBP1/FGL2/CX3CL1/APOBEC3G/RSAD2/HLA-F/HLA-B/ISG15/GBP2/STAT1/SLC15A3/OAS3/CASP1/IRF1/SAMHD1/PARP9/HLA-E/DDX60/NCF1/IFI35/HLA-A/AIM2/OASL/USP18/IFIH1/IFI16/PARP14/STAT2/BIRC3/CARD16/EREG/OAS1/DTX3L/CTSS/NLRP6/TRIM22/IL15/NMI/IL12A/PIK3AP1/MICB/ZNFX1/TRIM31/TRIM21/HLA-G/DHX58/TRAFD1/LGALS9/HERC5/EIF2AK2/CD55/CCL5/TLR3/PML/ADAR/ALPK1/OPTN/CYLD/NR1H4/CEACAM1/APOBEC3F/RIPK2/CASP8/ZC3HAV1/TRIL/TRIM56/ZDHHC11/TRIM38/TRIM15/OTUD4/LYN/RASGRP1/WDFY1/USP38/TICAM1/NLRC3/PLSCR1/RNF31/S100A14/NOD1/TRIM25/USP15/POLR3G/HSPD1/IRF7 |
| GO:0045069 | regulation of viral genome replication | 50 | 0.84115 | 2.59450 | 1.00E-10 | 5.89E-09 | 4.73E-09 | tags=54%, list=7%, signal=51% | BST2/OAS2/IFITM1/APOBEC3G/RSAD2/ISG15/MX1/OAS3/IFITM3/IFIT1/OASL/IFIT5/IFIH1/IFI16/APOBEC3D/ISG20/OAS1/PROX1/ZNFX1/EIF2AK2/IFITM2/APOBEC3H/CCL5/ADAR/APOBEC3F/ZC3HAV1/TRIM38 |
| GO:0002504 | antigen processing and presentation of peptide or polysaccharide antigen via MHC class II | 26 | 0.93422 | 2.59172 | 1.00E-10 | 5.89E-09 | 4.73E-09 | tags=69%, list=4%, signal=67% | HLA-DPA1/HLA-DPB1/HLA-DOA/HLA-DRA/HLA-DQA1/HLA-DRB1/HLA-DRB5/HLA-DQB1/CD74/HLA-DQA2/HLA-DQB2/HLA-DMB/HLA-DMA/IFI30/HLA-DOB/B2M/CTSS/LGMN |

ES, enrichment Score; NES, normalized enrichment Score

6. GSEA (Gene Set Enrichment Analysis): Canonical Pathways (C2)

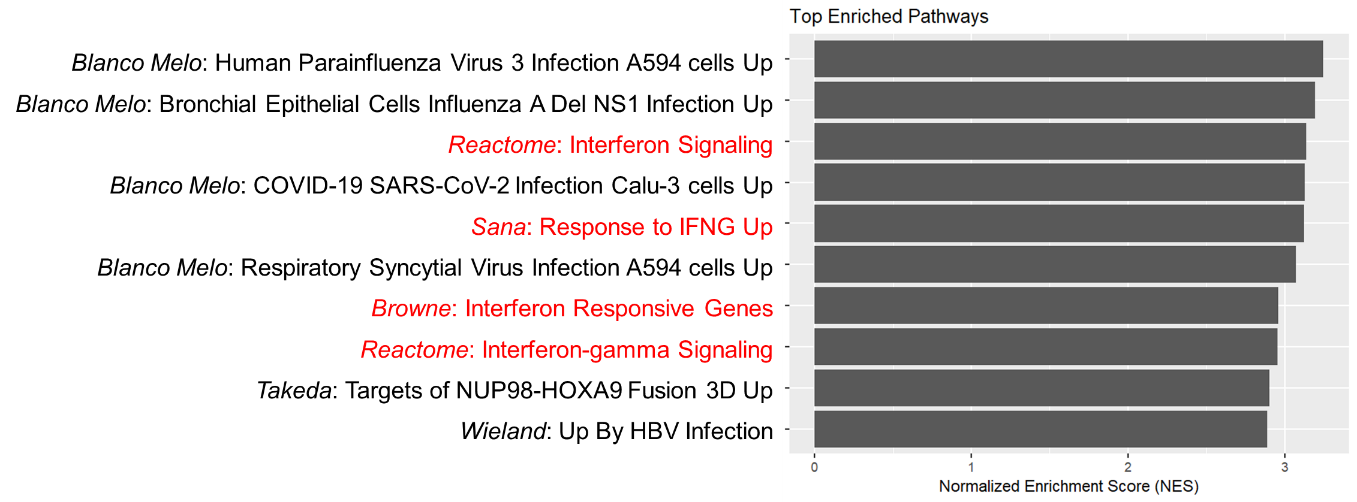

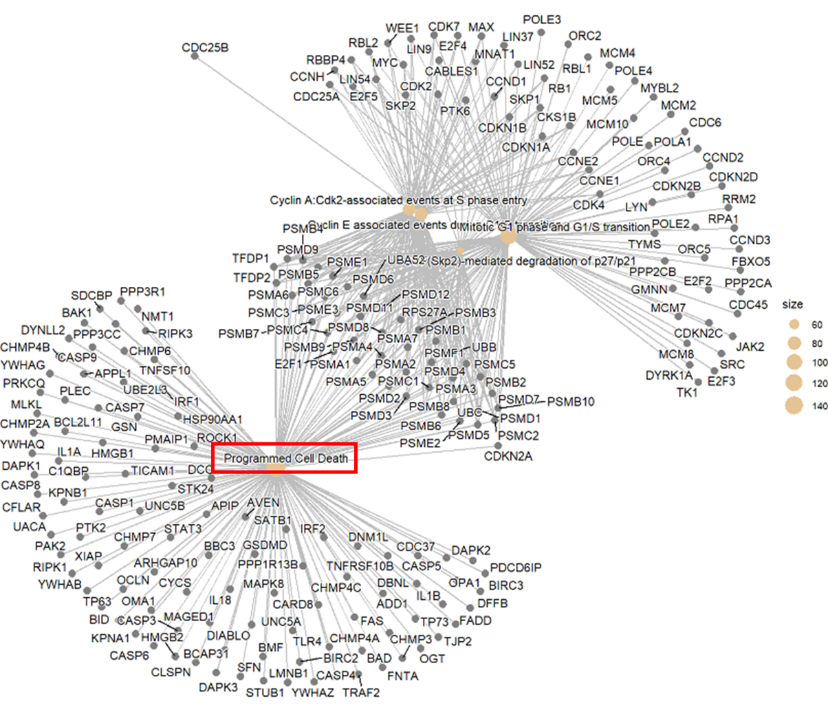
Supplementary Material S4. REACTOME Network Plot

Supplementary Material S5. Morphology of intestinal organoids and number of isolated intestinal epithelial cells used for single-cell RNA-sequencing

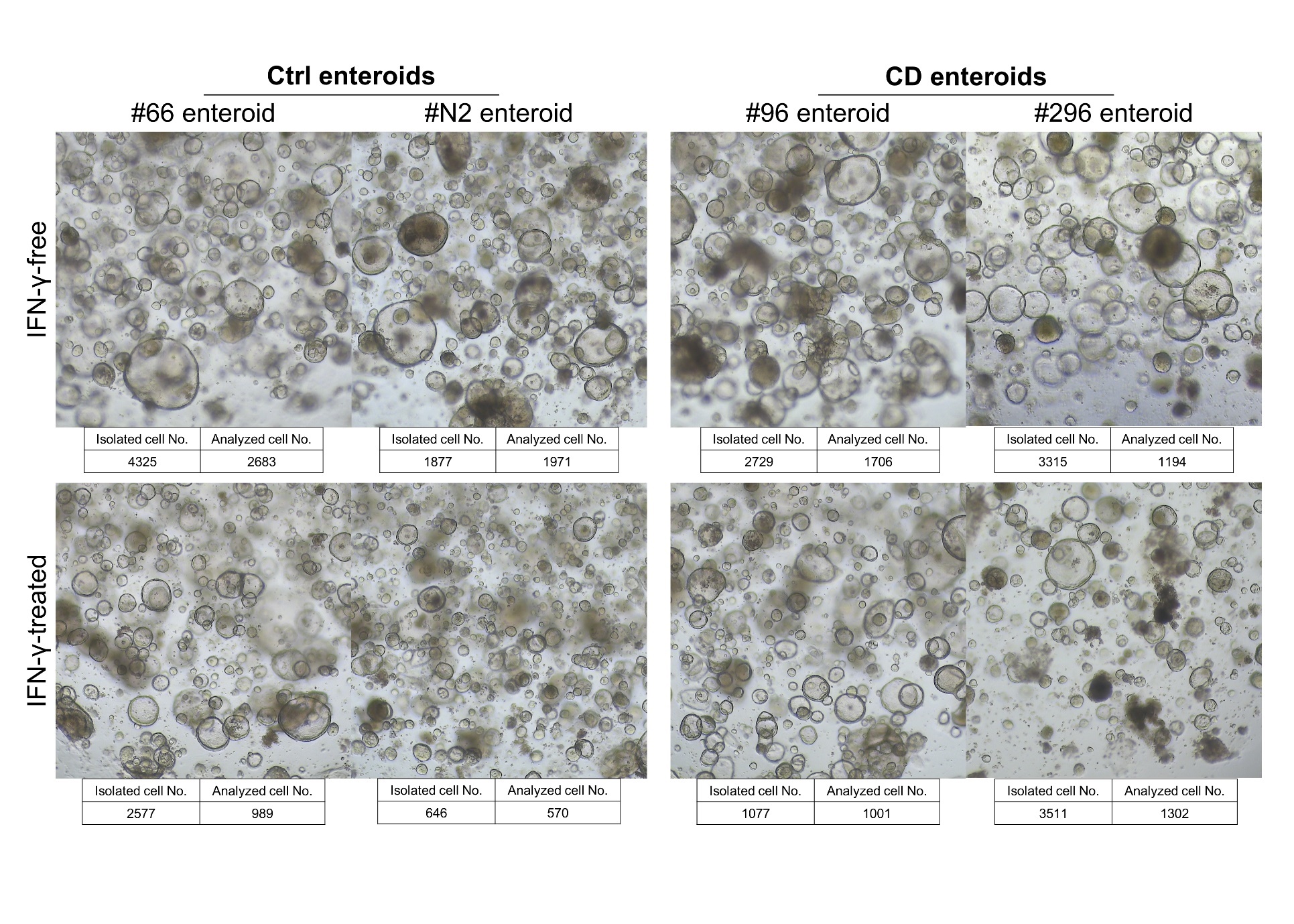

Supplementary Material S6. Differentially expressed genes in cell clusters. (A) Top 10 differentially expressed genes in each cluster. (B) Top 10 gene ontology categories enriched with differentially expressed genes in each cluster identified by Gene Set Enrichment Analysis (GSEA)

(A) Top 10 differentially expressed genes (positively differentially expressed gene only, minimum percentage of cells in cluster > 25%, and logFC > 0.25) in each cluster

| Cluster | Gene symbol | Average log2 fold change (logFC) | Percentage of cells where the gene is detected in the cluster | Percentage of cells where the gene is detected on average in the other clusters | *p*-value | Adjusted *p*-value, (Bonferroni correction) |
| --- | --- | --- | --- | --- | --- | --- |
| 0 | MT1X | 1.10 | 85.0% | 64.3% | 8.78E-176 | 1.60E-171 |
|  | CLIC3 | 0.60 | 74.9% | 50.1% | 8.59E-101 | 1.56E-96 |
|  | TFF1 | 0.54 | 98.6% | 91.1% | 2.09E-113 | 3.80E-109 |
|  | SLC6A8 | 0.54 | 86.0% | 58.6% | 2.05E-150 | 3.73E-146 |
|  | SLURP2 | 0.45 | 27.5% | 13.3% | 9.62E-51 | 1.75E-46 |
|  | CA9 | 0.44 | 71.9% | 44.1% | 1.31E-93 | 2.39E-89 |
|  | TFF2 | 0.44 | 87.6% | 68.7% | 4.62E-85 | 8.39E-81 |
|  | NDRG1 | 0.39 | 75.2% | 49.2% | 4.44E-102 | 8.07E-98 |
|  | ANKRD37 | 0.37 | 36.2% | 18.0% | 3.38E-65 | 6.15E-61 |
|  | DUOXA2 | 0.36 | 92.4% | 70.8% | 4.68E-123 | 8.50E-119 |
| 1 | UBD | 0.57 | 64.7% | 51.8% | 9.54E-42 | 1.74E-37 |
|  | MT-ATP8 | 0.43 | 82.3% | 61.1% | 5.08E-74 | 9.24E-70 |
|  | PI3 | 0.42 | 66.5% | 48.7% | 1.43E-41 | 2.59E-37 |
| 2 | S100P | 2.64 | 86.3% | 63.3% | 4.94E-221 | 8.98E-217 |
|  | GAS5 | 2.30 | 88.4% | 59.5% | 3.85E-245 | 7.01E-241 |
|  | RPS2 | 2.10 | 99.9% | 90.7% | 0 | 0 |
|  | RPL21 | 2.07 | 100.0% | 89.4% | 0 | 0 |
|  | RPL10A | 2.06 | 99.9% | 85.8% | 0 | 0 |
|  | RPS8 | 2.06 | 100.0% | 97.0% | 0 | 0 |
|  | RPL17 | 2.03 | 100.0% | 95.0% | 0 | 0 |
|  | RPS16 | 2.02 | 100.0% | 90.6% | 0 | 0 |
|  | RPL37A | 2.02 | 99.9% | 88.2% | 0 | 0 |
|  | SNHG8 | 2.00 | 96.7% | 67.9% | 0 | 0 |
| 4 | MT1X | 2.42 | 91.6% | 64.7% | 1.50E-255 | 2.73E-251 |
|  | SLC6A8 | 1.90 | 96.4% | 59.0% | 0 | 0 |
|  | ERO1A | 1.88 | 99.3% | 74.7% | 0 | 0 |
|  | TFF1 | 1.87 | 99.5% | 91.4% | 1.01E-235 | 1.84E-231 |
|  | DUOXA2 | 1.78 | 97.9% | 71.3% | 2.47E-302 | 4.49E-298 |
|  | SCD | 1.78 | 97.3% | 74.4% | 5.66E-274 | 1.03E-269 |
|  | NDRG1 | 1.70 | 88.8% | 49.2% | 3.58E-272 | 6.52E-268 |
|  | IGFBP3 | 1.69 | 75.0% | 53.7% | 1.69E-89 | 3.07E-85 |
|  | TFF2 | 1.62 | 94.6% | 69.0% | 9.84E-219 | 1.79E-214 |
|  | EGLN3 | 1.58 | 94.9% | 55.1% | 0 | 0 |
| 5 | ALDH1A3 | 3.21 | 95.5% | 41.7% | 0 | 0 |
|  | PLAAT4 | 3.07 | 95.9% | 74.0% | 6.90E-171 | 1.25E-166 |
|  | CXCL5 | 3.07 | 77.2% | 58.7% | 7.72E-114 | 1.40E-109 |
|  | MMP7 | 2.73 | 95.9% | 75.5% | 7.80E-107 | 1.42E-102 |
|  | CCND1 | 2.65 | 99.2% | 55.6% | 0 | 0 |
|  | IFI6 | 2.60 | 90.2% | 61.4% | 2.05E-110 | 3.73E-106 |
|  | CCND2 | 2.31 | 45.7% | 21.9% | 1.12E-65 | 2.04E-61 |
|  | CCN2 | 2.20 | 79.5% | 28.2% | 4.49E-238 | 8.16E-234 |
|  | DCBLD2 | 2.18 | 83.6% | 27.4% | 1.14E-307 | 2.06E-303 |
|  | KRT7 | 2.13 | 99.9% | 90.0% | 3.44E-257 | 6.25E-253 |
| 7 | NEAT1 | 3.36 | 99.1% | 84.4% | 4.82E-218 | 8.76E-214 |
|  | MALAT1 | 3.19 | 99.6% | 91.6% | 3.07E-215 | 5.58E-211 |
|  | MTRNR2L12 | 2.09 | 76.1% | 23.3% | 4.86E-215 | 8.83E-211 |
|  | WSB1 | 1.87 | 90.8% | 42.2% | 3.36E-205 | 6.11E-201 |
|  | XIST | 1.82 | 46.9% | 20.3% | 1.37E-72 | 2.50E-68 |
|  | KCNQ1OT1 | 1.81 | 69.7% | 13.7% | 3.34E-296 | 6.08E-292 |
|  | PARP14 | 1.77 | 85.7% | 39.0% | 7.03E-177 | 1.28E-172 |
|  | DST | 1.66 | 91.3% | 54.0% | 4.33E-156 | 7.87E-152 |
|  | VMP1 | 1.54 | 88.7% | 47.6% | 2.20E-170 | 4.01E-166 |
|  | CYP3A5 | 1.41 | 89.6% | 47.4% | 2.26E-155 | 4.11E-151 |
| 8 | MT1G | 3.30 | 36.8% | 12.2% | 1.92E-64 | 3.50E-60 |
|  | MT2A | 2.57 | 41.8% | 22.4% | 6.80E-32 | 1.24E-27 |
|  | PHGR1 | 2.54 | 99.6% | 95.4% | 2.01E-164 | 3.65E-160 |
|  | KRT20 | 2.41 | 84.2% | 31.5% | 3.15E-210 | 5.72E-206 |
|  | FABP2 | 2.39 | 84.8% | 28.5% | 1.73E-227 | 3.15E-223 |
|  | PIGR | 2.38 | 99.8% | 75.0% | 1.54E-219 | 2.80E-215 |
|  | REG4 | 2.28 | 93.8% | 76.7% | 4.46E-109 | 8.11E-105 |
|  | ALDOB | 2.27 | 90.5% | 37.9% | 1.34E-213 | 2.44E-209 |
|  | RBP2 | 2.20 | 55.6% | 18.5% | 5.98E-107 | 1.09E-102 |
|  | LGALS3 | 2.05 | 100.0% | 97.2% | 3.13E-186 | 5.70E-182 |
| 9 | PLA2G2A | 0.26 | 29.5% | 14.6% | 2.71E-14 | 4.92E-10 |
| 10 | UBE2C | 3.22 | 92.1% | 3.8% | 0 | 0 |
|  | CKS2 | 3.06 | 99.7% | 42.3% | 9.07E-259 | 1.65E-254 |
|  | PTTG1 | 2.94 | 94.9% | 8.6% | 0 | 0 |
|  | HIST1H4C | 2.86 | 84.1% | 36.4% | 8.31E-150 | 1.51E-145 |
|  | H2AFZ | 2.84 | 100.0% | 70.2% | 1.03E-215 | 1.87E-211 |
|  | TUBA1B | 2.69 | 100.0% | 58.8% | 9.29E-222 | 1.69E-217 |
|  | PCLAF | 2.66 | 98.9% | 13.1% | 0 | 0 |
|  | UBE2S | 2.64 | 97.2% | 24.5% | 0 | 0 |
|  | HMGB2 | 2.48 | 94.6% | 12.0% | 0 | 0 |
|  | CENPW | 2.37 | 99.4% | 27.7% | 1.42E-307 | 2.59E-303 |
| 11 | APOC3 | 5.76 | 77.8% | 16.4% | 5.01E-226 | 9.11E-222 |
|  | APOA1 | 5.70 | 81.4% | 16.3% | 3.81E-251 | 6.92E-247 |
|  | APOA4 | 5.25 | 71.2% | 9.8% | 2.35E-288 | 4.27E-284 |
|  | RBP2 | 4.72 | 89.5% | 18.1% | 3.16E-283 | 5.75E-279 |
|  | ANPEP | 3.57 | 88.9% | 11.1% | 0 | 0 |
|  | SELENOP | 3.20 | 84.6% | 11.8% | 0 | 0 |
|  | ALDOB | 3.20 | 98.4% | 38.5% | 6.06E-209 | 1.10E-204 |
|  | MT1G | 3.01 | 50.3% | 12.2% | 5.00E-100 | 9.09E-96 |
|  | FABP2 | 2.85 | 95.1% | 29.1% | 7.56E-213 | 1.38E-208 |
|  | APOB | 2.67 | 70.6% | 4.4% | 0 | 0 |
| 12 | APOC3 | 2.67 | 77.6% | 16.6% | 2.10E-183 | 3.81E-179 |
|  | APOA1 | 2.67 | 77.2% | 16.6% | 4.45E-181 | 8.08E-177 |
|  | RBP2 | 2.24 | 90.7% | 18.3% | 1.38E-230 | 2.51E-226 |
|  | APOA4 | 2.16 | 63.8% | 10.2% | 4.83E-182 | 8.79E-178 |
|  | ANPEP | 1.64 | 80.2% | 11.6% | 5.15E-248 | 9.36E-244 |
|  | MT1G | 1.61 | 54.1% | 12.3% | 1.09E-100 | 1.99E-96 |
|  | SELENOP | 1.14 | 69.4% | 12.4% | 4.03E-167 | 7.32E-163 |
|  | PRAP1 | 1.01 | 92.5% | 69.4% | 5.09E-46 | 9.25E-42 |
|  | MT1H | 0.99 | 26.1% | 3.5% | 2.54E-79 | 4.62E-75 |
|  | ALDOB | 0.89 | 87.3% | 39.0% | 8.93E-76 | 1.62E-71 |
| 14 | IGFBP3 | 1.51 | 90.8% | 55.0% | 9.01E-57 | 1.64E-52 |
|  | CCN2 | 1.50 | 78.9% | 30.4% | 1.32E-71 | 2.41E-67 |
|  | SLC2A3 | 1.49 | 43.9% | 4.0% | 1.48E-174 | 2.70E-170 |
|  | LOXL4 | 1.40 | 78.9% | 24.3% | 1.39E-93 | 2.53E-89 |
|  | HSPG2 | 1.33 | 99.6% | 63.8% | 1.56E-76 | 2.85E-72 |
|  | LAMC2 | 1.26 | 89.5% | 47.7% | 1.51E-62 | 2.74E-58 |
|  | CDA | 1.19 | 96.5% | 66.8% | 1.47E-60 | 2.68E-56 |
|  | F3 | 1.19 | 99.6% | 67.4% | 5.12E-62 | 9.31E-58 |
|  | SLC2A1 | 1.19 | 93.9% | 54.8% | 2.05E-62 | 3.72E-58 |
|  | CST6 | 1.16 | 68.9% | 21.1% | 8.50E-79 | 1.55E-74 |
| 15 | UBD | 3.14 | 89.6% | 53.0% | 1.29E-54 | 2.35E-50 |
|  | LTB | 2.81 | 74.0% | 16.1% | 1.01E-109 | 1.84E-105 |
|  | CLDN18 | 2.42 | 61.3% | 10.0% | 1.46E-114 | 2.65E-110 |
|  | PLAU | 2.41 | 79.8% | 12.8% | 1.55E-168 | 2.83E-164 |
|  | CCL20 | 2.30 | 63.6% | 29.2% | 2.26E-31 | 4.12E-27 |
|  | CXCL8 | 2.27 | 78.0% | 30.4% | 2.99E-59 | 5.44E-55 |
|  | SERPINB7 | 2.05 | 52.0% | 8.4% | 5.73E-95 | 1.04E-90 |
|  | CXCL1 | 2.02 | 86.1% | 48.8% | 2.68E-46 | 4.87E-42 |
|  | ISG20 | 2.00 | 93.6% | 47.4% | 3.75E-78 | 6.81E-74 |
|  | SAA1 | 2.00 | 52.6% | 21.4% | 1.70E-28 | 3.10E-24 |
| 16 | ALDH1A3 | 1.38 | 84.1% | 44.8% | 1.16E-16 | 2.10E-12 |
|  | CCND2 | 1.33 | 53.7% | 23.1% | 2.32E-13 | 4.23E-09 |
|  | MT-ND3 | 1.33 | 100.0% | 92.5% | 7.27E-17 | 1.32E-12 |
|  | GAS5 | 1.33 | 95.1% | 62.9% | 2.93E-25 | 5.32E-21 |
|  | S100P | 1.32 | 92.7% | 66.0% | 4.18E-19 | 7.60E-15 |
|  | CCND1 | 1.29 | 87.8% | 58.1% | 6.95E-14 | 1.26E-09 |
|  | DDX21 | 1.27 | 90.2% | 35.2% | 9.04E-36 | 1.64E-31 |
|  | EIF5B | 1.23 | 98.8% | 47.5% | 1.68E-33 | 3.05E-29 |
|  | ENC1 | 1.23 | 75.6% | 41.2% | 2.86E-15 | 5.19E-11 |
|  | AC020656.1 | 1.21 | 95.1% | 43.3% | 6.37E-32 | 1.16E-27 |
| * If no features pass the logFC threshold, DEGs were not identified or a limited number of DEGs were identified. | | | | | | |

(B) Top 10 gene ontology categories enriched in each cluster identified by Gene Set Enrichment Analysis (GSEA) of Differentially Expressed Genes (DEGs)

| Cluster | Gene ontology | *p*-value | FDR *q*-value |
| --- | --- | --- | --- |
| 0 | GOMF_Lyase_Activity [200] | 1.35 e^-6^ | 1.1 e^-2^ |
|  | GOMF_Carbon_Carbon_Lyase_Activity [52] | 1.37 e^-6^ | 1.1 e^-2^ |
|  | HP_Porokeratosis [8] | 4.69 e^-6^ | 1.93 e^-2^ |
|  | GOBP_Fructose_1_6_Bisphosphate_Metabolic_Process [9] | 6.03 e^-6^ | 1.93 e^-2^ |
|  | GOMF_Aldehyde_Lyase_Activity [9] | 6.03 e^-6^ | 1.93 e^-2^ |
|  | GOBP_Fructose_Metabolic_Process [14] | 1.52 e^-5^ | 4.06 |
| 2 | GOCC_Cytosolic_Ribosome [118] | 1.72 e^-172^ | 2.75 e^-168^ |
|  | GOBP_Cytoplasmic_Translation [156] | 2.39 e^-166^ | 1.92 e^-162^ |
|  | GOMF_Structural_Constituent_Of_Ribosome [169] | 9.34 e^-154^ | 4.98 e^-150^ |
|  | GOCC_Ribosomal_Subunit [188] | 1.53 e^-152^ | 6.13 e^-149^ |
|  | GOCC_Ribosome [239] | 4.53 e^-149^ | 1.45 e^-145^ |
|  | GOBP_Peptide_Biosynthetic_Process [866] | 6.64 e^-115^ | 1.77 e^-111^ |
|  | GOBP_Peptide_Metabolic_Process [997] | 6.95 e^-110^ | 1.59 e^-106^ |
|  | GOBP_Amide_Biosynthetic_Process [1004] | 1.23 e^-109^ | 2.46 e^-106^ |
|  | GOMF_Structural_Molecule_Activity [809] | 1.17 e^-101^ | 2.07 e^-98^ |
|  | GOBP_Amide_Metabolic_Process [1301] | 1.72 e^-100^ | 2.76 e^-97^ |
| 4 | GOBP_Sterol_Biosynthetic_Process [68] | 7.28 e^-16^ | 1.06 e^-11^ |
|  | GOBP_Small_Molecule_Metabolic_Process [1871] | 1.32 e^-15^ | 1.06 e^-11^ |
|  | GOBP_Small_Molecule_Biosynthetic_Process [608] | 6.06 e^-14^ | 3.23 e^-10^ |
|  | GOBP_Sterol_Metabolic_Process [155] | 9.58 e^-14^ | 3.84 e^-10^ |
|  | GOBP_Organic_Hydroxy_Compound_Metabolic_Process [567] | 3.08 e^-13^ | 9.86 e^-10^ |
|  | GOBP_Alcohol_Metabolic_Process [373] | 4.37 e^-12^ | 1.17 e^-8^ |
|  | GOBP_Steroid_Metabolic_Process [327] | 1.69 e^-11^ | 3.85 e^-8^ |
|  | GOBP_Steroid_Biosynthetic_Process [185] | 2.03 e^-11^ | 4.05 e^-8^ |
|  | GOCC_Nuclear_Outer_Membrane_Endoplasmic_Reticulum_Membrane_Network [1212] | 4.38 e^-11^ | 7.26 e^-8^ |
|  | GOBP_Negative_Regulation_Of_Molecular_Function [792] | 4.54 e^-11^ | 7.26 e^-8^ |
| 5 | GOBP_Apoptotic_Process [1965] | 5.75 e^-17^ | 9.2 e^-13^ |
|  | GOCC_Anchoring_Junction [901] | 1.3 e^-16^ | 1.04 e^-12^ |
|  | GOBP_Regulation_Of_Programmed_Cell_Death [1563] | 1.35 e^-14^ | 7.2 e^-11^ |
|  | GOBP_Cell_Motility [1813] | 4.55 e^-14^ | 1.53 e^-10^ |
|  | GOBP_Regulation_Of_Catalytic_Activity [1497] | 4.78 e^-14^ | 1.53 e^-10^ |
|  | GOBP_Circulatory_System_Development [1224] | 6.64 e^-14^ | 1.77 e^-10^ |
|  | GOMF_Cell_Adhesion_Molecule_Binding [558] | 2.42 e^-13^ | 5.53 e^-10^ |
|  | GOBP_Regulation_Of_Multicellular_Organismal_Development [1484] | 3.54 e^-13^ | 7.09 e^-10^ |
|  | GOBP_Anatomical_Structure_Formation_Involved_In_Morphogenesis [1259] | 1.11 e^-12^ | 1.97 e^-9^ |
|  | GOMF_Cytoskeletal_Protein_Binding [991] | 1.38 e^-12^ | 2.22 e^-9^ |
| 7 | HP_Centrocecal_Scotoma [13] | 6.7 e^-19^ | 1.07 e^-14^ |
|  | HP_Leber_Optic_Atrophy [10] | 2.98 e^-17^ | 2.38 e^-13^ |
|  | HP_Mitochondrial_Respiratory_Chain_Defects [20] | 6.47 e^-17^ | 3.45 e^-13^ |
|  | HP_Central_Retinal_Vessel_Vascular_Tortuosity [12] | 1.96 e^-16^ | 7.83 e^-13^ |
|  | HP_Abnormal_Cardiovascular_System_Physiology [1408] | 7.51 e^-16^ | 2.41 e^-12^ |
|  | HP_Retinal_Telangiectasia [27] | 1.13 e^-15^ | 3 e^-12^ |
|  | HP_Mitochondrial_Inheritance [29] | 2.17 e^-15^ | 4.34 e^-12^ |
|  | HP_Vascular_Tortuosity [29] | 2.17 e^-15^ | 4.34 e^-12^ |
|  | HP_Abnormality_Of_Cardiovascular_System_Electrophysiology [542] | 6.61 e^-15^ | 1.18 e^-11^ |
|  | HP_Slow_Decrease_In_Visual_Acuity [34] | 9.09 e^-15^ | 1.46 e^-11^ |
| 8 | GOCC_Inner_Mitochondrial_Membrane_Protein_Complex [158] | 1.95 e^-47^ | 3.12 e^-43^ |
|  | GOBP_Oxidative_Phosphorylation [148] | 4.69 e^-44^ | 3.75 e^-40^ |
|  | GOCC_Mitochondrial_Protein_Containing_Complex [300] | 1.54 e^-42^ | 7.89 e^-39^ |
|  | GOBP_Aerobic_Respiration [197] | 1.97 e^-42^ | 7.89 e^-39^ |
|  | GOBP_Cellular_Respiration [243] | 9.69 e^-40^ | 3.1 e^-36^ |
|  | GOBP_Generation_Of_Precursor_Metabolites_And_Energy [511] | 7.86 e^-37^ | 2.1 e^-33^ |
|  | GOCC_Organelle_Inner_Membrane [553] | 9.95 e^-36^ | 2.28 e^-32^ |
|  | GOBP_Energy_Derivation_By_Oxidation_Of_Organic_Compounds [337] | 1.25 e^-35^ | 2.51 e^-32^ |
|  | GOCC_Mitochondrial_Envelope [804] | 1.96 e^-33^ | 3.34 e^-30^ |
|  | GOBP_Atp_Synthesis_Coupled_Electron_Transport [102] | 2.29 e^-33^ | 3.34 e^-30^ |
| 10 | GOBP_Cell_Cycle [1841] | 1.12 e^-29^ | 1.8 e^-25^ |
|  | GOBP_Cell_Cycle_Process [1323] | 8.69 e^-29^ | 6.95 e^-25^ |
|  | GOBP_Mitotic_Cell_Cycle [932] | 1.44 e^-28^ | 7.66 e^-25^ |
|  | GOBP_Chromosome_Organization [630] | 5.41 e^-28^ | 2.16 e^-24^ |
|  | GOBP_Cell_Division [639] | 5.28 e^-25^ | 1.69 e^-21^ |
|  | GOBP_Chromosome_Segregation [441] | 1.4 e^-24^ | 3.74 e^-21^ |
|  | GOBP_Organelle_Fission [491] | 1.61 e^-23^ | 3.69 e^-20^ |
|  | GOBP_Dna_Metabolic_Process [1037] | 2.42 e^-23^ | 4.84 e^-20^ |
|  | GOBP_Mitotic_Cell_Cycle_Process [775] | 6.94 e^-23^ | 1.23 e^-19^ |
|  | GOBP_Mitotic_Nuclear_Division [277] | 4.47 e^-21^ | 7.16 e^-18^ |
| 11 | GOBP_Lipid_Metabolic_Process [1407] | 7.44 e^-20^ | 1.19 e^-15^ |
|  | GOBP_Small_Molecule_Metabolic_Process [1871] | 1.52 e^-18^ | 1.22 e^-14^ |
|  | GOBP_Cellular_Lipid_Metabolic_Process [1005] | 7.17 e^-18^ | 3.83 e^-14^ |
|  | GOCC_Brush_Border [108] | 2.79 e^-17^ | 1.12 e^-13^ |
|  | GOBP_Intestinal_Absorption [41] | 4.37 e^-16^ | 1.4 e^-12^ |
|  | GOCC_Cluster_Of_Actin_Based_Cell_Projections [165] | 5.08 e^-15^ | 1.35 e^-11^ |
|  | GOBP_Organic_Acid_Metabolic_Process [980] | 9.01 e^-15^ | 2.06 e^-11^ |
|  | GOBP_Hormone_Metabolic_Process [241] | 1.7 e^-14^ | 3.04 e^-11^ |
|  | GOCC_Apical_Part_Of_Cell [471] | 1.81 e^-14^ | 3.04 e^-11^ |
|  | GOBP_Digestion [134] | 1.9 e^-14^ | 3.04 e^-11^ |
| 12 | GOMF_Lipid_Binding [852] | 2.87 e^-10^ | 2.19 e^-6^ |
|  | GOBP_Response_To_Zinc_Ion [51] | 3.29 e^-10^ | 2.19 e^-6^ |
|  | GOBP_Detoxification_Of_Copper_Ion [16] | 4.1 e^-10^ | 2.19 e^-6^ |
|  | GOCC_Triglyceride_Rich_Plasma_Lipoprotein_Particle [20] | 1.09 e^-9^ | 3.76 e^-6^ |
|  | GOBP_Detoxification [158] | 1.59 e^-9^ | 3.76 e^-6^ |
|  | GOBP_Detoxification_Of_Inorganic_Compound [22] | 1.64 e^-9^ | 3.76 e^-6^ |
|  | GOBP_Intestinal_Lipid_Absorption [22] | 1.64 e^-9^ | 3.76 e^-6^ |
|  | GOBP_Chemical_Homeostasis [1037] | 1.9 e^-9^ | 3.8 e^-6^ |
|  | GOBP_Lipid_Localization [503] | 2.17 e^-9^ | 3.82 e^-6^ |
|  | GOBP_Cellular_Response_To_Zinc_Ion [24] | 2.38 e^-9^ | 3.82 e^-6^ |
| 14 | GOCC_Collagen_Containing_Extracellular_Matrix [430] | 1.89 e^-23^ | 3.03 e^-19^ |
|  | GOCC_External_Encapsulating_Structure [564] | 6.68 e^-21^ | 5.35 e^-17^ |
|  | GOBP_Cell_Adhesion [1531] | 6.11 e^-20^ | 3.26 e^-16^ |
|  | GOCC_Cell_Substrate_Junction [434] | 4.63 e^-19^ | 1.85 e^-15^ |
|  | GOCC_Anchoring_Junction [901] | 6.03 e^-19^ | 1.93 e^-15^ |
|  | GOBP_Cell_Motility [1813] | 6.45 e^-18^ | 1.72 e^-14^ |
|  | GOBP_Anatomical_Structure_Formation_Involved_In_Morphogenesis [1259] | 8.22 e^-18^ | 1.88 e^-14^ |
|  | GOMF_Extracellular_Matrix_Structural_Constituent [166] | 1.32 e^-16^ | 2.65 e^-13^ |
|  | GOMF_Cell_Adhesion_Molecule_Binding [558] | 8.82 e^-16^ | 1.48 e^-12^ |
|  | HP_Hypoplastic_Dermoepidermal_Hemidesmosomes [7] | 9.27 e^-16^ | 1.48 e^-12^ |
| 15 | GOBP_Biological_Process_Involved_In_Interspecies_Interaction_Between_Organisms [1759] | 4.7 e^-27^ | 7.52 e^-23^ |
|  | GOBP_Cell_Adhesion [1531] | 1.98 e^-24^ | 1.59 e^-20^ |
|  | GOBP_Defense_Response [1855] | 8.8 e^-22^ | 4.7 e^-18^ |
|  | GOBP_Locomotion [1335] | 1.32 e^-21^ | 5.28 e^-18^ |
|  | GOBP_Cell_Motility [1813] | 5.76 e^-20^ | 1.84 e^-16^ |
|  | GOMF_Signaling_Receptor_Binding [1495] | 3.81 e^-19^ | 9.28 e^-16^ |
|  | GOBP_Defense_Response_To_Other_Organisnism [1235] | 4.06 e^-19^ | 9.28 e^-16^ |
|  | GOBP_Response_To_Bacterium [742] | 2.26 e^-18^ | 4.52 e^-15^ |
|  | GOBP_Wound_Healing [425] | 7.65 e^-18^ | 1.36 e^-14^ |
|  | GOCC_Anchoring_Junction [901] | 9.09 e^-18^ | 1.46 e^-14^ |
| 16 | HP_Abnormality_Of_The_Upper_Urinary_Tract [1443] | 1.64 e^-16^ | 1.65 e^-12^ |
|  | GOCC_Cell_Substrate_Junction [434] | 2.06 e^-16^ | 1.65 e^-12^ |
|  | GOMF_Protein_Containing_Complex_Binding [1762] | 1.81 e^-14^ | 9.63 e^-11^ |
|  | GOMF_Cadherin_Binding [335] | 4.57 e^-14^ | 1.83 e^-10^ |
|  | GOBP_Cytoplasmic_Translation [156] | 9.18 e^-14^ | 2.48 e^-10^ |
|  | HP_Abnormal_Renal_Morphology [1118] | 9.3 e^-14^ | 2.48 e^-10^ |
|  | GOBP_Peptide_Metabolic_Process [997] | 1.17 e^-13^ | 2.68 e^-10^ |
|  | GOCC_Cytosolic_Ribosome [118] | 2 e^-13^ | 3.65 e^-10^ |
|  | GOMF_Cell_Adhesion_Molecule_Binding [558] | 2.05 e^-13^ | 3.65 e^-10^ |
|  | GOBP_Peptide_Biosynthetic_Process [866] | 1.27 e^-12^ | 2.03 e^-9^ |

Supplementary material S7. Epithelial lineage-specific marker gene expression in each cluster of single-cell RNA sequencing. (Bold indicates the epithelial cell lineage marker in each single-cell cluster)

| Cluster | Gene symbol | Epithelial cell lineage | Average log2 fold change | Percentage of cells where the gene is detected in the cluster | Percentage of cells where the gene is detected on average in the other clusters | *p*-value | Adjusted *p*-value, (Bonferroni correction) |
| --- | --- | --- | --- | --- | --- | --- | --- |
| 0 | RNF186 | Enterocytes | 0.3004778 | 50.3% | 28.7% | 1.57E-64 | 2.85E-60 |
| 0 | DMBT1 | Enterocytes | 0.2882969 | 65.2% | 45.7% | 3.46E-55 | 6.30E-51 |
| 2 | GPX2 | Paneth cells | 1.8443475 | 100.0% | 90.8% | 0.00E+00 | 0.00E+00 |
| 2 | **OLFM4** | **Crypt cells** | 1.7992902 | 87.6% | 63.0% | 7.23E-185 | 1.31E-180 |
| 2 | LYZ | Paneth cells | 1.3852443 | 99.9% | 96.0% | 0.00E+00 | 0.00E+00 |
| 2 | TNFAIP2 | Microfold cells | 1.3478442 | 84.7% | 64.1% | 3.95E-131 | 7.19E-127 |
| 2 | PRSS1 | Paneth cells | 1.3334085 | 43.0% | 15.7% | 5.09E-153 | 9.25E-149 |
| 2 | AGR2 | Paneth cells | 1.3140263 | 99.8% | 89.5% | 0.00E+00 | 0.00E+00 |
| 2 | GSTM3 | Enterocytes | 1.2738238 | 92.7% | 58.4% | 0.00E+00 | 0.00E+00 |
| 2 | KRTCAP3 | Enterocytes | 1.271556 | 93.2% | 51.5% | 0.00E+00 | 0.00E+00 |
| 2 | PPARG | Enterocytes | 1.2666207 | 82.1% | 45.9% | 2.16E-269 | 3.92E-265 |
| 2 | KLF5 | Enterocytes | 1.2602163 | 87.1% | 55.7% | 1.54E-233 | 2.79E-229 |
| 2 | CD55 | Enterocytes | 1.2365755 | 87.7% | 67.4% | 1.13E-136 | 2.05E-132 |
| 2 | ANXA5 | Microfold cells | 1.2360449 | 96.0% | 68.6% | 1.04E-304 | 1.89E-300 |
| 2 | KLF3 | Tuft cells | 1.2147825 | 74.0% | 43.5% | 3.42E-175 | 6.22E-171 |
| 2 | G0S2 | Enterocytes | 1.2042635 | 35.5% | 12.4% | 3.10E-129 | 5.64E-125 |
| 2 | CDC42EP5 | Enterocytes | 1.1910175 | 99.7% | 84.9% | 7.31E-304 | 1.33E-299 |
| 2 | ELF3 | Enterocytes | 1.1852043 | 97.7% | 81.1% | 4.80E-215 | 8.73E-211 |
| 2 | CD24 | Paneth cells | 1.1762028 | 99.6% | 89.4% | 9.15E-293 | 1.66E-288 |
| 2 | TRNP1 | Enteroendocrine cells | 1.1599809 | 83.8% | 49.4% | 8.26E-237 | 1.50E-232 |
| 2 | KRT7 | Goblet cells | 1.1369919 | 99.5% | 89.3% | 4.37E-227 | 7.94E-223 |
| 2 | FABP1 | Enterocytes | 1.1308551 | 98.9% | 94.1% | 7.60E-117 | 1.38E-112 |
| 2 | SLC40A1 | Enterocytes | 1.1214392 | 85.0% | 67.2% | 9.51E-118 | 1.73E-113 |
| 2 | CYBA | Microfold cells | 1.1114272 | 100.0% | 90.3% | 0.00E+00 | 0.00E+00 |
| 2 | CFB | Paneth cells | 1.1013568 | 73.2% | 54.8% | 1.18E-90 | 2.15E-86 |
| 2 | PMEPA1 | Enterocytes | 1.0935637 | 73.0% | 52.4% | 2.92E-89 | 5.32E-85 |
| 2 | TM4SF20 | Paneth cells | 1.0372421 | 70.9% | 54.5% | 2.59E-55 | 4.72E-51 |
| 2 | FABP5 | Enteroendocrine cells | 1.0247479 | 63.4% | 21.5% | 5.01E-287 | 9.12E-283 |
| 2 | MARCKSL1 | Microfold cells | 0.9992996 | 95.5% | 68.0% | 1.44E-273 | 2.61E-269 |
| 2 | TMEM45B | Enterocytes | 0.9856707 | 90.8% | 63.9% | 1.48E-181 | 2.69E-177 |
| 2 | SOX9 | Tuft cells | 0.9759621 | 74.0% | 44.7% | 1.20E-141 | 2.18E-137 |
| 2 | EHF | Enterocytes | 0.973753 | 62.8% | 28.2% | 7.02E-193 | 1.28E-188 |
| 2 | DMBT1 | Paneth cells | 0.9691498 | 66.9% | 45.8% | 4.14E-62 | 7.53E-58 |
| 2 | BACE2 | Goblet cells | 0.9389672 | 89.4% | 59.9% | 8.16E-202 | 1.48E-197 |
| 2 | CCL20 | Microfold cells | 0.9353321 | 38.3% | 28.5% | 3.55E-16 | 6.45E-12 |
| 2 | AHR | Microfold cells | 0.9275281 | 63.8% | 31.4% | 6.42E-166 | 1.17E-161 |
| 2 | MUC4 | Goblet cells | 0.922067 | 45.3% | 23.5% | 9.21E-91 | 1.67E-86 |
| 2 | FERMT1 | Enterocytes | 0.9071143 | 76.4% | 39.7% | 7.77E-202 | 1.41E-197 |
| 2 | ONECUT2 | Microfold cells | 0.8963527 | 60.0% | 27.1% | 8.44E-170 | 1.54E-165 |
| 2 | CFD | Paneth cells | 0.8929138 | 51.0% | 21.5% | 1.39E-145 | 2.53E-141 |
| 2 | PPP1R14D | Enterocytes | 0.8878413 | 80.3% | 56.5% | 4.24E-141 | 7.72E-137 |
| 2 | FUOM | Enterocytes | 0.879743 | 89.9% | 58.6% | 9.93E-212 | 1.81E-207 |
| 2 | PRSS2 | Enterocytes | 0.879006 | 41.1% | 20.7% | 3.22E-76 | 5.85E-72 |
| 2 | KLF6 | Tuft cells | 0.8721274 | 79.2% | 55.5% | 2.07E-104 | 3.77E-100 |
| 2 | PRR15L | Enterocytes | 0.8714892 | 83.8% | 47.7% | 7.95E-224 | 1.45E-219 |
| 2 | REG4 | Enterochromaffin cells | 0.8135389 | 87.6% | 75.9% | 2.15E-59 | 3.91E-55 |
| 2 | CBR1 | Enterocytes | 0.8027643 | 82.1% | 41.3% | 1.01E-248 | 1.84E-244 |
| 2 | TFF3 | Goblet cells | 0.7971737 | 99.7% | 98.9% | 2.32E-125 | 4.22E-121 |
| 2 | CXCL16 | Microfold cells | 0.7968075 | 81.2% | 49.4% | 2.72E-172 | 4.95E-168 |
| 2 | TMPRSS2 | Microfold cells | 0.7935473 | 77.0% | 56.3% | 9.50E-81 | 1.73E-76 |
| 2 | IRF2 | Microfold cells | 0.7901825 | 56.3% | 25.0% | 6.03E-166 | 1.10E-161 |
| 2 | CDH17 | Enterocytes | 0.7889938 | 82.8% | 55.3% | 2.48E-136 | 4.52E-132 |
| 2 | CTSD | Microfold cells | 0.7758681 | 95.7% | 83.1% | 1.42E-131 | 2.59E-127 |
| 2 | NFIB | Microfold cells | 0.7547055 | 50.1% | 19.6% | 1.29E-167 | 2.35E-163 |
| 2 | FOSL2 | Microfold cells | 0.7276502 | 53.3% | 30.7% | 1.02E-89 | 1.86E-85 |
| 2 | CTNNB1 | Paneth cells | 0.717171 | 70.5% | 43.7% | 9.28E-117 | 1.69E-112 |
| 2 | KCNE3 | Crypt cells | 0.6969017 | 70.2% | 31.5% | 2.71E-205 | 4.92E-201 |
| 2 | MMP7 | Paneth cells | 0.6876848 | 89.9% | 74.9% | 1.82E-101 | 3.30E-97 |
| 2 | CREB3L1 | Goblet cells | 0.6684001 | 73.1% | 40.2% | 6.98E-159 | 1.27E-154 |
| 2 | VEGFA | Enteroendocrine cells | 0.6535365 | 71.0% | 48.5% | 3.62E-80 | 6.58E-76 |
| 2 | PLIN3 | Enterocytes | 0.6335838 | 68.3% | 39.0% | 1.13E-130 | 2.05E-126 |
| 2 | IL13RA1 | Tuft cells | 0.6252154 | 50.8% | 27.9% | 6.59E-92 | 1.20E-87 |
| 2 | PROM1 | Crypt cells | 0.6174054 | 51.5% | 27.2% | 1.35E-93 | 2.45E-89 |
| 2 | PLA2G10 | Goblet cells | 0.6171797 | 75.3% | 45.3% | 7.10E-137 | 1.29E-132 |
| 2 | PDHA1 | Enterocytes | 0.6154997 | 58.3% | 22.9% | 2.28E-198 | 4.15E-194 |
| 2 | MANF | Goblet cells | 0.6046249 | 67.4% | 31.9% | 2.59E-172 | 4.71E-168 |
| 2 | ALCAM | Enteroendocrine cells | 0.5904739 | 42.3% | 19.3% | 1.81E-92 | 3.29E-88 |
| 2 | NFIC | Microfold cells | 0.5820787 | 43.6% | 14.6% | 1.98E-168 | 3.60E-164 |
| 2 | TMC4 | Enterocytes | 0.5748455 | 66.4% | 45.9% | 3.13E-80 | 5.69E-76 |
| 2 | CTSH | Microfold cells | 0.5744379 | 68.2% | 36.6% | 1.34E-143 | 2.43E-139 |
| 2 | GALNT12 | Goblet cells | 0.5709297 | 45.9% | 17.2% | 6.24E-151 | 1.14E-146 |
| 2 | KCNN4 | Paneth cells | 0.5592862 | 53.6% | 23.4% | 4.52E-144 | 8.22E-140 |
| 2 | KCNQ1 | Crypt cells | 0.5495315 | 47.7% | 16.1% | 7.31E-185 | 1.33E-180 |
| 2 | ACSL5 | Enterocytes | 0.5459609 | 68.8% | 38.7% | 7.34E-125 | 1.33E-120 |
| 2 | RAC2 | Microfold cells | 0.507982 | 53.8% | 26.4% | 7.68E-114 | 1.40E-109 |
| 2 | DGAT1 | Enterocytes | 0.4980593 | 66.5% | 38.6% | 5.98E-108 | 1.09E-103 |
| 2 | CDX1 | Enterocytes | 0.4955692 | 46.4% | 21.8% | 4.68E-102 | 8.50E-98 |
| 2 | LGR4 | Paneth cells | 0.4954727 | 49.8% | 25.2% | 1.88E-94 | 3.42E-90 |
| 2 | CCK | Enteroendocrine cells | 0.4851624 | 44.5% | 25.0% | 9.59E-63 | 1.74E-58 |
| 2 | RAP1A | Paneth cells | 0.4795732 | 49.6% | 23.5% | 6.22E-109 | 1.13E-104 |
| 2 | VNN1 | Enterocytes | 0.4763629 | 48.0% | 27.4% | 1.16E-68 | 2.10E-64 |
| 2 | RIPK2 | Paneth cells | 0.4721917 | 35.5% | 13.4% | 1.16E-107 | 2.10E-103 |
| 2 | CDX2 | Enterocytes | 0.4715356 | 48.2% | 23.8% | 9.91E-94 | 1.80E-89 |
| 2 | VIL1 | Enterocytes | 0.4678996 | 72.5% | 44.3% | 3.25E-100 | 5.91E-96 |
| 2 | NUCB2 | Enteroendocrine cells | 0.4674282 | 43.7% | 18.2% | 7.63E-116 | 1.39E-111 |
| 2 | SLC11A2 | Enterocytes | 0.4651626 | 59.1% | 37.2% | 7.58E-66 | 1.38E-61 |
| 2 | ALOX5 | Tuft cells | 0.438358 | 46.7% | 23.0% | 2.01E-91 | 3.66E-87 |
| 2 | IRF6 | Microfold cells | 0.4328625 | 46.5% | 24.3% | 9.99E-79 | 1.82E-74 |
| 2 | CEACAM1 | Enterocytes | 0.4236273 | 61.4% | 45.3% | 4.56E-32 | 8.28E-28 |
| 2 | CASP6 | Enterocytes | 0.4023809 | 43.7% | 18.4% | 2.30E-111 | 4.18E-107 |
| 2 | HNF4A | Enterocytes | 0.4018946 | 47.2% | 24.3% | 1.03E-83 | 1.87E-79 |
| 2 | MAOA | Enterocytes | 0.399945 | 38.6% | 20.3% | 2.68E-60 | 4.88E-56 |
| 2 | GJB2 | Microfold cells | 0.3841629 | 30.5% | 10.2% | 5.69E-108 | 1.04E-103 |
| 2 | ABHD11 | Enterocytes | 0.3674104 | 42.8% | 19.7% | 4.82E-90 | 8.76E-86 |
| 2 | IL4R | Paneth cells | 0.3672267 | 38.0% | 17.9% | 7.63E-74 | 1.39E-69 |
| 2 | TULP4 | Microfold cells | 0.3489819 | 35.7% | 17.9% | 2.76E-60 | 5.01E-56 |
| 2 | FAHD1 | Enterocytes | 0.3461915 | 36.9% | 15.0% | 1.66E-95 | 3.02E-91 |
| 2 | GLS | Enterocytes | 0.3377488 | 38.1% | 21.9% | 3.32E-44 | 6.04E-40 |
| 2 | SLC12A2 | Crypt cells | 0.3205903 | 64.0% | 32.8% | 4.39E-124 | 7.99E-120 |
| 2 | DEGS2 | Enterocytes | 0.3183624 | 36.0% | 14.4% | 2.41E-92 | 4.39E-88 |
| 2 | KLK1 | Goblet cells | 0.3121654 | 27.3% | 8.7% | 4.38E-99 | 7.96E-95 |
| 2 | GJB1 | Enterochromaffin cells | 0.3110341 | 38.6% | 19.3% | 7.97E-66 | 1.45E-61 |
| 2 | GULP1 | Enterocytes | 0.3026288 | 33.0% | 15.1% | 6.00E-65 | 1.09E-60 |
| 2 | RELB | Microfold cells | 0.2994006 | 37.7% | 20.1% | 1.28E-53 | 2.32E-49 |
| 2 | MUC13 | Goblet cells | 0.2985961 | 73.3% | 55.9% | 9.08E-35 | 1.65E-30 |
| 2 | BMI1 | Crypt cells | 0.2896259 | 27.1% | 9.4% | 4.14E-86 | 7.54E-82 |
| 2 | SLC5A1 | Enterocytes | 0.2875894 | 40.6% | 25.7% | 2.80E-32 | 5.09E-28 |
| 2 | NPC1L1 | Enterocytes | 0.2683014 | 39.6% | 21.9% | 6.78E-50 | 1.23E-45 |
| 2 | NR5A2 | Enterocytes | 0.2612851 | 34.7% | 17.3% | 8.69E-55 | 1.58E-50 |
| 4 | TM4SF20 | Paneth cells | 1.3097768 | 75.0% | 54.7% | 2.54E-77 | 4.62E-73 |
| 4 | MUC13 | Goblet cells | 1.2413327 | 86.3% | 55.2% | 1.78E-171 | 3.25E-167 |
| 4 | CEACAM1 | Enterocytes | 1.1959813 | 83.9% | 43.6% | 2.29E-229 | 4.16E-225 |
| 4 | RNF186 | Enterocytes | 1.1642879 | 75.2% | 27.3% | 1.03E-275 | 1.87E-271 |
| 4 | FABP1 | Enterocytes | 1.1520904 | 98.2% | 94.4% | 1.05E-67 | 1.91E-63 |
| 4 | CD55 | Enterocytes | 1.1245983 | 94.2% | 67.5% | 1.20E-179 | 2.18E-175 |
| 4 | SLC40A1 | Enterocytes | 1.0942233 | 95.4% | 66.8% | 1.33E-203 | 2.42E-199 |
| 4 | CCL20 | Microfold cells | 1.048947 | 51.1% | 27.5% | 1.50E-72 | 2.72E-68 |
| 4 | TMPRSS2 | Microfold cells | 1.0075329 | 88.3% | 56.0% | 5.41E-157 | 9.83E-153 |
| 4 | DMBT1 | Paneth cells | 1.0011236 | 70.2% | 46.2% | 2.43E-72 | 4.41E-68 |
| 4 | TMEM45B | Enterocytes | 0.9915064 | 94.8% | 64.5% | 7.18E-207 | 1.31E-202 |
| 4 | CTSD | Microfold cells | 0.9321794 | 99.0% | 83.2% | 1.62E-175 | 2.94E-171 |
| 4 | ELF3 | Enterocytes | 0.9249493 | 98.7% | 81.6% | 3.23E-185 | 5.87E-181 |
| 4 | CDC42EP5 | Enterocytes | 0.8885963 | 99.1% | 85.5% | 7.90E-164 | 1.44E-159 |
| 4 | TFF3 | Goblet cells | 0.8690179 | 99.9% | 98.9% | 3.97E-134 | 7.23E-130 |
| 4 | VEGFA | Enteroendocrine cells | 0.8476351 | 84.7% | 47.9% | 8.20E-176 | 1.49E-171 |
| 4 | SLC11A2 | Enterocytes | 0.8385241 | 71.6% | 36.7% | 1.26E-142 | 2.30E-138 |
| 4 | TNFAIP2 | Microfold cells | 0.8207 | 87.1% | 64.7% | 6.02E-77 | 1.10E-72 |
| 4 | ADM | Enterochromaffin cells | 0.7892354 | 57.9% | 21.6% | 8.11E-167 | 1.48E-162 |
| 4 | CFB | Paneth cells | 0.7567676 | 84.4% | 54.3% | 2.90E-119 | 5.28E-115 |
| 4 | KLF3 | Tuft cells | 0.7552432 | 74.9% | 44.6% | 3.20E-103 | 5.82E-99 |
| 4 | KLF6 | Tuft cells | 0.7534225 | 85.8% | 55.7% | 1.07E-117 | 1.94E-113 |
| 4 | BACE2 | Goblet cells | 0.745421 | 92.1% | 60.7% | 7.62E-161 | 1.39E-156 |
| 4 | FOSL2 | Microfold cells | 0.7192228 | 63.2% | 30.5% | 1.92E-120 | 3.50E-116 |
| 4 | PRAP1 | Enterocytes | 0.7027482 | 88.3% | 68.1% | 2.76E-86 | 5.02E-82 |
| 4 | PPP1R14D | Enterocytes | 0.6238619 | 83.9% | 57.0% | 1.83E-97 | 3.32E-93 |
| 4 | SOX9 | Tuft cells | 0.6126189 | 72.8% | 45.9% | 2.49E-77 | 4.52E-73 |
| 4 | KLF5 | Enterocytes | 0.6067136 | 83.6% | 57.3% | 2.11E-86 | 3.84E-82 |
| 4 | TMC4 | Enterocytes | 0.5956275 | 76.9% | 45.5% | 5.50E-110 | 1.00E-105 |
| 4 | **MUC4** | **Goblet cells** | 0.539216 | 48.0% | 24.1% | 6.84E-65 | 1.24E-60 |
| 4 | PPARG | Enterocytes | 0.5282967 | 74.3% | 48.1% | 1.25E-80 | 2.28E-76 |
| 4 | IRF6 | Microfold cells | 0.5132067 | 52.2% | 24.5% | 3.85E-91 | 7.01E-87 |
| 4 | CFD | Paneth cells | 0.5074274 | 41.0% | 23.6% | 4.71E-40 | 8.56E-36 |
| 4 | CD24 | Paneth cells | 0.5032573 | 98.7% | 89.9% | 7.02E-95 | 1.28E-90 |
| 4 | IRF2 | Microfold cells | 0.496896 | 52.3% | 26.5% | 2.57E-76 | 4.67E-72 |
| 4 | ALCAM | Enteroendocrine cells | 0.4910988 | 39.2% | 20.4% | 9.03E-50 | 1.64E-45 |
| 4 | IL13RA1 | Tuft cells | 0.4701594 | 56.1% | 28.2% | 3.91E-81 | 7.11E-77 |
| 4 | PMEPA1 | Enterocytes | 0.4543878 | 72.5% | 53.2% | 2.34E-40 | 4.26E-36 |
| 4 | OLFM4 | Crypt cells | 0.4528637 | 82.7% | 64.4% | 9.93E-47 | 1.81E-42 |
| 4 | PROM1 | Crypt cells | 0.4503314 | 47.6% | 28.5% | 2.99E-44 | 5.44E-40 |
| 4 | CDH17 | Enterocytes | 0.4445248 | 81.4% | 56.5% | 1.85E-75 | 3.37E-71 |
| 4 | AGR2 | Goblet cells | 0.4427061 | 98.8% | 90.0% | 6.83E-85 | 1.24E-80 |
| 4 | KRT20 | Goblet cells | 0.4308565 | 43.0% | 32.8% | 7.89E-15 | 1.43E-10 |
| 4 | PRR15L | Enterocytes | 0.4302311 | 74.0% | 50.0% | 2.48E-59 | 4.50E-55 |
| 4 | TRNP1 | Enteroendocrine cells | 0.4261066 | 73.6% | 51.7% | 1.27E-50 | 2.31E-46 |
| 4 | IRF7 | Tuft cells | 0.4122107 | 39.0% | 13.9% | 3.58E-104 | 6.51E-100 |
| 4 | LYZ | Paneth cells | 0.409537 | 99.9% | 96.1% | 1.17E-89 | 2.13E-85 |
| 4 | PLIN3 | Enterocytes | 0.4093718 | 65.0% | 40.5% | 5.30E-61 | 9.64E-57 |
| 4 | NUCB2 | Enteroendocrine cells | 0.4001016 | 39.0% | 19.6% | 1.42E-53 | 2.58E-49 |
| 4 | FUOM | Enterocytes | 0.3804752 | 78.8% | 60.9% | 2.96E-41 | 5.38E-37 |
| 4 | VIL1 | Enterocytes | 0.3570593 | 66.1% | 46.0% | 3.07E-42 | 5.58E-38 |
| 4 | RAC2 | Microfold cells | 0.3538066 | 49.8% | 27.8% | 6.16E-52 | 1.12E-47 |
| 4 | CXCL16 | Microfold cells | 0.3502451 | 72.5% | 51.5% | 1.11E-43 | 2.02E-39 |
| 4 | MARCKSL1 | Microfold cells | 0.3304908 | 89.9% | 69.6% | 3.79E-51 | 6.89E-47 |
| 4 | LGR4 | Paneth cells | 0.3240335 | 47.1% | 26.4% | 2.73E-48 | 4.96E-44 |
| 4 | IL4R | Paneth cells | 0.3177515 | 35.8% | 18.8% | 5.36E-42 | 9.74E-38 |
| 4 | RELB | Microfold cells | 0.3176998 | 39.1% | 20.6% | 3.16E-47 | 5.74E-43 |
| 4 | CYBA | Microfold cells | 0.3134769 | 98.9% | 90.8% | 8.45E-44 | 1.54E-39 |
| 4 | ONECUT2 | Microfold cells | 0.302114 | 47.2% | 29.6% | 1.91E-31 | 3.47E-27 |
| 4 | AHR | Microfold cells | 0.29998 | 51.4% | 33.9% | 6.71E-32 | 1.22E-27 |
| 4 | REG4 | Enterochromaffin cells | 0.2958033 | 82.1% | 76.9% | 2.45E-12 | 4.46E-08 |
| 4 | GPX2 | Paneth cells | 0.2931944 | 97.6% | 91.4% | 1.40E-38 | 2.54E-34 |
| 4 | CTNNB1 | Paneth cells | 0.2822729 | 62.6% | 45.5% | 1.87E-31 | 3.40E-27 |
| 4 | MAOA | Enterocytes | 0.2797748 | 36.5% | 21.2% | 1.07E-30 | 1.94E-26 |
| 4 | MANF | Goblet cells | 0.2750719 | 50.9% | 34.9% | 4.55E-27 | 8.27E-23 |
| 4 | SLC15A1 | Enterocytes | 0.2718359 | 33.5% | 17.7% | 3.89E-37 | 7.07E-33 |
| 4 | EHF | Enterocytes | 0.2682849 | 46.5% | 31.2% | 1.67E-25 | 3.04E-21 |
| 4 | PLA2G10 | Goblet cells | 0.26469 | 68.1% | 47.2% | 1.38E-35 | 2.50E-31 |
| 4 | PHGR1 | Goblet cells | 0.2631344 | 98.5% | 95.3% | 3.58E-23 | 6.51E-19 |
| 4 | CREB3L1 | Goblet cells | 0.2631336 | 62.2% | 42.5% | 1.01E-33 | 1.83E-29 |
| 5 | **MMP7** | **Paneth cells** | 2.7289648 | 95.9% | 75.5% | 7.80E-107 | 1.42E-102 |
| 5 | KRT7 | Goblet cells | 2.1298867 | 99.9% | 90.0% | 3.44E-257 | 6.25E-253 |
| 5 | SPINK4 | Paneth cells | 1.9687568 | 29.0% | 14.6% | 1.19E-30 | 2.17E-26 |
| 5 | PMEPA1 | Enterocytes | 1.8662328 | 98.3% | 52.1% | 3.06E-260 | 5.56E-256 |
| 5 | SLC12A2 | Crypt cells | 1.6983929 | 70.9% | 34.5% | 1.14E-116 | 2.07E-112 |
| 5 | ANXA5 | Microfold cells | 1.4382588 | 97.0% | 70.4% | 1.57E-173 | 2.85E-169 |
| 5 | CD24 | Paneth cells | 1.4040072 | 99.7% | 90.1% | 3.96E-182 | 7.20E-178 |
| 5 | CCK | Enteroendocrine cells | 1.2807094 | 62.8% | 25.1% | 2.11E-135 | 3.83E-131 |
| 5 | CTNNB1 | Paneth cells | 1.0944337 | 74.3% | 45.3% | 2.20E-76 | 4.00E-72 |
| 5 | MARCKSL1 | Microfold cells | 1.0271872 | 97.9% | 69.7% | 4.25E-147 | 7.72E-143 |
| 5 | TNFAIP2 | Microfold cells | 1.010284 | 87.8% | 65.3% | 1.44E-77 | 2.62E-73 |
| 5 | FERMT1 | Enterocytes | 1.0095148 | 84.0% | 41.7% | 3.37E-146 | 6.12E-142 |
| 5 | COL18A1 | Enterocytes | 0.9677843 | 58.0% | 12.1% | 3.69E-260 | 6.70E-256 |
| 5 | KCNE3 | Crypt cells | 0.9604159 | 61.8% | 34.7% | 2.05E-67 | 3.73E-63 |
| 5 | CYBA | Microfold cells | 0.9496354 | 99.2% | 91.0% | 5.64E-79 | 1.03E-74 |
| 5 | SOX9 | Tuft cells | 0.9084428 | 78.8% | 46.4% | 2.60E-83 | 4.72E-79 |
| 5 | PRSS2 | Enterocytes | 0.8647325 | 41.2% | 22.1% | 1.63E-33 | 2.96E-29 |
| 5 | TFF3 | Goblet cells | 0.8302375 | 100.0% | 98.9% | 2.17E-84 | 3.95E-80 |
| 5 | LYZ | Paneth cells | 0.7995394 | 99.7% | 96.3% | 5.79E-59 | 1.05E-54 |
| 5 | AHR | Microfold cells | 0.7662859 | 59.7% | 33.9% | 1.54E-51 | 2.80E-47 |
| 5 | GSTM3 | Enterocytes | 0.7417366 | 83.5% | 61.3% | 1.53E-54 | 2.78E-50 |
| 5 | TULP4 | Microfold cells | 0.7068923 | 49.2% | 18.2% | 1.47E-99 | 2.68E-95 |
| 5 | PLA2G10 | Goblet cells | 0.7059944 | 73.4% | 47.5% | 2.86E-56 | 5.21E-52 |
| 5 | KLF6 | Tuft cells | 0.7034208 | 80.2% | 57.0% | 6.92E-42 | 1.26E-37 |
| 5 | CXCL16 | Microfold cells | 0.6828763 | 79.8% | 51.7% | 5.18E-71 | 9.41E-67 |
| 5 | CDH17 | Enterocytes | 0.6528447 | 73.4% | 57.8% | 2.51E-27 | 4.57E-23 |
| 5 | PPP1R14D | Enterocytes | 0.6252819 | 72.4% | 58.6% | 9.33E-27 | 1.70E-22 |
| 5 | EHF | Enterocytes | 0.6164408 | 48.1% | 31.6% | 2.57E-25 | 4.67E-21 |
| 5 | CTSD | Microfold cells | 0.6138042 | 96.7% | 83.9% | 1.76E-40 | 3.20E-36 |
| 5 | CTSH | Microfold cells | 0.5982448 | 60.8% | 39.3% | 1.11E-38 | 2.03E-34 |
| 5 | VNN1 | Enterocytes | 0.5971123 | 43.3% | 29.1% | 2.44E-21 | 4.44E-17 |
| 5 | ACSL5 | Enterocytes | 0.5777376 | 65.8% | 41.0% | 1.15E-51 | 2.09E-47 |
| 5 | PROM1 | Crypt cells | 0.5736268 | 48.4% | 29.0% | 6.30E-33 | 1.15E-28 |
| 5 | GULP1 | Enterocytes | 0.5715908 | 33.8% | 16.3% | 7.17E-37 | 1.30E-32 |
| 5 | BACE2 | Goblet cells | 0.5713216 | 82.0% | 62.4% | 2.82E-40 | 5.12E-36 |
| 5 | GPX2 | Paneth cells | 0.5567288 | 97.3% | 91.6% | 8.31E-46 | 1.51E-41 |
| 5 | SLC40A1 | Enterocytes | 0.5515893 | 83.5% | 68.5% | 5.28E-23 | 9.60E-19 |
| 5 | MFGE8 | Microfold cells | 0.5469483 | 36.8% | 10.2% | 7.97E-105 | 1.45E-100 |
| 5 | CREB3L1 | Goblet cells | 0.5437355 | 69.3% | 42.7% | 1.57E-51 | 2.85E-47 |
| 5 | PLIN3 | Enterocytes | 0.5197411 | 63.2% | 41.4% | 1.95E-40 | 3.54E-36 |
| 5 | CFB | Paneth cells | 0.4932686 | 66.3% | 56.5% | 1.47E-11 | 2.67E-07 |
| 5 | TMPRSS2 | Microfold cells | 0.4854017 | 78.8% | 57.6% | 6.45E-34 | 1.17E-29 |
| 5 | PHGR1 | Goblet cells | 0.4774958 | 95.9% | 95.5% | 1.68E-13 | 3.06E-09 |
| 5 | GLS | Enterocytes | 0.4730555 | 44.0% | 22.6% | 1.45E-41 | 2.63E-37 |
| 5 | IL13RA1 | Tuft cells | 0.4547128 | 49.2% | 29.6% | 9.38E-32 | 1.70E-27 |
| 5 | TRNP1 | Enteroendocrine cells | 0.4481098 | 79.6% | 52.0% | 3.24E-49 | 5.89E-45 |
| 5 | EPHB2 | Crypt cells | 0.4382521 | 31.4% | 9.0% | 3.42E-83 | 6.22E-79 |
| 5 | NFIC | Microfold cells | 0.4106828 | 40.0% | 16.8% | 1.64E-56 | 2.97E-52 |
| 5 | KLF5 | Enterocytes | 0.3918638 | 74.5% | 58.7% | 4.52E-17 | 8.23E-13 |
| 5 | TMC4 | Enterocytes | 0.38373 | 66.8% | 47.2% | 7.01E-28 | 1.27E-23 |
| 5 | ALCAM | Enteroendocrine cells | 0.370132 | 35.6% | 21.3% | 4.36E-21 | 7.94E-17 |
| 5 | ATG16L1 | Paneth cells | 0.3609958 | 33.8% | 13.0% | 1.25E-55 | 2.26E-51 |
| 5 | BMI1 | Crypt cells | 0.3590221 | 27.2% | 10.6% | 3.72E-43 | 6.76E-39 |
| 5 | LGR4 | Paneth cells | 0.3570892 | 42.7% | 27.3% | 1.51E-21 | 2.74E-17 |
| 5 | RAC2 | Microfold cells | 0.3558436 | 48.5% | 28.6% | 8.47E-32 | 1.54E-27 |
| 5 | MAOA | Enterocytes | 0.3423954 | 38.3% | 21.6% | 3.68E-27 | 6.69E-23 |
| 5 | HNF4A | Enterocytes | 0.3140995 | 41.6% | 26.2% | 1.69E-21 | 3.08E-17 |
| 5 | AGR2 | Paneth cells | 0.3085367 | 94.9% | 90.5% | 1.34E-04 | 1.00E+00 |
| 5 | RIPK2 | Paneth cells | 0.3024056 | 32.2% | 15.2% | 2.18E-33 | 3.97E-29 |
| 5 | CBR1 | Enterocytes | 0.2915066 | 61.7% | 45.5% | 4.15E-18 | 7.54E-14 |
| 5 | FLOT2 | Enterocytes | 0.2876335 | 34.8% | 15.0% | 1.90E-44 | 3.45E-40 |
| 5 | ALOX5 | Tuft cells | 0.2817476 | 39.5% | 25.1% | 2.54E-19 | 4.61E-15 |
| 5 | MANF | Goblet cells | 0.2768123 | 48.1% | 35.6% | 6.00E-13 | 1.09E-08 |
| 5 | RAP1A | Paneth cells | 0.2737808 | 40.2% | 25.9% | 2.22E-18 | 4.05E-14 |
| 5 | CASP6 | Enterocytes | 0.2734154 | 33.9% | 20.8% | 3.55E-19 | 6.45E-15 |
| 5 | ELF3 | Enterocytes | 0.270675 | 93.2% | 82.6% | 2.52E-11 | 4.58E-07 |
| 5 | IRF6 | Microfold cells | 0.2623737 | 39.9% | 26.2% | 1.82E-16 | 3.32E-12 |
| 7 | **MUC4** | **Goblet cells** | 1.222235 | 68.7% | 24.2% | 5.79E-138 | 1.05E-133 |
| 7 | ONECUT2 | Microfold cells | 1.0860054 | 71.6% | 29.3% | 4.86E-122 | 8.84E-118 |
| 7 | KLF6 | Tuft cells | 1.0477885 | 90.4% | 56.9% | 6.71E-97 | 1.22E-92 |
| 7 | TNFAIP2 | Microfold cells | 0.9916834 | 89.6% | 65.6% | 9.57E-69 | 1.74E-64 |
| 7 | KLF5 | Enterocytes | 0.8291456 | 89.1% | 58.3% | 1.31E-80 | 2.38E-76 |
| 7 | GLS | Enterocytes | 0.7959174 | 62.3% | 22.1% | 5.69E-114 | 1.04E-109 |
| 7 | FOSL2 | Microfold cells | 0.7602482 | 70.8% | 31.7% | 1.94E-88 | 3.52E-84 |
| 7 | AHR | Microfold cells | 0.7588435 | 70.1% | 33.8% | 1.94E-80 | 3.52E-76 |
| 7 | VEGFA | Enteroendocrine cells | 0.7584788 | 80.4% | 49.9% | 2.80E-66 | 5.09E-62 |
| 7 | ELF3 | Enterocytes | 0.7556588 | 97.9% | 82.5% | 8.78E-84 | 1.60E-79 |
| 7 | TMPRSS2 | Microfold cells | 0.7390017 | 83.4% | 57.7% | 1.65E-53 | 2.99E-49 |
| 7 | KLF3 | Tuft cells | 0.7061643 | 81.5% | 45.7% | 1.13E-80 | 2.06E-76 |
| 7 | CTNNB1 | Paneth cells | 0.6802778 | 79.5% | 45.5% | 1.75E-69 | 3.18E-65 |
| 7 | CD55 | Enterocytes | 0.6333278 | 91.9% | 68.9% | 1.01E-60 | 1.84E-56 |
| 7 | SOX9 | Tuft cells | 0.6139187 | 71.2% | 47.3% | 5.08E-37 | 9.23E-33 |
| 7 | MUC13 | Goblet cells | 0.6087357 | 82.5% | 56.9% | 3.17E-42 | 5.76E-38 |
| 7 | CFB | Paneth cells | 0.6041751 | 80.0% | 56.0% | 3.11E-44 | 5.66E-40 |
| 7 | SLC11A2 | Enterocytes | 0.5866516 | 62.3% | 38.9% | 6.10E-36 | 1.11E-31 |
| 7 | ALCAM | Enteroendocrine cells | 0.5731076 | 49.7% | 20.8% | 3.22E-59 | 5.85E-55 |
| 7 | PROM1 | Crypt cells | 0.5640026 | 57.8% | 28.9% | 2.45E-50 | 4.45E-46 |
| 7 | EHF | Enterocytes | 0.5561717 | 61.6% | 31.2% | 2.77E-52 | 5.04E-48 |
| 7 | IL4R | Paneth cells | 0.510359 | 46.9% | 19.1% | 6.11E-59 | 1.11E-54 |
| 7 | OLFM4 | Crypt cells | 0.4970924 | 87.6% | 65.0% | 3.27E-25 | 5.95E-21 |
| 7 | CEACAM1 | Enterocytes | 0.4901348 | 68.9% | 46.2% | 6.59E-28 | 1.20E-23 |
| 7 | SLC15A1 | Enterocytes | 0.4816837 | 38.2% | 18.2% | 2.43E-34 | 4.42E-30 |
| 7 | LGR4 | Paneth cells | 0.4691746 | 53.9% | 27.0% | 4.64E-46 | 8.43E-42 |
| 7 | VNN1 | Enterocytes | 0.459369 | 55.7% | 28.8% | 3.82E-45 | 6.95E-41 |
| 7 | IL13RA1 | Tuft cells | 0.4233827 | 53.9% | 29.7% | 4.63E-35 | 8.41E-31 |
| 7 | DMBT1 | Enterocytes | 0.4227728 | 58.8% | 47.9% | 4.49E-06 | 8.16E-02 |
| 7 | TULP4 | Microfold cells | 0.4191586 | 43.9% | 19.0% | 4.86E-46 | 8.83E-42 |
| 7 | PMEPA1 | Enterocytes | 0.4182813 | 77.0% | 53.9% | 1.20E-31 | 2.19E-27 |
| 7 | NFIB | Microfold cells | 0.4167072 | 49.5% | 22.2% | 1.11E-47 | 2.03E-43 |
| 7 | SLC40A1 | Enterocytes | 0.3365306 | 84.7% | 68.7% | 3.80E-23 | 6.92E-19 |
| 7 | NR5A2 | Enterocytes | 0.3332789 | 39.4% | 18.6% | 3.81E-34 | 6.92E-30 |
| 7 | HNF4A | Enterocytes | 0.3066401 | 46.3% | 26.2% | 7.56E-25 | 1.38E-20 |
| 7 | SLC12A2 | Crypt cells | 0.3014612 | 50.8% | 36.1% | 5.98E-12 | 1.09E-07 |
| 7 | TMEM45B | Enterocytes | 0.2989397 | 87.0% | 66.3% | 1.86E-25 | 3.38E-21 |
| 7 | CTSD | Microfold cells | 0.2915682 | 95.3% | 84.1% | 1.92E-20 | 3.48E-16 |
| 7 | TMC4 | Enterocytes | 0.2816079 | 68.9% | 47.4% | 1.58E-22 | 2.87E-18 |
| 7 | CDH17 | Enterocytes | 0.2672011 | 72.3% | 58.1% | 3.12E-13 | 5.68E-09 |
| 7 | PLA2G2A | Paneth cells | 0.2633057 | 33.0% | 14.3% | 2.60E-30 | 4.72E-26 |
| 7 | GALNT12 | Goblet cells | 0.2511636 | 38.2% | 20.0% | 9.97E-24 | 1.81E-19 |
| 8 | PHGR1 | Enterocytes | 2.5436771 | 99.6% | 95.4% | 2.01E-164 | 3.65E-160 |
| 8 | KRT20 | Enterocytes | 2.408736 | 84.2% | 31.5% | 3.15E-210 | 5.72E-206 |
| 8 | FABP2 | Enterocytes | 2.3939266 | 84.8% | 28.5% | 1.73E-227 | 3.15E-223 |
| 8 | REG4 | Enterochromaffin cells | 2.2839268 | 93.8% | 76.7% | 4.46E-109 | 8.11E-105 |
| 8 | ALDOB | Enterocytes | 2.2734611 | 90.5% | 37.9% | 1.34E-213 | 2.44E-209 |
| 8 | RBP2 | Enterocytes | 2.2042808 | 55.6% | 18.5% | 5.98E-107 | 1.09E-102 |
| 8 | FABP1 | Enterocytes | 1.7882132 | 99.8% | 94.5% | 2.73E-131 | 4.97E-127 |
| 8 | PRAP1 | Enterocytes | 1.7305408 | 95.1% | 68.8% | 9.08E-123 | 1.65E-118 |
| 8 | GSTA1 | Enterocytes | 1.6873732 | 59.9% | 12.8% | 2.08E-203 | 3.78E-199 |
| 8 | ANPEP | Enterocytes | 1.644009 | 38.3% | 12.1% | 1.76E-70 | 3.20E-66 |
| 8 | **SI** | **Enterocytes** | 1.6205421 | 60.5% | 12.9% | 3.78E-205 | 6.87E-201 |
| 8 | AGR2 | Goblet cells | 1.5945104 | 98.4% | 90.5% | 2.28E-111 | 4.15E-107 |
| 8 | MTTP | Enterocytes | 1.5785546 | 68.7% | 15.2% | 4.46E-240 | 8.10E-236 |
| 8 | TM4SF5 | Enterocytes | 1.5236956 | 81.7% | 29.2% | 5.53E-187 | 1.01E-182 |
| 8 | CBR1 | Enterocytes | 1.4168701 | 90.9% | 44.5% | 5.55E-153 | 1.01E-148 |
| 8 | OLFM4 | Crypt cells | 1.3659362 | 79.2% | 65.5% | 2.28E-25 | 4.15E-21 |
| 8 | TM4SF20 | Paneth cells | 1.3612555 | 83.7% | 55.3% | 1.15E-57 | 2.08E-53 |
| 8 | LYZ | Paneth cells | 1.3397759 | 100.0% | 96.3% | 4.40E-94 | 8.01E-90 |
| 8 | SLC12A2 | Crypt cells | 1.1517261 | 56.0% | 35.9% | 8.67E-32 | 1.58E-27 |
| 8 | DGAT1 | Enterocytes | 1.1489495 | 83.1% | 40.3% | 1.01E-126 | 1.83E-122 |
| 8 | DMBT1 | Enterocytes | 1.130379 | 49.8% | 48.4% | 1.31E-05 | 2.38E-01 |
| 8 | MUC13 | Enterocytes | 1.1214093 | 91.2% | 56.6% | 3.25E-93 | 5.90E-89 |
| 8 | SLC5A1 | Enterocytes | 1.1117776 | 76.1% | 25.4% | 1.44E-160 | 2.63E-156 |
| 8 | PLA2G10 | Goblet cells | 1.1088908 | 84.2% | 47.6% | 5.12E-103 | 9.31E-99 |
| 8 | CDH17 | Enterocytes | 1.1047202 | 91.4% | 57.3% | 2.40E-101 | 4.36E-97 |
| 8 | **VIL1** | **Enterocytes** | 1.0957501 | 85.6% | 46.1% | 3.64E-112 | 6.61E-108 |
| 8 | FUOM | Enterocytes | 1.0328843 | 92.8% | 61.2% | 9.54E-108 | 1.74E-103 |
| 8 | CDC42EP5 | Enterocytes | 1.0314011 | 100.0% | 86.1% | 1.89E-109 | 3.44E-105 |
| 8 | CYBA | Microfold cells | 0.9629255 | 99.4% | 91.2% | 3.66E-86 | 6.65E-82 |
| 8 | MEP1A | Enterocytes | 0.9537389 | 49.8% | 8.6% | 1.82E-195 | 3.31E-191 |
| 8 | TFF3 | Goblet cells | 0.9279267 | 99.4% | 99.0% | 3.38E-31 | 6.14E-27 |
| 8 | CD24 | Paneth cells | 0.8916264 | 94.7% | 90.5% | 2.50E-45 | 4.54E-41 |
| 8 | TM6SF2 | Enterocytes | 0.8700119 | 59.5% | 12.8% | 1.92E-192 | 3.50E-188 |
| 8 | CLDN15 | Enterocytes | 0.8453627 | 55.1% | 21.0% | 2.05E-84 | 3.73E-80 |
| 8 | PPP1R14D | Enterocytes | 0.844764 | 93.0% | 58.0% | 4.63E-97 | 8.41E-93 |
| 8 | SLC40A1 | Enterocytes | 0.840645 | 89.3% | 68.5% | 2.16E-49 | 3.92E-45 |
| 8 | ACSL5 | Enterocytes | 0.8066853 | 76.7% | 41.0% | 2.11E-77 | 3.83E-73 |
| 8 | SLC39A5 | Enterocytes | 0.8031998 | 61.7% | 16.4% | 2.75E-153 | 5.01E-149 |
| 8 | PRR15L | Enterocytes | 0.8002476 | 84.2% | 50.8% | 1.89E-76 | 3.44E-72 |
| 8 | KCNE3 | Crypt cells | 0.767786 | 61.5% | 35.3% | 9.02E-43 | 1.64E-38 |
| 8 | CTSD | Microfold cells | 0.7491277 | 98.6% | 84.0% | 1.63E-71 | 2.96E-67 |
| 8 | KHK | Enterocytes | 0.7482688 | 30.7% | 8.3% | 4.60E-66 | 8.37E-62 |
| 8 | CDX1 | Enterocytes | 0.7167067 | 64.0% | 23.1% | 1.74E-101 | 3.16E-97 |
| 8 | CTSH | Microfold cells | 0.7125573 | 73.9% | 39.1% | 2.65E-69 | 4.83E-65 |
| 8 | SLC51B | Enterocytes | 0.7059789 | 31.1% | 3.0% | 9.00E-201 | 1.64E-196 |
| 8 | CDX2 | Enterocytes | 0.6904999 | 56.8% | 25.5% | 5.23E-64 | 9.51E-60 |
| 8 | MYO1A | Enterocytes | 0.6709348 | 43.0% | 8.7% | 3.78E-138 | 6.87E-134 |
| 8 | ISX | Enterocytes | 0.6373578 | 42.4% | 7.8% | 2.40E-151 | 4.37E-147 |
| 8 | OTC | Enterocytes | 0.6148372 | 29.6% | 2.3% | 1.33E-231 | 2.42E-227 |
| 8 | TMEM45B | Enterocytes | 0.6103095 | 90.9% | 66.2% | 1.84E-50 | 3.35E-46 |
| 8 | EPHX2 | Enterocytes | 0.603544 | 42.2% | 9.1% | 1.23E-124 | 2.23E-120 |
| 8 | MARCKSL1 | Microfold cells | 0.5911924 | 87.7% | 70.7% | 5.90E-36 | 1.07E-31 |
| 8 | CYBRD1 | Enterocytes | 0.5892586 | 26.1% | 6.1% | 1.63E-67 | 2.96E-63 |
| 8 | MOGAT2 | Enterocytes | 0.5802543 | 38.5% | 9.8% | 1.41E-92 | 2.57E-88 |
| 8 | CTNNB1 | Paneth cells | 0.5772495 | 71.2% | 46.0% | 4.57E-31 | 8.31E-27 |
| 8 | CREB3L1 | Goblet cells | 0.5753941 | 67.9% | 43.3% | 2.59E-38 | 4.72E-34 |
| 8 | CPS1 | Enterocytes | 0.5744862 | 30.5% | 3.2% | 2.31E-185 | 4.21E-181 |
| 8 | KLF6 | Tuft cells | 0.5713265 | 84.2% | 57.3% | 1.07E-41 | 1.94E-37 |
| 8 | TRNP1 | Enteroendocrine cells | 0.5705205 | 72.2% | 52.9% | 4.82E-29 | 8.77E-25 |
| 8 | RNF186 | Enterocytes | 0.5560172 | 48.8% | 31.0% | 9.13E-21 | 1.66E-16 |
| 8 | VNN1 | Enterocytes | 0.5559764 | 50.2% | 29.1% | 1.97E-28 | 3.58E-24 |
| 8 | CDHR2 | Tuft cells | 0.5555172 | 40.3% | 12.0% | 1.06E-75 | 1.94E-71 |
| 8 | RAP1A | Paneth cells | 0.5443644 | 58.2% | 25.4% | 5.37E-64 | 9.77E-60 |
| 8 | GPX2 | Paneth cells | 0.5387191 | 90.5% | 92.0% | 3.99E-18 | 7.25E-14 |
| 8 | GJB1 | Enterochromaffin cells | 0.5374033 | 51.4% | 20.4% | 6.54E-67 | 1.19E-62 |
| 8 | GSTM3 | Enterocytes | 0.5280689 | 83.1% | 61.8% | 5.42E-34 | 9.85E-30 |
| 8 | SLC11A2 | Enterocytes | 0.5264476 | 57.2% | 39.2% | 3.61E-19 | 6.56E-15 |
| 8 | SLC51A | Enterocytes | 0.5221982 | 29.4% | 9.2% | 4.08E-51 | 7.42E-47 |
| 8 | CXCL16 | Microfold cells | 0.5175707 | 82.7% | 52.1% | 3.81E-48 | 6.92E-44 |
| 8 | PDHA1 | Enterocytes | 0.5147105 | 57.2% | 26.1% | 4.17E-56 | 7.58E-52 |
| 8 | NPC1L1 | Enterocytes | 0.487783 | 51.4% | 22.9% | 4.60E-52 | 8.37E-48 |
| 8 | APOA1 | Enterocytes | 0.4819974 | 25.7% | 17.7% | 6.85E-07 | 1.25E-02 |
| 8 | HNF4A | Enterocytes | 0.4752325 | 50.4% | 26.1% | 4.86E-36 | 8.83E-32 |
| 8 | ABCG2 | Enterocytes | 0.4446536 | 25.7% | 3.2% | 7.59E-131 | 1.38E-126 |
| 8 | GLS | Enterocytes | 0.4374502 | 44.7% | 23.0% | 8.20E-29 | 1.49E-24 |
| 8 | KRTCAP3 | Enterocytes | 0.4352378 | 81.9% | 55.7% | 5.61E-35 | 1.02E-30 |
| 8 | MANF | Goblet cells | 0.4176367 | 57.4% | 35.4% | 5.76E-27 | 1.05E-22 |
| 8 | DEGS2 | Enterocytes | 0.4067052 | 38.5% | 16.2% | 9.02E-40 | 1.64E-35 |
| 8 | ABHD11 | Enterocytes | 0.4062454 | 42.8% | 21.7% | 1.53E-30 | 2.79E-26 |
| 8 | ELF3 | Enterocytes | 0.4009566 | 95.3% | 82.7% | 3.03E-28 | 5.52E-24 |
| 8 | KLF5 | Enterocytes | 0.396615 | 82.7% | 58.7% | 8.59E-27 | 1.56E-22 |
| 8 | APOBEC1 | Enterocytes | 0.3955206 | 27.6% | 8.1% | 1.94E-51 | 3.53E-47 |
| 8 | LGR4 | Paneth cells | 0.3852105 | 50.0% | 27.3% | 5.48E-29 | 9.96E-25 |
| 8 | CCL20 | Microfold cells | 0.3823222 | 34.6% | 29.5% | 2.17E-03 | 1.00E+00 |
| 8 | FAHD1 | Enterocytes | 0.3821849 | 37.7% | 16.9% | 7.90E-35 | 1.44E-30 |
| 8 | CASP6 | Enterocytes | 0.3817683 | 43.0% | 20.6% | 6.53E-34 | 1.19E-29 |
| 8 | SDSL | Enterocytes | 0.3814496 | 29.8% | 8.3% | 2.29E-60 | 4.16E-56 |
| 8 | HMOX1 | Enterocytes | 0.3680304 | 31.5% | 14.0% | 3.00E-28 | 5.46E-24 |
| 8 | TMC4 | Enterocytes | 0.3678626 | 70.4% | 47.5% | 1.27E-26 | 2.30E-22 |
| 8 | FABP5 | Enteroendocrine cells | 0.3547392 | 41.6% | 26.1% | 6.67E-16 | 1.21E-11 |
| 8 | CD55 | Enterocytes | 0.3147977 | 84.0% | 69.4% | 8.18E-16 | 1.49E-11 |
| 8 | NUCB2 | Enteroendocrine cells | 0.312254 | 36.8% | 20.7% | 1.10E-18 | 2.00E-14 |
| 8 | PLIN3 | Enterocytes | 0.3040705 | 56.0% | 42.2% | 7.93E-14 | 1.44E-09 |
| 8 | CFB | Paneth cells | 0.301982 | 67.9% | 56.6% | 3.42E-09 | 6.22E-05 |
| 8 | RAC2 | Microfold cells | 0.3015192 | 47.1% | 29.1% | 3.39E-18 | 6.17E-14 |
| 8 | NFIB | Microfold cells | 0.2995167 | 39.7% | 22.8% | 5.58E-18 | 1.01E-13 |
| 8 | NR5A2 | Enterocytes | 0.2985843 | 37.4% | 18.7% | 1.74E-25 | 3.16E-21 |
| 8 | AHR | Microfold cells | 0.2923789 | 53.3% | 34.7% | 2.70E-17 | 4.91E-13 |
| 8 | NR1H4 | Enterocytes | 0.2823862 | 25.5% | 8.0% | 4.18E-41 | 7.60E-37 |
| 8 | ESPN | Tuft cells | 0.272884 | 26.5% | 10.5% | 1.14E-28 | 2.07E-24 |
| 8 | SOX9 | Tuft cells | 0.2687947 | 64.2% | 47.7% | 3.04E-12 | 5.53E-08 |
| 8 | DPP4 | Enterocytes | 0.2580845 | 25.1% | 10.7% | 2.35E-23 | 4.27E-19 |
| 9 | PLA2G2A | Paneth cells | 0.2597873 | 29.5% | 14.6% | 2.71E-14 | 4.92E-10 |
| 10 | MKI67 | Proliferating cells | 2.0368014 | 85.3% | 3.2% | 0.00E+00 | 0.00E+00 |
| 10 | FABP5 | Enteroendocrine cells | 1.5901609 | 80.7% | 25.1% | 3.68E-162 | 6.68E-158 |
| 10 | GPX2 | Paneth cells | 1.0215553 | 99.4% | 91.7% | 3.47E-63 | 6.30E-59 |
| 10 | KRTCAP3 | Enterocytes | 0.9920984 | 91.5% | 55.7% | 2.34E-76 | 4.25E-72 |
| 10 | AGR2 | Goblet cells | 0.926293 | 99.4% | 90.5% | 2.03E-55 | 3.69E-51 |
| 10 | GSTM3 | Enterocytes | 0.7703505 | 92.6% | 61.8% | 3.09E-55 | 5.63E-51 |
| 10 | REG4 | Enterochromaffin cells | 0.7478527 | 92.1% | 76.9% | 1.22E-37 | 2.22E-33 |
| 10 | CYBA | Microfold cells | 0.7277815 | 100.0% | 91.3% | 1.73E-57 | 3.15E-53 |
| 10 | KCNE3 | Crypt cells | 0.5878441 | 70.8% | 35.3% | 1.77E-48 | 3.23E-44 |
| 10 | PDHA1 | Enterocytes | 0.5765646 | 61.5% | 26.3% | 3.29E-53 | 5.99E-49 |
| 10 | PRSS1 | Paneth cells | 0.5340767 | 36.0% | 18.6% | 1.02E-16 | 1.85E-12 |
| 10 | LYZ | Paneth cells | 0.524251 | 100.0% | 96.4% | 1.58E-28 | 2.87E-24 |
| 10 | PRSS2 | Enterocytes | 0.5225928 | 35.7% | 22.9% | 2.35E-08 | 4.27E-04 |
| 10 | ANXA5 | Microfold cells | 0.5213797 | 95.5% | 71.3% | 1.25E-45 | 2.28E-41 |
| 10 | MANF | Goblet cells | 0.5043534 | 65.7% | 35.4% | 1.91E-35 | 3.47E-31 |
| 10 | CBR1 | Enterocytes | 0.499504 | 75.1% | 45.6% | 6.65E-33 | 1.21E-28 |
| 10 | MARCKSL1 | Microfold cells | 0.4706219 | 94.3% | 70.7% | 5.25E-32 | 9.54E-28 |
| 10 | VIL1 | Enterocytes | 0.4549234 | 74.5% | 47.0% | 1.04E-27 | 1.89E-23 |
| 10 | CDX2 | Goblet cells | 0.4330837 | 50.7% | 26.1% | 3.18E-27 | 5.78E-23 |
| 10 | FERMT1 | Enterocytes | 0.4306654 | 70.5% | 43.5% | 1.43E-26 | 2.59E-22 |
| 10 | FUOM | Enterocytes | 0.4080464 | 85.0% | 61.9% | 1.35E-21 | 2.45E-17 |
| 10 | CASP6 | Enterocytes | 0.40656 | 44.2% | 20.9% | 2.24E-28 | 4.08E-24 |
| 10 | CCK | Enteroendocrine cells | 0.3939813 | 44.2% | 26.9% | 1.80E-13 | 3.28E-09 |
| 10 | PRR15L | Enterocytes | 0.3938123 | 71.7% | 51.6% | 1.59E-16 | 2.89E-12 |
| 10 | CTSH | Microfold cells | 0.3823203 | 62.6% | 39.9% | 9.94E-21 | 1.81E-16 |
| 10 | CDC42EP5 | Enterocytes | 0.3561452 | 98.3% | 86.4% | 3.47E-15 | 6.31E-11 |
| 10 | ABHD11 | Enterocytes | 0.3214443 | 41.6% | 22.0% | 2.49E-19 | 4.52E-15 |
| 10 | KLK1 | Goblet cells | 0.307423 | 28.0% | 10.5% | 2.16E-25 | 3.93E-21 |
| 10 | RAP1A | Paneth cells | 0.2977624 | 46.5% | 26.2% | 6.15E-18 | 1.12E-13 |
| 10 | SLC12A2 | Crypt cells | 0.2959854 | 60.6% | 36.0% | 3.06E-19 | 5.56E-15 |
| 10 | FABP1 | Enterocytes | 0.2775483 | 96.3% | 94.7% | 2.88E-11 | 5.25E-07 |
| 10 | ACSL5 | Enterocytes | 0.2716839 | 66.0% | 41.7% | 2.07E-18 | 3.76E-14 |
| 10 | GALNT12 | Goblet cells | 0.2699027 | 36.5% | 20.3% | 1.62E-13 | 2.94E-09 |
| 10 | GATA4 | Enterocytes | 0.2534376 | 31.4% | 13.6% | 1.44E-21 | 2.62E-17 |
| 11 | APOC3 | Enterocytes | 5.7566624 | 77.8% | 16.4% | 5.01E-226 | 9.11E-222 |
| 11 | APOA1 | Enterocytes | 5.6963616 | 81.4% | 16.3% | 3.81E-251 | 6.92E-247 |
| 11 | APOA4 | Enterocytes | 5.2452214 | 71.2% | 9.8% | 2.35E-288 | 4.27E-284 |
| 11 | RBP2 | Enterocytes | 4.7208414 | 89.5% | 18.1% | 3.16E-283 | 5.75E-279 |
| 11 | ANPEP | Enterocytes | 3.5682527 | 88.9% | 11.1% | 0.00E+00 | 0.00E+00 |
| 11 | ALDOB | Enterocytes | 3.1952812 | 98.4% | 38.5% | 6.06E-209 | 1.10E-204 |
| 11 | FABP2 | Enterocytes | 2.8472746 | 95.1% | 29.1% | 7.56E-213 | 1.38E-208 |
| 11 | APOB | Enterocytes | 2.6686397 | 70.6% | 4.4% | 0.00E+00 | 0.00E+00 |
| 11 | PRAP1 | Enterocytes | 2.5659005 | 99.0% | 69.1% | 2.42E-147 | 4.41E-143 |
| 11 | **SI** | **Enterocytes** | 2.524305 | 82.4% | 13.1% | 7.05E-288 | 1.28E-283 |
| 11 | GSTA1 | Enterocytes | 2.2120824 | 62.4% | 13.5% | 8.76E-146 | 1.59E-141 |
| 11 | CYP3A4 | Enterocytes | 2.185416 | 64.7% | 5.4% | 0.00E+00 | 0.00E+00 |
| 11 | PHGR1 | Enterocytes | 2.1045386 | 100.0% | 95.4% | 7.09E-123 | 1.29E-118 |
| 11 | TM4SF5 | Enterocytes | 2.011504 | 89.5% | 29.8% | 1.71E-178 | 3.11E-174 |
| 11 | MTTP | Enterocytes | 1.7889031 | 76.8% | 15.8% | 4.18E-193 | 7.60E-189 |
| 11 | MEP1A | Enterocytes | 1.6463815 | 81.7% | 8.3% | 0.00E+00 | 0.00E+00 |
| 11 | KHK | Enterocytes | 1.6096191 | 65.0% | 7.7% | 2.16E-280 | 3.94E-276 |
| 11 | SLC5A1 | Enterocytes | 1.6028598 | 85.6% | 26.0% | 5.82E-158 | 1.06E-153 |
| 11 | DGAT1 | Enterocytes | 1.2809376 | 88.9% | 40.8% | 3.66E-103 | 6.66E-99 |
| 11 | CBR1 | Enterocytes | 1.2646486 | 79.1% | 45.6% | 2.49E-58 | 4.52E-54 |
| 11 | KRT20 | Enterocytes | 1.2379269 | 87.6% | 32.3% | 7.83E-106 | 1.42E-101 |
| 11 | SLC2A5 | Enterocytes | 1.175924 | 43.1% | 1.3% | 0.00E+00 | 0.00E+00 |
| 11 | ZG16 | Goblet cells | 1.0747598 | 52.0% | 1.9% | 0.00E+00 | 0.00E+00 |
| 11 | MUC13 | Enterocytes | 1.0669953 | 90.5% | 57.2% | 1.71E-58 | 3.11E-54 |
| 11 | MYO1A | Enterocytes | 1.0639568 | 62.7% | 8.7% | 1.86E-221 | 3.38E-217 |
| 11 | ACSL5 | Enterocytes | 0.9656912 | 77.1% | 41.5% | 2.24E-53 | 4.07E-49 |
| 11 | CDHR2 | Tuft cells | 0.9474572 | 59.8% | 12.0% | 2.39E-142 | 4.34E-138 |
| 11 | CREB3L3 | Enterocytes | 0.9159887 | 44.4% | 2.1% | 0.00E+00 | 0.00E+00 |
| 11 | SLC51B | Enterocytes | 0.8957053 | 50.7% | 3.0% | 0.00E+00 | 0.00E+00 |
| 11 | SLC39A5 | Enterocytes | 0.8591075 | 62.1% | 17.2% | 6.35E-101 | 1.15E-96 |
| 11 | ALPI | Enterocytes | 0.8311837 | 41.8% | 1.5% | 0.00E+00 | 0.00E+00 |
| 11 | TM4SF20 | Paneth cells | 0.8196883 | 87.9% | 55.7% | 2.46E-35 | 4.47E-31 |
| 11 | CYBRD1 | Enterocytes | 0.8112268 | 41.2% | 6.0% | 2.97E-128 | 5.40E-124 |
| 11 | DPP4 | Enterocytes | 0.8067993 | 48.4% | 10.3% | 2.11E-103 | 3.83E-99 |
| 11 | ABCG2 | Enterocytes | 0.788737 | 41.5% | 3.2% | 3.09E-243 | 5.63E-239 |
| 11 | FABP1 | Enterocytes | 0.7830006 | 100.0% | 94.6% | 1.03E-40 | 1.88E-36 |
| 11 | RNF186 | Enterocytes | 0.7808505 | 69.3% | 30.7% | 2.66E-51 | 4.83E-47 |
| 11 | MOGAT2 | Enterocytes | 0.7700609 | 54.2% | 9.8% | 4.66E-140 | 8.48E-136 |
| 11 | TM6SF2 | Enterocytes | 0.752264 | 57.5% | 13.6% | 1.17E-107 | 2.14E-103 |
| 11 | SLC15A1 | Enterocytes | 0.7443247 | 57.8% | 18.1% | 5.17E-75 | 9.40E-71 |
| 11 | **VIL1** | **Enterocytes** | 0.7238992 | 78.1% | 47.0% | 5.53E-37 | 1.01E-32 |
| 11 | APOBEC1 | Enterocytes | 0.7056592 | 45.1% | 7.9% | 3.11E-116 | 5.65E-112 |
| 11 | AQP11 | Enterocytes | 0.6941777 | 39.2% | 5.1% | 1.46E-140 | 2.65E-136 |
| 11 | SLC51A | Enterocytes | 0.6177331 | 44.4% | 9.1% | 7.39E-94 | 1.34E-89 |
| 11 | TREH | Enterocytes | 0.6009434 | 35.6% | 2.3% | 1.39E-234 | 2.53E-230 |
| 11 | EPHX2 | Enterocytes | 0.5904811 | 43.1% | 9.7% | 1.39E-82 | 2.52E-78 |
| 11 | NPC1L1 | Enterocytes | 0.5716448 | 52.0% | 23.4% | 3.30E-35 | 6.01E-31 |
| 11 | CLDN15 | Enterocytes | 0.5567555 | 51.6% | 21.7% | 9.87E-38 | 1.79E-33 |
| 11 | SLC28A1 | Enterocytes | 0.5539743 | 32.0% | 0.7% | 0.00E+00 | 0.00E+00 |
| 11 | CTSD | Microfold cells | 0.522439 | 98.7% | 84.3% | 1.89E-29 | 3.44E-25 |
| 11 | ABCC2 | Enterocytes | 0.5141934 | 28.4% | 2.5% | 9.94E-144 | 1.81E-139 |
| 11 | PLIN2 | Enterocytes | 0.5070213 | 35.3% | 10.2% | 1.74E-46 | 3.16E-42 |
| 11 | FUOM | Enterocytes | 0.4927881 | 86.9% | 61.9% | 1.62E-23 | 2.95E-19 |
| 11 | PDZK1 | Enterocytes | 0.4899581 | 33.3% | 3.4% | 2.20E-148 | 4.00E-144 |
| 11 | ESPN | Tuft cells | 0.453329 | 39.9% | 10.4% | 2.16E-59 | 3.92E-55 |
| 11 | AQP7 | Enterocytes | 0.4405109 | 29.1% | 1.7% | 1.16E-204 | 2.12E-200 |
| 11 | DMBT1 | Enterocytes | 0.4081332 | 33.3% | 48.9% | 7.53E-03 | 1.00E+00 |
| 11 | HMOX1 | Enterocytes | 0.3617841 | 37.6% | 14.1% | 7.39E-31 | 1.34E-26 |
| 11 | SLC11A2 | Enterocytes | 0.3440392 | 54.2% | 39.6% | 1.98E-08 | 3.60E-04 |
| 11 | ISX | Enterocytes | 0.3406397 | 30.1% | 8.7% | 2.43E-37 | 4.42E-33 |
| 11 | MAOA | Enterocytes | 0.2961257 | 36.6% | 22.3% | 6.20E-10 | 1.13E-05 |
| 12 | APOC3 | Enterocytes | 2.6740208 | 77.6% | 16.6% | 2.10E-183 | 3.81E-179 |
| 12 | APOA1 | Enterocytes | 2.6713295 | 77.2% | 16.6% | 4.45E-181 | 8.08E-177 |
| 12 | RBP2 | Enterocytes | 2.239862 | 90.7% | 18.3% | 1.38E-230 | 2.51E-226 |
| 12 | APOA4 | Enterocytes | 2.1560274 | 63.8% | 10.2% | 4.83E-182 | 8.79E-178 |
| 12 | ANPEP | Enterocytes | 1.642971 | 80.2% | 11.6% | 5.15E-248 | 9.36E-244 |
| 12 | PRAP1 | Enterocytes | 1.0070679 | 92.5% | 69.4% | 5.09E-46 | 9.25E-42 |
| 12 | ALDOB | Enterocytes | 0.8902022 | 87.3% | 39.0% | 8.93E-76 | 1.62E-71 |
| 12 | CYP3A4 | Enterocytes | 0.6723362 | 48.9% | 6.0% | 1.20E-160 | 2.18E-156 |
| 12 | TM4SF5 | Enterocytes | 0.583481 | 68.7% | 30.5% | 7.95E-44 | 1.45E-39 |
| 12 | GSTA1 | Enterocytes | 0.5817792 | 44.0% | 14.1% | 8.83E-44 | 1.61E-39 |
| 12 | APOB | Enterocytes | 0.5356857 | 44.8% | 5.3% | 3.59E-151 | 6.53E-147 |
| 12 | FABP2 | Enterocytes | 0.5107734 | 73.1% | 29.9% | 1.53E-53 | 2.79E-49 |
| 12 | PHGR1 | Enterocytes | 0.4890928 | 100.0% | 95.4% | 2.52E-40 | 4.59E-36 |
| 12 | KHK | Enterocytes | 0.4697529 | 42.5% | 8.5% | 4.16E-79 | 7.57E-75 |
| 12 | **SI** | **Enterocytes** | 0.4208177 | 44.4% | 14.3% | 1.06E-41 | 1.93E-37 |
| 12 | CDHR2 | Tuft cells | 0.2933289 | 29.5% | 12.9% | 6.59E-16 | 1.20E-11 |
| 12 | MYO1A | Enterocytes | 0.2719944 | 28.0% | 9.7% | 1.72E-22 | 3.13E-18 |
| 12 | MEP1A | Enterocytes | 0.2556866 | 30.6% | 9.8% | 6.08E-27 | 1.11E-22 |
| 14 | PMEPA1 | Enterocytes | 1.1535249 | 97.4% | 54.1% | 9.23E-66 | 1.68E-61 |
| 14 | KRT7 | Goblet cells | 1.0773666 | 100.0% | 90.4% | 2.28E-57 | 4.15E-53 |
| 14 | TNFAIP2 | Microfold cells | 0.5895659 | 86.4% | 66.3% | 5.70E-18 | 1.04E-13 |
| 14 | MFGE8 | Microfold cells | 0.4682454 | 37.3% | 11.3% | 2.16E-34 | 3.92E-30 |
| 14 | COL18A1 | Enterocytes | 0.4197152 | 39.0% | 14.5% | 6.55E-25 | 1.19E-20 |
| 14 | KLF6 | Tuft cells | 0.3335969 | 77.2% | 58.1% | 3.47E-07 | 6.32E-03 |
| 14 | CTSD | Microfold cells | 0.3169752 | 96.1% | 84.4% | 1.90E-08 | 3.46E-04 |
| 14 | MAOA | Enterocytes | 0.302542 | 42.1% | 22.2% | 6.51E-13 | 1.18E-08 |
| 14 | ALCAM | Enteroendocrine cells | 0.2861744 | 31.1% | 22.0% | 2.91E-04 | 1.00E+00 |
| 14 | PROM1 | Crypt cells | 0.2748418 | 42.1% | 30.0% | 2.12E-05 | 3.86E-01 |
| 14 | FERMT1 | Enterocytes | 0.274835 | 65.8% | 43.9% | 1.05E-09 | 1.91E-05 |
| 14 | CFB | Paneth cells | 0.2581828 | 68.4% | 56.9% | 3.62E-04 | 1.00E+00 |
| 14 | MUC4 | Goblet cells | 0.2514177 | 38.6% | 26.0% | 6.55E-05 | 1.00E+00 |
| 15 | CCL20 | Microfold cells | 2.3042834 | 63.6% | 29.2% | 2.26E-31 | 4.12E-27 |
| 15 | MUC13 | Goblet cells | 1.7396875 | 76.9% | 57.8% | 1.30E-18 | 2.37E-14 |
| 15 | TM4SF20 | Paneth cells | 1.7173912 | 65.9% | 56.4% | 3.56E-10 | 6.48E-06 |
| 15 | TMPRSS2 | Microfold cells | 1.3799495 | 89.6% | 58.5% | 2.67E-43 | 4.85E-39 |
| 15 | CXCL16 | Microfold cells | 1.3496596 | 94.8% | 52.8% | 1.73E-61 | 3.15E-57 |
| 15 | KRT20 | Goblet cells | 1.3120793 | 54.9% | 33.5% | 4.07E-14 | 7.41E-10 |
| 15 | CD55 | Enterocytes | 1.3075549 | 87.9% | 69.7% | 1.63E-20 | 2.97E-16 |
| 15 | MARCKSL1 | Microfold cells | 1.258107 | 98.3% | 71.0% | 5.22E-45 | 9.49E-41 |
| 15 | MMP7 | Paneth cells | 1.2562385 | 90.8% | 76.6% | 5.26E-19 | 9.56E-15 |
| 15 | TNFAIP2 | Microfold cells | 1.2180817 | 91.3% | 66.4% | 9.24E-42 | 1.68E-37 |
| 15 | KLF6 | Tuft cells | 0.9506938 | 90.8% | 58.0% | 1.50E-32 | 2.73E-28 |
| 15 | CDC42EP5 | Enterocytes | 0.9305525 | 99.4% | 86.5% | 2.12E-33 | 3.86E-29 |
| 15 | CFB | Paneth cells | 0.90961 | 86.7% | 56.7% | 2.29E-24 | 4.17E-20 |
| 15 | SLC5A1 | Enterocytes | 0.8722516 | 54.9% | 27.2% | 1.18E-20 | 2.15E-16 |
| 15 | ELF3 | Enterocytes | 0.7545213 | 97.7% | 83.0% | 2.81E-19 | 5.11E-15 |
| 15 | GULP1 | Enterocytes | 0.7456127 | 43.9% | 17.0% | 1.47E-23 | 2.68E-19 |
| 15 | ABHD11 | Enterocytes | 0.7278028 | 60.1% | 22.0% | 3.69E-38 | 6.71E-34 |
| 15 | RELB | Microfold cells | 0.6983259 | 62.4% | 21.7% | 1.63E-44 | 2.96E-40 |
| 15 | CYBA | Microfold cells | 0.6900823 | 99.4% | 91.4% | 3.99E-19 | 7.25E-15 |
| 15 | VNN1 | Enterocytes | 0.6686265 | 54.3% | 29.6% | 6.20E-16 | 1.13E-11 |
| 15 | IRF6 | Microfold cells | 0.6629595 | 61.3% | 26.6% | 1.06E-26 | 1.92E-22 |
| 15 | SLC12A2 | Crypt cells | 0.6118334 | 58.4% | 36.4% | 3.01E-12 | 5.47E-08 |
| 15 | CLDN15 | Enterocytes | 0.5962495 | 37.6% | 22.3% | 1.28E-07 | 2.33E-03 |
| 15 | ALDOB | Enterocytes | 0.5530126 | 60.7% | 39.8% | 1.58E-10 | 2.88E-06 |
| 15 | KRT7 | Goblet cells | 0.5529729 | 97.7% | 90.5% | 1.77E-12 | 3.23E-08 |
| 15 | KLF5 | Enterocytes | 0.525437 | 81.5% | 59.4% | 1.45E-13 | 2.64E-09 |
| 15 | RIPK2 | Paneth cells | 0.5131565 | 43.9% | 15.8% | 4.52E-24 | 8.21E-20 |
| 15 | KLF3 | Tuft cells | 0.5096692 | 72.8% | 47.0% | 5.75E-15 | 1.05E-10 |
| 15 | FERMT1 | Enterocytes | 0.5092051 | 63.6% | 44.0% | 4.39E-09 | 7.98E-05 |
| 15 | TM4SF5 | Enterocytes | 0.4702122 | 48.6% | 31.2% | 1.97E-07 | 3.59E-03 |
| 15 | SOX9 | Tuft cells | 0.4654268 | 68.8% | 48.1% | 1.17E-09 | 2.12E-05 |
| 15 | GLS | Enterocytes | 0.4597126 | 48.6% | 23.6% | 5.59E-15 | 1.02E-10 |
| 15 | CTNNB1 | Paneth cells | 0.4514236 | 76.9% | 46.7% | 4.74E-18 | 8.63E-14 |
| 15 | MUC4 | Goblet cells | 0.4481927 | 46.2% | 26.0% | 1.23E-09 | 2.23E-05 |
| 15 | CDX1 | Enterocytes | 0.4406712 | 47.4% | 24.5% | 8.15E-13 | 1.48E-08 |
| 15 | CD24 | Paneth cells | 0.4129769 | 97.7% | 90.6% | 2.31E-15 | 4.20E-11 |
| 15 | BACE2 | Goblet cells | 0.3933818 | 86.1% | 63.3% | 4.16E-11 | 7.57E-07 |
| 15 | ALCAM | Enteroendocrine cells | 0.384968 | 40.5% | 21.9% | 2.29E-09 | 4.17E-05 |
| 15 | PRAP1 | Enterocytes | 0.3445322 | 79.2% | 69.8% | 2.24E-04 | 1.00E+00 |
| 15 | MIER3 | Microfold cells | 0.3434085 | 25.4% | 12.0% | 2.04E-08 | 3.70E-04 |
| 15 | TMEM45B | Enterocytes | 0.3372559 | 77.5% | 67.1% | 1.04E-05 | 1.89E-01 |
| 15 | PPP1R14D | Enterocytes | 0.3355069 | 74.6% | 59.2% | 1.06E-05 | 1.93E-01 |
| 15 | KCNE3 | Crypt cells | 0.3216387 | 45.7% | 36.3% | 7.44E-04 | 1.00E+00 |
| 15 | LYZ | Paneth cells | 0.3143866 | 99.4% | 96.4% | 1.45E-08 | 2.63E-04 |
| 15 | GJB2 | Microfold cells | 0.2785151 | 28.3% | 12.6% | 3.79E-10 | 6.89E-06 |
| 15 | ACSL5 | Enterocytes | 0.2730936 | 63.6% | 42.2% | 2.55E-08 | 4.63E-04 |
| 15 | FUOM | Enterocytes | 0.2500377 | 79.8% | 62.3% | 1.39E-06 | 2.54E-02 |
| 16 | SPINK4 | Paneth cells | 1.1023833 | 36.6% | 15.4% | 5.44E-08 | 9.89E-04 |
| 16 | SLC12A2 | Crypt cells | 1.018017 | 62.2% | 36.6% | 6.81E-09 | 1.24E-04 |
| 16 | KRT7 | Goblet cells | 0.91287 | 100.0% | 90.6% | 1.93E-14 | 3.51E-10 |
| 16 | G0S2 | Enterocytes | 0.779919 | 35.4% | 15.2% | 1.91E-07 | 3.47E-03 |
| 16 | KLF3 | Tuft cells | 0.7397934 | 86.6% | 47.1% | 5.38E-18 | 9.78E-14 |
| 16 | ELF3 | Enterocytes | 0.7356122 | 100.0% | 83.1% | 5.94E-18 | 1.08E-13 |
| 16 | MMP7 | Paneth cells | 0.7179741 | 100.0% | 76.6% | 3.62E-19 | 6.58E-15 |
| 16 | CFD | Paneth cells | 0.7138688 | 50.0% | 25.1% | 4.33E-09 | 7.87E-05 |
| 16 | CTNNB1 | Paneth cells | 0.6922666 | 79.3% | 46.9% | 1.51E-12 | 2.74E-08 |
| 16 | SOX9 | Tuft cells | 0.6891953 | 87.8% | 48.1% | 4.09E-16 | 7.44E-12 |
| 16 | PMEPA1 | Enterocytes | 0.6824712 | 89.0% | 54.8% | 4.47E-15 | 8.13E-11 |
| 16 | CTSD | Microfold cells | 0.6804099 | 98.8% | 84.6% | 5.87E-12 | 1.07E-07 |
| 16 | KCNE3 | Crypt cells | 0.6510025 | 62.2% | 36.2% | 1.88E-08 | 3.43E-04 |
| 16 | AHR | Microfold cells | 0.6146585 | 84.1% | 35.2% | 2.21E-20 | 4.02E-16 |
| 16 | COL18A1 | Enterocytes | 0.5819668 | 28.0% | 14.9% | 7.93E-05 | 1.00E+00 |
| 16 | IRF2 | Microfold cells | 0.5532893 | 65.9% | 28.7% | 2.22E-14 | 4.03E-10 |
| 16 | PRSS1 | Paneth cells | 0.5437518 | 37.8% | 19.0% | 7.48E-06 | 1.36E-01 |
| 16 | TRNP1 | Enteroendocrine cells | 0.5418598 | 80.5% | 53.5% | 3.67E-09 | 6.67E-05 |
| 16 | MUC4 | Goblet cells | 0.5353656 | 64.6% | 26.0% | 1.51E-14 | 2.74E-10 |
| 16 | KLF5 | Enterocytes | 0.5313557 | 98.8% | 59.4% | 9.82E-18 | 1.79E-13 |
| 16 | VNN1 | Enterocytes | 0.5204808 | 58.5% | 29.8% | 2.26E-09 | 4.11E-05 |
| 16 | PPARG | Enterocytes | 0.5045623 | 76.8% | 50.3% | 6.94E-10 | 1.26E-05 |
| 16 | KCNN4 | Paneth cells | 0.501225 | 58.5% | 27.0% | 1.44E-11 | 2.62E-07 |
| 16 | NFIB | Microfold cells | 0.4880876 | 56.1% | 23.3% | 1.55E-12 | 2.81E-08 |
| 16 | EHF | Enterocytes | 0.4623502 | 70.7% | 32.3% | 1.53E-13 | 2.78E-09 |
| 16 | ANXA5 | Microfold cells | 0.4540507 | 98.8% | 71.9% | 9.35E-12 | 1.70E-07 |
| 16 | NFIC | Microfold cells | 0.4328474 | 50.0% | 18.0% | 1.10E-13 | 2.00E-09 |
| 16 | ALCAM | Enteroendocrine cells | 0.4188159 | 58.5% | 21.9% | 4.75E-14 | 8.64E-10 |
| 16 | IL13RA1 | Tuft cells | 0.4108764 | 61.0% | 30.6% | 2.67E-09 | 4.85E-05 |
| 16 | LYZ | Paneth cells | 0.3942749 | 100.0% | 96.5% | 3.53E-11 | 6.42E-07 |
| 16 | PLA2G2A | Paneth cells | 0.3924946 | 47.6% | 14.9% | 1.96E-16 | 3.57E-12 |
| 16 | AGR2 | Paneth cells | 0.3304038 | 100.0% | 90.7% | 8.28E-05 | 1.00E+00 |
| 16 | CDH17 | Enterocytes | 0.3127759 | 76.8% | 58.7% | 6.61E-05 | 1.00E+00 |
| 16 | FERMT1 | Enterocytes | 0.3113866 | 68.3% | 44.2% | 8.70E-06 | 1.58E-01 |
| 16 | TMC4 | Enterocytes | 0.3093676 | 70.7% | 48.3% | 2.92E-05 | 5.30E-01 |
| 16 | PDHA1 | Enterocytes | 0.3070068 | 53.7% | 27.2% | 2.43E-07 | 4.41E-03 |
| 16 | CCK | Enteroendocrine cells | 0.3061564 | 50.0% | 27.3% | 8.15E-06 | 1.48E-01 |
| 16 | CD24 | Paneth cells | 0.301182 | 100.0% | 90.6% | 4.98E-05 | 9.05E-01 |
| 16 | EPHB2 | Crypt cells | 0.2886267 | 30.5% | 10.2% | 2.00E-09 | 3.64E-05 |
| 16 | ALOX5 | Tuft cells | 0.2852735 | 47.6% | 25.9% | 6.54E-06 | 1.19E-01 |
| 16 | GALNT12 | Goblet cells | 0.2807799 | 46.3% | 20.6% | 5.49E-08 | 9.98E-04 |
| 16 | CXCL16 | Microfold cells | 0.2742985 | 76.8% | 53.2% | 7.74E-06 | 1.41E-01 |
| 16 | KLF6 | Tuft cells | 0.2659476 | 89.0% | 58.3% | 1.92E-08 | 3.49E-04 |
| 16 | TULP4 | Microfold cells | 0.2658237 | 37.8% | 20.0% | 9.03E-05 | 1.00E+00 |
| 16 | DPP4 | Enterocytes | 0.2623695 | 28.0% | 11.2% | 1.52E-06 | 2.77E-02 |
| 16 | CTSH | Microfold cells | 0.261104 | 58.5% | 40.5% | 4.64E-04 | 1.00E+00 |
| 16 | ATG16L1 | Paneth cells | 0.254944 | 31.7% | 14.2% | 4.28E-06 | 7.78E-02 |
| 16 | LGR4 | Paneth cells | 0.2531563 | 56.1% | 28.1% | 2.36E-07 | 4.29E-03 |
| 16 | HNF4A | Enterocytes | 0.2512606 | 46.3% | 27.0% | 5.13E-05 | 9.33E-01 |

Supplementary material S8. Number and percentage of epithelial cell types occupying in the Ctrl and CD enteroids treated with and without IFN-γ.

|  | Number | % of total analyzed cells | **Ctrl enteroids** | | | | | **CD enteroids** | | | | |
| --- | --- | --- | --- | --- | --- | --- | --- | --- | --- | --- | --- | --- |
|  |  |  | IFN-γ 0 pg/mL | | IFN-γ 100 pg/mL | | *p*-value | IFN-γ 0 pg/mL | | IFN-γ 100 pg/mL | | *p*-value |
|  |  |  | Number | % of cells in IFN-γ-free Ctrl enteroids | Number | % of cells in IFN-γ-treated Ctrl enteroids |  | Number | % of cells in IFN-γ-free CD enteroids | Number | % of cells in IFN-γ-treated CD enteroids |  |
| Intestinal stem cells | 1,434 | 13% | 487 | 11% | 318 | 20% | < 0.0001 | 322 | 11% | 307 | 13% | 0.0147 |
| Proliferating cells | 353 | 3% | 227 | 5% | 14 | 1% | < 0.0001 | 98 | 3% | 14 | 1% | < 0.0001 |
| Enterocytes | 1,060 | 9% | 302 | 7% | 66 | 4% | 0.0004 | 602 | 21% | 90 | 4% | < 0.0001 |
| Goblet cells | 1,581 | 14% | 461 | 10% | 452 | 29% | < 0.0001 | 287 | 10% | 381 | 17% | < 0.0001 |
| Paneth cells | 707 | 6% | 296 | 6% | 74 | 5% | 0.0116 | 220 | 8% | 117 | 5% | 0.0003 |
| Unclassified | 6,181 | 55% | 2,781 | 61% | 635 | 41% | < 0.0001 | 1,371 | 47% | 1,394 | 61% | < 0.0001 |
| Total | 11,316 | 100% | 4,554 | 100% | 1,559 | 100% |  | 2,900 | 100% | 2,303 | 100% |  |

Supplementary material S9. Determination of therapeutic concentration of upadacitinib. Bright field microscopic image of human enteroids treated with IFN-γ 100 pg/mL and different concentrations of upadacitinib.

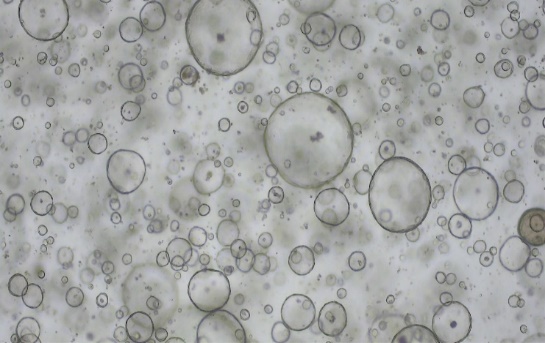

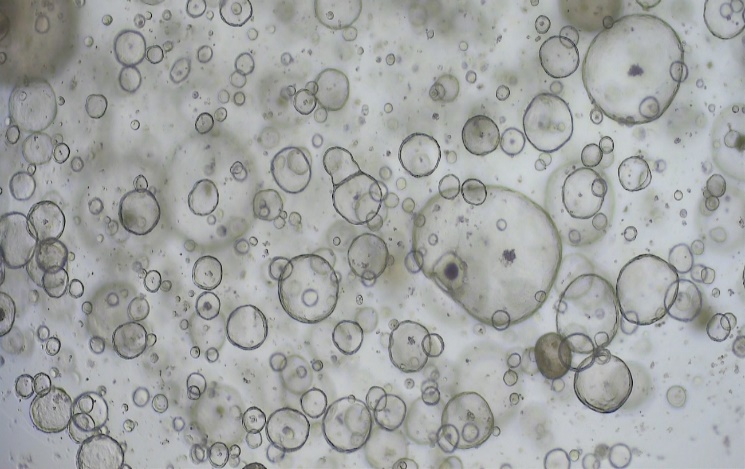

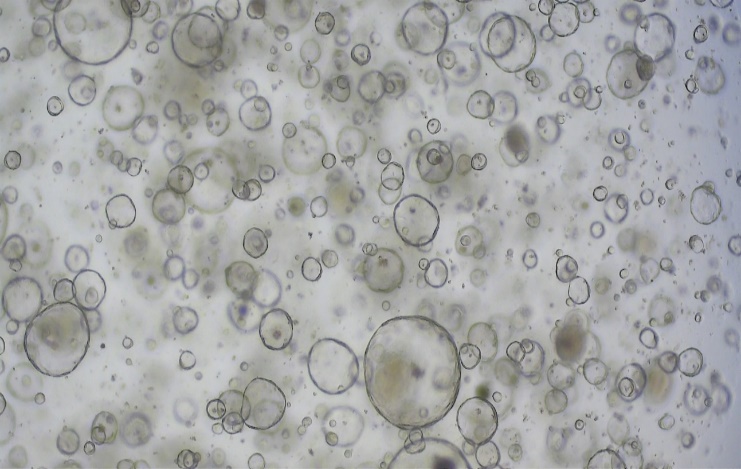

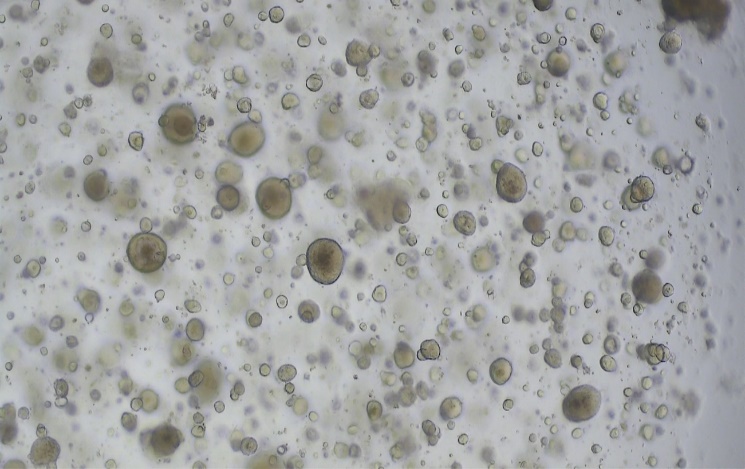

1. Control entroids

2. IFN-γ 100 pg/ml

3. IFN-γ 100 pg/ml + upa-dacitinib 0.35 μg/ml ≈ 1 μM

4. IFN-γ 100 pg/ml + upa-dacitinib 3.5 μg/ ml ≈ 10 μM

5. IFN-γ 100 pg/ml + upa-dacitinib 35 μg/ ml ≈ 100 μM

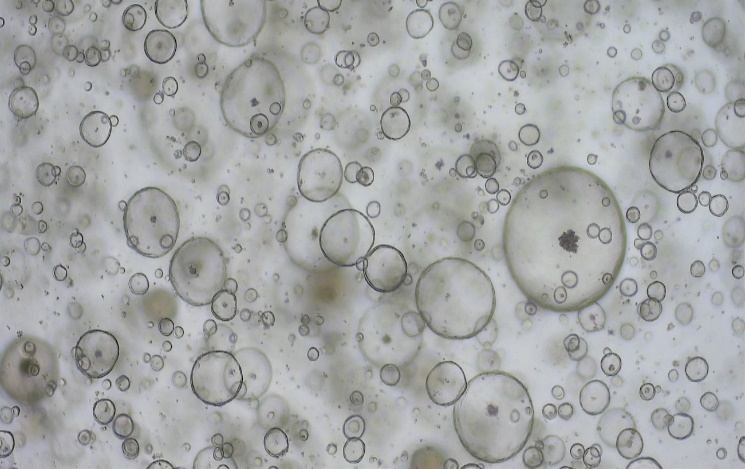

Supplementary material S10. mTOR and autophagy-related gene expression in Ctrl and CD treated with and without IFN-γ.

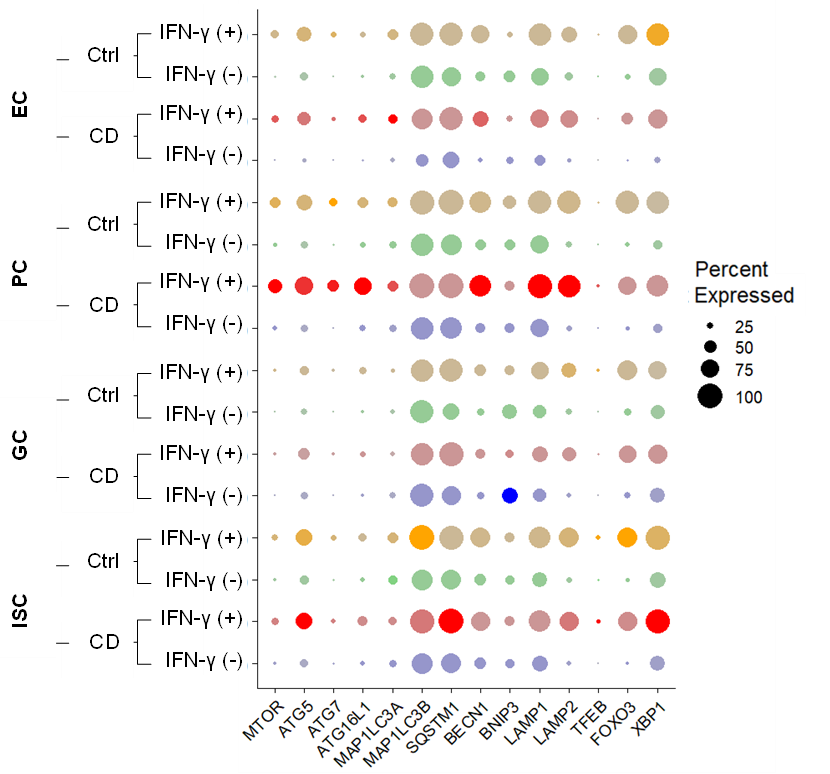
